# Identifiability of Bayesian Models of Cognition

**DOI:** 10.1101/2025.06.25.661321

**Authors:** Michael Hahn, Entang Wang, Xue-Xin Wei

**Affiliations:** Saarland University, Saarbrücken, Germany; Department of Neuroscience, Department of Psychology, Center for Perceptual Systems, Center for Learning and Memory, Center for Theoretical and Computational Neuroscience, The University of Texas at Austin

## Abstract

Inferring the underlying computational processes from behavioral measurements is a fundamental approach in cognitive science and neuroscience. Although Bayesian decision theory has become a major normative framework for modeling cognition, it is unclear to what extent its modeling components (i.e., prior belief, likelihood function, and the loss function) can be recovered from behavioral data. Here, we systematically investigated the problem of inferring such Bayesian models from behavioral tasks. In contrast to a pessimistic picture often painted in previous research, our analytical results guarantee in-principle identifiability under broadly applicable conditions, without any *a priori* knowledge of prior or encoding. Simulations and applications on the basis of behavioral datasets validate that the predictions of this theory apply in realistic settings. Importantly, our results demonstrate that reliable recovery of the model often requires having data from multiple noise levels. This is a crucial insight that will guide future experimental design.

## 1 Introduction

In cognitive science and neuroscience, Bayesian decision theory has emerged as a major principled framework for understanding human perception (1, 2, 3, 4, 5, 6) and cognition (7, 8, 9). This framework provides a normative recipe for modeling behavior by combining three ingredients (10). The first is the prior belief, which is a probability distribution that describes which stimuli the observer *a priori* considers more or less likely to have appeared, prior to any sensory input. The second is the likelihood function, which describes how likely the sensory observation is to be generated based on a given stimulus. The prior belief and the likelihood function are combined via Bayes’ rule to obtain a posterior probability distribution, i.e., an estimate of how likely each possible stimulus value is to have produced the noisy encoding. The third ingredient is the loss function, an assignment of costs to estimation errors. It may, for instance, be relatively tolerant of small estimation errors, or could penalize all estimation errors equally. Bayesian decision theory prescribes choosing the stimulus that minimizes the posterior expectation of the loss function.

There has been substantial interests in reverse-engineering these components from measured human behavior (11, 12, 13, 14, 15). This approach has been applied to the domains of perception (11, 12, 15), sensory-motor processing (16, 13) and, more recently, in neuroeconomics (17, 18). The basic idea is to make inference about the prior, encoding and loss functions from behavioral observations. While Bayesian models have been widely used in behavioral sciences, surprisingly, the question of whether Bayesian models are identifiable has not been systematically investigated. It is sometimes believed that the Bayesian approach is underconstrained and inherently degenerate (15), in that multiple different models can give rise to the same behavior. Prior studies have often presupposed some part of the model, as a way of constraining the model fit. As noted by (19), work often fixes one of the three components when estimating observer models, e.g. constraining the encoding by measuring discriminability (e.g. 20), measuring naturalistic priors (e.g. 21), or imposing a loss function experimentally (e.g. 22). So far, it remains unknown whether the model components of Bayesian models are actually identifiable from behavioral data. Perhaps not surprisingly, the lack of a theoretical foundation in model recovery has led to serious criticism of the Bayesian approach (23, 24).

Understanding whether a computational approach can recover the ground truth from empirical data is a basic problem in virtually all areas of science that use model-based inference, not limited to Bayesian models. In statistical estimation, this problem can be formulated as the consistency of the estimator (25, 26), and is fairly well understood for simple estimation problems, *e*.*g*., estimating the mean of a Gaussian random variable from samples. However, for more complex models, it can be challenging to precisely understand the conditions under which the models are identifiable (27). The question of model identifiablity has drawn more attention in economics (28, 29) and systems biology (30, 31). However, these results are not directly applicable for Bayesian models of cognition. The lack of understanding of identifiablity of Bayesian models poses a fundamental challenge for interpreting empirical observations from experiments.

We fill this critical gap by developing a rigorous theory of the identifiability of Bayesian models of perceptual decision making from behavioral data. On the one hand, we find that, indeed, Bayesian models have substantial degeneracy, in that simple experimental setups will, in systematic ways, fail to fully identify the three model components. On the other hand, we rigorously prove that, using more carefully designed experimental setups, e.g., by experimentally varying the magnitude of neural noise between trials, one can systematically eliminate this degeneracy. Remarkably, we find that Bayesian models are identifiable without any of the parametric assumptions often made for the encoding or the prior. A crucial insight from these results is that it is beneficial to collect measurements at multiple levels of sensory/neural noise in experiments in order to recover the model. Simulations and applications on the basis of a series of previously published behavioral datasets validate the theoretical results in realistic settings. Our results have important implications in interpreting results from fitting Bayesian observer models to empirical data and provide guiding principles for designing new experiments to understand how perceptual decisions are computed in the brain based on behavioral measurements. A preliminary version of this work was presented earlier (32).

## 2 Results

Following previous studies (33, 34, 5, 17, 6), we model the neural processing of a stimulus variable θ (for example, the orientation of a bar) as a cascade of encoding and decoding computations. In particular, we model the encoding of a one-dimensional variable using a general nonlineary transformation *F*, subject to additive noise (see Fig. 1):

**Figure 1:**
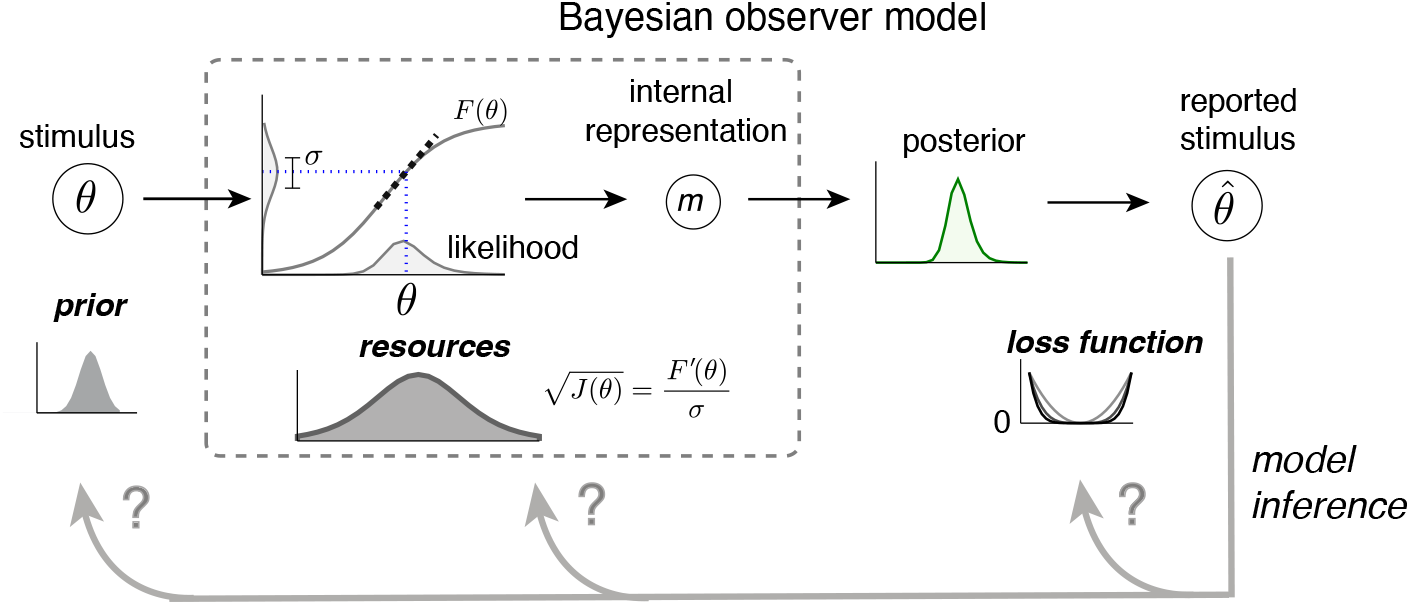
The Bayesian model framework and the problem of model identification. For a given stimulus variable θ, the neural code is modeled as a one-dimensional encoding function plus sensory noise. The sensitivity of the population activity to the stimulus is described by the Fisher information. Given the encoding function *f* and sensory (neural) noise, Bayesian inference is used to derive a point estimate 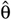. In continuous estimation tasks, observers reproduce 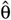 subject to motor noise. The identifiability problem asks to what extent it is possible to recover the three components of the Bayesian model – prior, encoding resources (Fisher information), and loss function – from the behavioral responses.

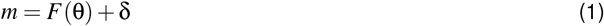

where δ represents the sensory noise, which is assumed to have a Gaussian distribution with variance σ^2^ describing the magnitude of sensory noise (internal noise). Sensory noise arises due to limitations in the number of spikes or cognitive resources. We focus on the 1-D case as most existing Bayesian models deal with a 1-D stimulus variable, but we expect the theory to extend to the multivariate case. The Bayesian framework contains several important components. The first component is the encoding function *F*. The second component is the prior, given by a probability distribution *p*_*prior*_(θ). The third component is the loss function, *𝓁*(*x, y*). Popular loss functions involve 0-1 loss, squared error loss, absolute distance, which are members of the general family of *L*_*p*_ loss functions. Under the Bayesian framework, the observer obtains a noisy sensory measurement on each trial and computes a posterior distribution by multiplying the prior and likelihood. The reported stimulus by the observer is modeled as the stimulus that minimizes the loss function.

We are interested in understanding whether the components of the Bayesian model can be recovered from behavioral responses. While our analytical results on the Bayesian models below focus on the small-noise regime, extensive numerical simulations are used to verify that conclusions hold more generally.

### 2.1 Prior and encoding can be identified when the loss function is known

Suppose that we have collected behavioral responses from a sufficiently large number of trials in an estimation task from a Bayesian observer. On each trial, a particular stimulus θ was presented, and the observer reported the stimulus that they perceived. We first consider the case that the loss function is known to be the squared error loss (a popular loss function assumed in the literature). We ask whether one can recover the prior and encoding from the behavior responses. Theoretically, we find that the answer is always “yes”, under mild assumptions on the smoothness of the resource allocation and the prior, and small sensory noise.

First, the resource allocation can be recovered by comparing the overlap between response distributions elicited by close-by stimuli; the smaller this overlap around a stimulus θ, the higher the resource allocation at θ. We show that this procedure can be used to exactly recover the resource allocation (Theorem 1).

Second, the prior distribution can be identified by further considering the relationship between the estimation bias and the slope of the prior, which largely follows a one-to-one correspondence under a squared error loss:

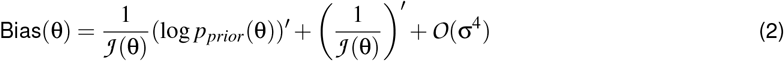

when 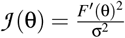 denotes the Fisher information, which represents the resource allocation of the encoding. Once the encoding has been identified, the prior can be identified from the bias via this identity.

Notably, the same conclusion holds when the known loss function belongs to the general *L*_*p*_ family, *i*.*e*., the observer’s loss is determined by a power function of the absolute error. This includes loss functions that are commonly adopted in the literature, e.g., MAP (*p →* 0), median (*p* =1), or mean squared error (*p* =2).

These analytical results are derived in the regime where sensory noise is small. To test their validity in realistic settings, we used a general fitting procedure that represents encoding and prior on a discretized grid, and optimizes these and the magnitude of noise to maximize trial-by-trial likelihood of observed data. Results based on extensive numerical simulations support our analytical results. We find that the model components are closely recovered when noise is small. Together, these results provide theoretical justifications of empirically reverse-engineering the prior and encoding when we have good knowledge of the observer’s decision rule.

Fig. 2 illustrates one such simulation example, inspired by data from orientation estimation tasks (35, 36, 6). In this particular simulation, the prior is assumed to be uniform and the encoding is such that cardinal orientations are encoded better. We found that, with about 1000 trials, the basic shape of the prior and encoding can be readily recovered. Notably, this suggests that one can leverage estimation tasks and the Bayesian framework to effectively infer the allocation of encoding resources. This observation may have important implications for experimental work. The coding resource is the inverse of the discrimination threshold, which is an important behavioral measure in psychophysics that is often estimated using binary classification tasks. One limitation of inferring discriminability using binary classification tasks is that it often requires a large number of trials. Our simulation results suggest that the continuous estimation paradigm offers an viable alternative in inferring stimulus-dependent coding resources for one-dimensional stimulus variables. Later we will test this procedure further using experimental data.

**Figure 2:**
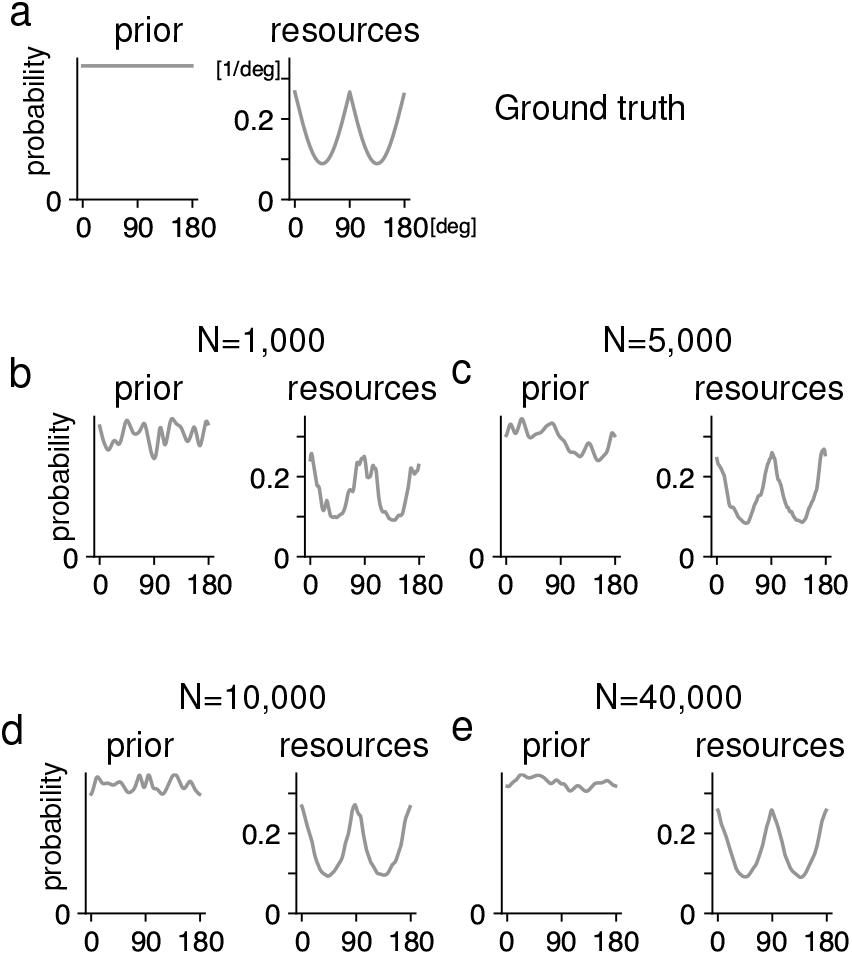
Prior and encoding can be identified when the loss function is known. Here we illustrate the identifiability of prior and encoding resources when the loss function is known using an example. In particular, we show that nonparametric priors and encodings with *L*_2_ loss can be identified when sufficiently many trials at small sensory noise are available. (a) The ground truth model. The prior is assumed to be uniform. The resource is assumed to be higher for the cardinal orientations (0 and 90 degee), and lower for the oblique orientations (45 and 135 degree). The loss function is the squared error loss, and is assumed to be known. (b) We simulated a dataset with 1000 trials by randomly drawing 1000 samples from the generative model in (a). We then fit the Bayesian model to this simulated dataset assuming a squared error loss function. The results show that inferred prior and resources capture the basic patterns of the ground truth. (c,d,e) Similar to (b), but for even larger sample sizes. Increasing the sample size leads to more accurate model recovery. SI Appendix, Figures S35–S36, has results for further models and other noise magnitudes.

When noise increases, the allocation of the encoding resource (i.e., encoding) can still be recovered, though the prior may become difficult to recover as large noise may wash out details of the prior (SI Appendix, Fig. S13). These results have a further implication for future experiments, that is, the inclusion of conditions at small sensory noise is important for identifying the prior. For vision experiments, this may be done by increasing the contrast or the presentation time of the visual stimuli.

### 2.2 Prior and loss function can be systematically confounded when the loss function is unknown

While, in some experiments, we may have *a priori* good knowledge about the loss function used by the observer, this does not hold generally. Thus, we next consider the question of model identifiability under a more general setting for which the loss function is unknown. For now, we will assume that the magnitude σ of sensory noise is small and shared across all trials, and that the loss function belongs to the general *L*_*p*_ loss family. Importantly, neither the magnitude of sensory noise nor the exponent *p* are known *a priori*.

Using the same theoretical arguments as above, the encoding (i.e., Fisher information) can be well identified even when the loss is unknown. However, for the loss function and prior, we find that they can be systematically confounded. In particular, when the sensory noise is small, two models are indistinguishable if and only if their encodings are identical, and their priors are linked as:

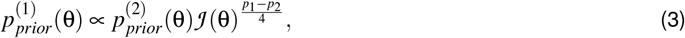

where the integers *p*_1_ and *p*_2_ are the exponents of the loss functions of the two encoding models (see SI Appendix, Theorem S9 for formal proof). The MAP estimator (*p →* 0 in the loss function) is special; here, the exponent needs to be replaced by *−* 1 rather than 0 in this formula. Thus, prior and loss function can be systematically confounded in experiments with only one level of small sensory noise.

These points are illustrated using an example in Fig. 3. We generate synthetic datasets based on the same ground truth model as in Fig. 2 : uniform prior, higher encoding precision for cardinal orientations, and squared error loss functions. We then fit the Bayesian model assuming the correct loss function. Fig. 3d shows that the fit can well recover the prior and the encoding model, as expected. However, because the loss function is not known in this problem, one may also fit a model assuming a different loss function, e.g., *L*_0_ (MAP estimator) or *L*_8_ loss. Fitting to the data with those functions, one finds that the encoding is again well recovered. However, the fitted prior in this case has substantial lower (*p* =0) or higher (*p* =4, 8) density at the cardinal orientations ((Fig. 3c,e,f). Which prior reflects the ground truth then? One may attempt to use the log-likelihood of the model fits to adjudicate between the models. However, models with different loss functions explain the data equally well (Fig. 3b). Thus, in this case, it is impossible to identify the model.

**Figure 3:**
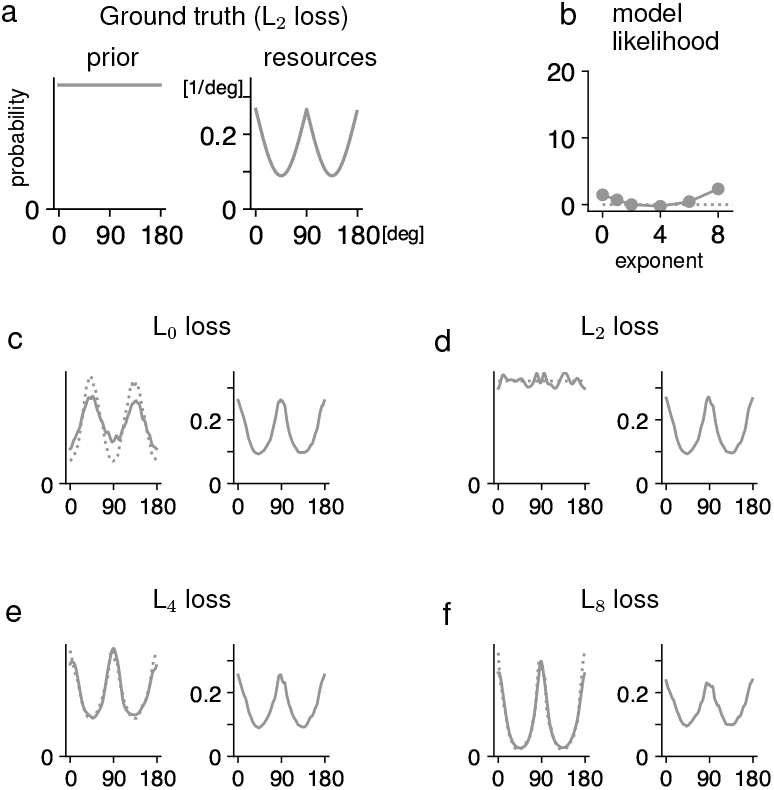
Prior and loss function can be confounded when both are unknown. We use a simulation example to demonstrate that simple experimental designs are typically sufficient for identifying the encoding, but cannot recover prior and loss function. (a) The ground truth model component. (b) We generate a dataset with 10K trials data assuming the model in (a). We then fit the data to models assuming different loss functions parameterized by a *L*_*p*_ family of functions. Model fit (heldout negative log likelihood, NLL) does not single out the ground-truth exponent, making it difficult to adjudicate between different models. (c,d,e,f) The best fitted prior and encoding resources corresponding different loss functions. In each case, we plot both an idealized curve predicted by the theory (dotted, Equation 3), and a fit generated by the fitting procedure (solid). We observe that, while the encoding is recovered throughout, the prior is recovered incorrectly (in accordance with Equation 3) when the loss function is misspecified. Overall, this means that the prior and loss function are confounded.

While Eq. (3) shows that the prior and loss functions generally cannot be identified when the loss is unknown, there is a special case where the prior can be identified: when the Fisher information is uniform. In this case, according to Eq. (3), the fitted prior is identical under different *L*_*p*_ loss functions, thus can be identified, even though the loss function may remain unidentifiable.

### 2.3 Leveraging behavioral responses based on multiple sensory noise levels makes most models identifiable

At first sight, the results described above may suggest a somewhat pessimistic view of the ability to identify Bayesian observer models when neither the prior nor the loss function is known *a priori*. However, it is important to realize that the setting studied above assumes that one only collects the behavioral responses of the observer under a single sensory noise condition, i.e., the magnitude of the sensory noise is shared in all trials. Crucially, we find that this degeneracy in identifiability can typically be avoided by experimental setups where the amount of sensory noise is not shared across all trials. In vision psychophysics experiments, sensory noise may be effectively manipulated by varying the length of the stimulus presentation time or the contrast of the stimulus.

Importantly, our theoretical analysis shows that the components of Bayesian observer models are usually identifiable if response distributions based on multiple small sensory noise levels and a sufficiently large number of trials are available (Theorem 3 in Materials and Methods). At a more technical level, the reason enabling this identifiability with multiple sensory noise levels lies in the second-order, nonlinear scaling of the bias with the magnitude of the sensory noise. Different combinations of prior and loss function often make different predictions regarding how the biases should change with the sensory noise levels, which enables ruling out non-ground-truth loss functions (SI Appendix, Fig. S12).

To validate these theoretical results numerically, we generate many random models with smooth encoding functions and smooth prior density functions, together with the exponent *p* of the *L*_*p*_ loss function. For each model, we generate synthetic datasets by sampling behavioral responses based on multiple noise levels, and then investigate whether the model components can be recovered from the simulated data. In these numerical simulations with experimentally relevant sample sizes, we consistently find that the model components (i.e., prior, encoding, and exponent of the loss function) can be reliably identified when behavioral responses from multiple noise levels are available. Fig. 4 shows three examples of randomly generated models. See Section S4.4 of the Appendix for further examples across loss functions.

**Figure 4:**
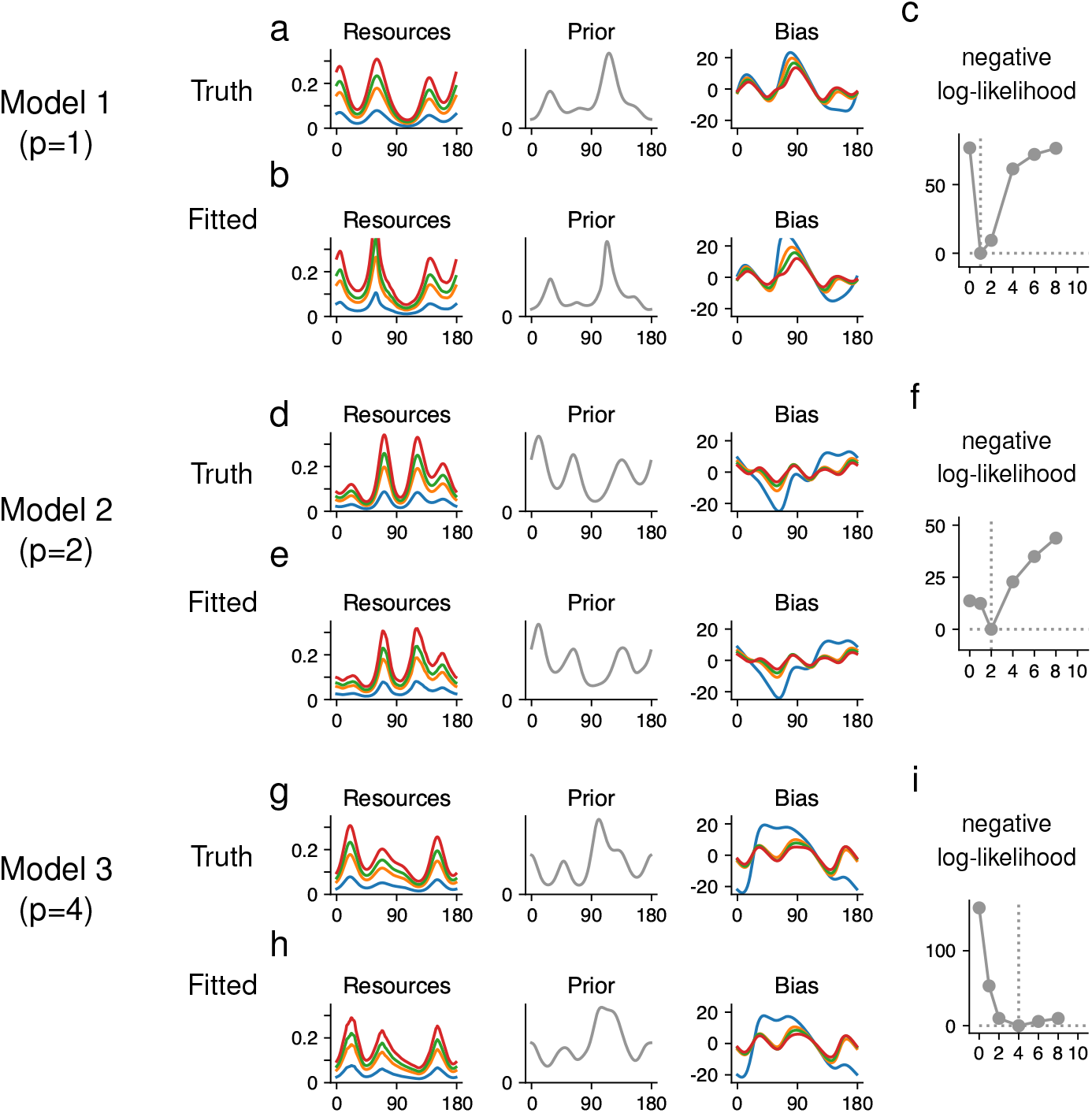
Simulation examples showing that random models can be identified with data from multiple noise levels. Here we show model recovery results from three randomly sampled models, on simulated datasets with 10K trials each. The sensory noise magnitudes are derived from the dataset collected by (35), but resources and prior are randomly generated. (a) The encoding resources, prior distribution of the ground truth model used in the simulation, as well as the predicted behavioral biases of the Bayesian observer. The ground truth loss function is *L*_1_ for this model. (b) The inferred resources, inferred prior and biases of the best-fit model. (c) The negative log-likelihood corresponding to the best-fit model under *L*_*p*_ loss functions with different exponent. (d-f) Similar to (a-c), but for a model with *L*_2_ loss function. (g-i) Similar to (a-c), but for a model with *L*_4_ loss function.

Psychophysical experiments are often limited by the number of trials. In light of our results above, an important question is, when the total number of trials is constrained, whether collecting behavioral responses at multiple noise levels (e.g., using visual stimuli with different contrast levels) would benefit the identifiability of models. To address this question, we systematically compared the ability to recover the model with a fixed total number of trials, while allowing the number of noise levels to vary in simulations. We perform this analysis across four different combinations of uniform or periodic priors and encodings (see Materials and Methods), across combinations of 1, 2, or 4 noise levels, and different numbers of trials, from 2,000 to 40,000.

These results are summarized in Fig. 5. We find that, for a given number of trials, increasing the number of noise levels is generally effective in removing the degeneracy found above. At one level of noise, the identifiability of the loss functions is variable: In some cases, particularly when *p* is small, a single noise level at finite noise already provides sufficient signal for constraining or even identifying *p*. This suggests that the response distribution obtained from finite noise may contain extra information of the loss functions compared to that of small noise. However, the data likelihoods across different exponents of the loss function are typically similar with experimentally realistic sample sizes, and the number of trials needed to faithfully identify the model with one noise level is not feasible in experiments. When including 2 levels of noise, identifiability is overall increased, in particular when high and low levels of noise are combined (SI Appendix, Fig. S14). Furthermore, we found that increasing the number of noise levels further beyond 2 was highly effective in further improving identifiability. With 4 levels of sensory noise, the negative log-likelihood (NLL) is steep and generally singles out the ground-truth *p*, or exponents close to it. Fig. 5d suggests that increasing the number of noise levels from one to four may be better for identifiability than increasing the number of trials by a factor of two or more. See SI Appendix S4.2 and S4.3 for more simulation results. An important consequence of these results for experimental design is that increasing the number of noise levels can be substantially more efficient for identifiability than increasing the number of trials.

**Figure 5:**
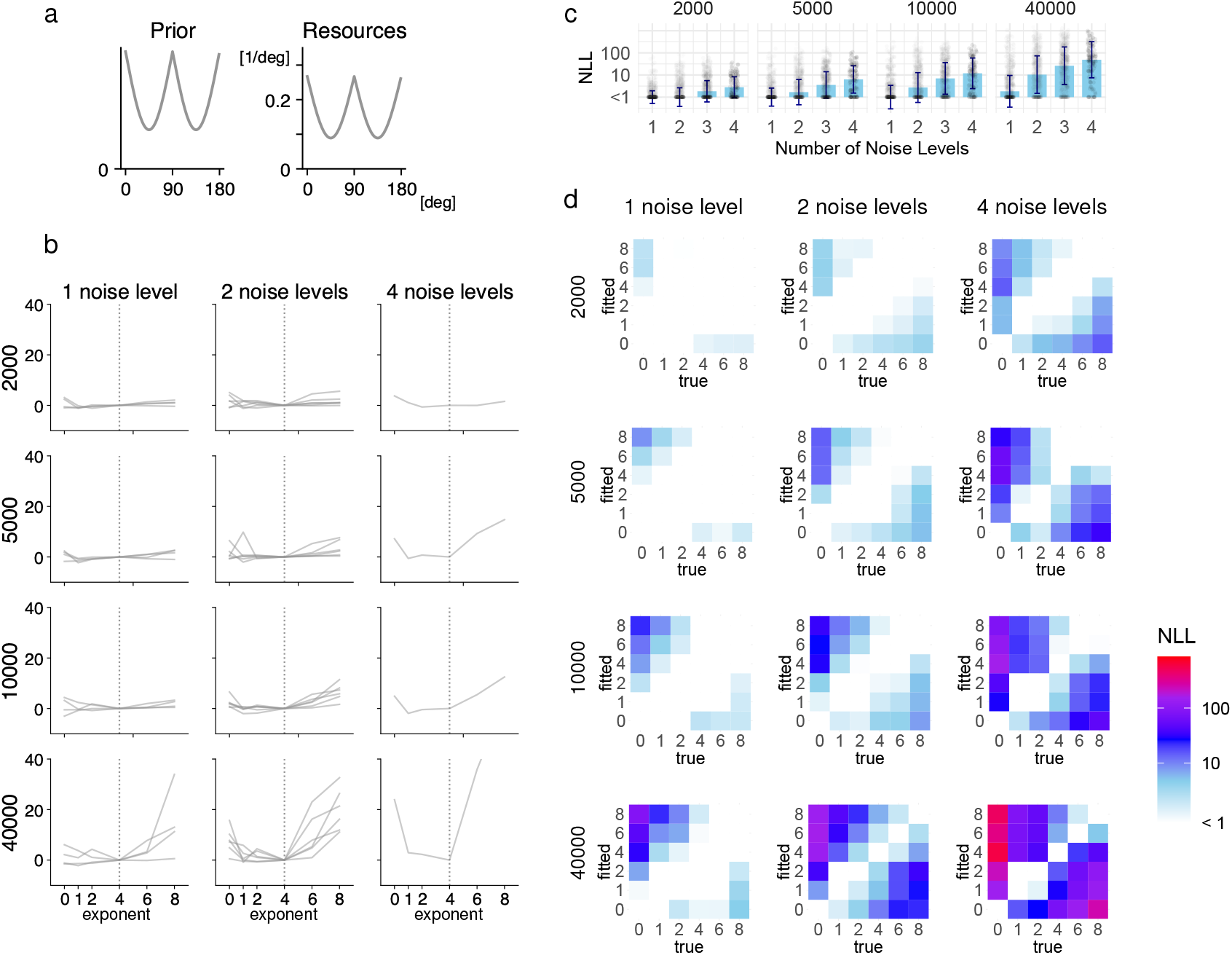
Example simulations showing the benefit of having behavioral responses from multiple noise levels for identifiability. (a) We simulated data with periodic prior and encoding at *p* =4. (b) Distinguishing loss functions, with an increasing number of sensory noise levels (columns: 1, 2, 4 levels) and an increasing number of trials (rows: 2K, 5K, 10K, 40K trials). Each line represents one synthetic dataset. The *y*-axis indicates model fit as quantified by heldout negative log-likelihood (NLL) relative to the ground-truth loss. With data from a single level of noise, even a dataset with 40K trials is not always sufficient for identifying the loss. In contrast, and in accordance with the theory, low and high exponents can be reliably distinguished using data from sufficiently many levels of sensory noise. (c) Summary across loss exponents. We repeated this experiment for a total of four synthetic combinations of prior and encoding (see Materials and Methods), for all exponents *p* =0, 1, 2, 4, 6, 8. Each dot indicates a combination of model, ground-truth exponent *p*, and non-ground-truth exponent *p*′ ≠ *p* used for fitting. Bars denote medians; error bars indicate SDs (on logarithmic scale). Despite substantial variability, increasing the number of noise levels tends to increase identifiability. (d) For each pair of ground-truth exponent *p* and exponent used in fitting *p*′, we computed the increase in NLL when fitting with *p*′ as compared to *p*, for four different synthetic combinations of prior and encoding (see Materials and Methods), with varying numbers of trials and noise levels. Heatmaps plot the median across model runs. High NLL increase off the diagonal indicates better identifiability of the loss function. Increasing the number of noise levels is highly effective in increasing identifiability.

### 2.4 Confronting models that are difficult to identify via adaption to a different stimulus statistics

We caution that there still exist models for which even having behavioral responses corresponding to multiple noise levels does not ensure identifiability. However, in a mathematical sense, such models are rare and exceptional. We theoretically analyze the probability of encountering such models in the space of all models. Under natural Gaussian-process-based notions of measure or volume on the space of models, the degenerate models have measure 0 (see Theorem 3, Materials and Methods). Note that this theoretical result is consistent with our numerical simulations above, which show that most randomly sampled models can be well identified.

While the identifiability guarantee is valid for all models except a set of measure zero, this exceptional set may nonetheless include models of real-world relevance. In particular, for commonly considered models for scalar variables where the encoding is based on Weber’s law (37, 38, 39) (i.e., the discrimination threshold increases in proportion to the stimulus magnitude), and the prior is lognormal (i.e., normal in the sensory space), an analytical calculation shows that this model belongs to the exceptional set. Interestingly, for this model, the loss function and prior are entangled regardless of the number of noise levels based on which the behavioral responses are collected (see SI Appendix, Section S1.1).

These results pose a challenge to experimental work, i.e., how might one still recover the loss function and the prior in this situation? Mathematically, the exceptional set consists of models where the encoding and the prior are aligned in a very specific way. We find that this ambiguity may be resolved by adapting the system to different contexts with different stimulus statistics to induce a change of the encoding or the prior. As the exceptional set has measure zero, almost all short-term stimulus priors would cause the overall model to exit the exceptional set. This holds when the prior and encoding adapt to the short-term stimulus statistics simultaneously, or when only one of the two adapts (see SI Appendix S2.7 for a detailed discussion).

### 2.5 Commonly considered motor noise in estimation tasks does not affect the identifiability

Estimation tasks typically involve the subject reproducing the stimulus variable, e.g., by adjusting a knob to report the perceived orientation of the stimulus. In such reproduction paradigms, motor noise, although generally small in many tasks, may contribute to the behavior response and may need to be explicitly modeled. Empirically, motor noise in these tasks is often modeled as Gaussian additive noise. We seek to understand whether motor noise will hinder the ability to identify the model. We find that adding Gaussian additive motor noise, more generally, symmetric additive noise, does not impact our result (see Theorem 4 in Materials and Methods): Sensory and motor sources of noise can be dissociated when a sufficient number of trials are available.

### 2.6 Applications to experimental datasets

We test our theory using multiple published datasets to demonstrate that the insights from our theory hold in practice. As we will show below, the results from analyzing the experimental data provide strong support for our theory. These results also suggest new experiments for future work.

#### Application 1: Inferring the stimulus-dependent neural resources from orientation estimation data

We first apply our theory to study a dataset collected by de Gardelle et al (35), which measured behavioral responses of human observers in an orientation-reproduction paradigm. In this experiment, the presentation time of the stimulus was systematically manipulated (20ms, 40ms, 80ms, 160ms, 1000ms; 5 conditions in total). We hypothesize that reducing the presentation time of the stimulus would increase the magnitude of the internal noise, and reduce the Fisher information at each stimulus orientation proportionally.

We start by estimating the FI from the behavioral responses by fitting the data from individual experimental conditions. Fig. 6a plots the recovered FI for each of the five conditions. As expected, the recovered FI are higher for the cardinal orientations than the oblique orientations. This result holds for each condition. These results are consistent with the previously reported oblique effect (40), that is, the discrimination threshold for the oblique orientations are higher than that of the cardinal orientations.

**Figure 6:**
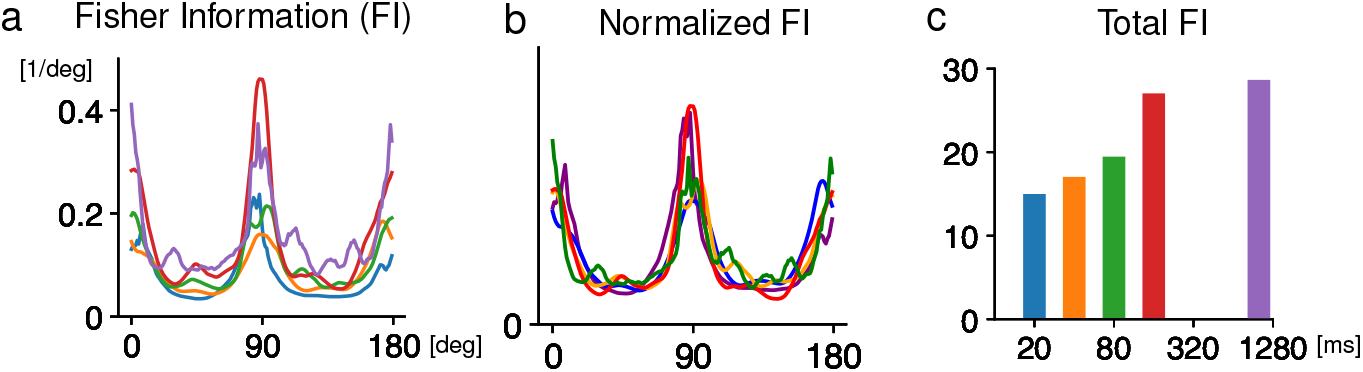
Application to experimental data: inferred the coding resource allocation for visual orientation from behavioral data. We seek to identify the encoding for orientation of Gabor arrays from human estimation data (collected by (35)). We report the results of fitting the encoding individually on 1K trials from each of the five exposure durations. (a) The square root of the Fisher information, 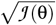 for each of the five experimental conditions. Each condition has a different amount of stimulus presentation time (i.e., 20ms, 40ms, 80ms, 160ms, 1000ms). (b) The normalized Fisher information, corresponding to the slope of the transfer function *F*′(θ), shows high consistency across the five conditions. (c) The total information 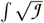 for the five conditions.

To test if the recovered allocation of coding resources is consistent across conditions, we normalized the square root of FI by the total information under each condition. As shown in Fig. 6b, the normalized information curves as a function of orientation are generally consistent across the conditions. These results, together with the results reported above based on simulations, suggest that the FI can be reliably identified from continuous estimation tasks. This provides a potentially more sample-efficient strategy to measure the precision of the neural representation for a continuous stimulus variable than the binary-choice paradigms.

To study how the information in the orientation code varies with the duration of the stimulus presentation, we quantify the total FI for each condition. We find that overall the integration of visual information over time in this task is sublinear (Fig. 6c), i.e., the total FI is substantially lower than that predicted from perfect integration of information over time. Interestingly, the largest increase of information happens between 80ms and 160ms. When comparing the total FI between the 160ms and 1000ms conditions, the FI for the latter only shows a mild increase.

#### Application 2: Identification of the prior and loss function from orientation estimation data

Using the same dataset from orientation estimation, we next investigate whether it would be possible to identify the loss function based on data from one noise level (operationally defined by the stimulus presentation time). We fit the models to data collected from individual conditions and examine the difference in the NLL for models with different loss functions. We find that, even with 2000 trials, the difference between the best-fitting exponent and worst-fitting exponent of the *L*_*p*_ loss function is generally smaller than 5. These results show that it was difficult to identify the loss function from the data collected for a single noise condition (Fig. 7b) for this task. It follows that the prior distribution is also difficult to identify (SI Appendix, Fig. S33).

**Figure 7:**
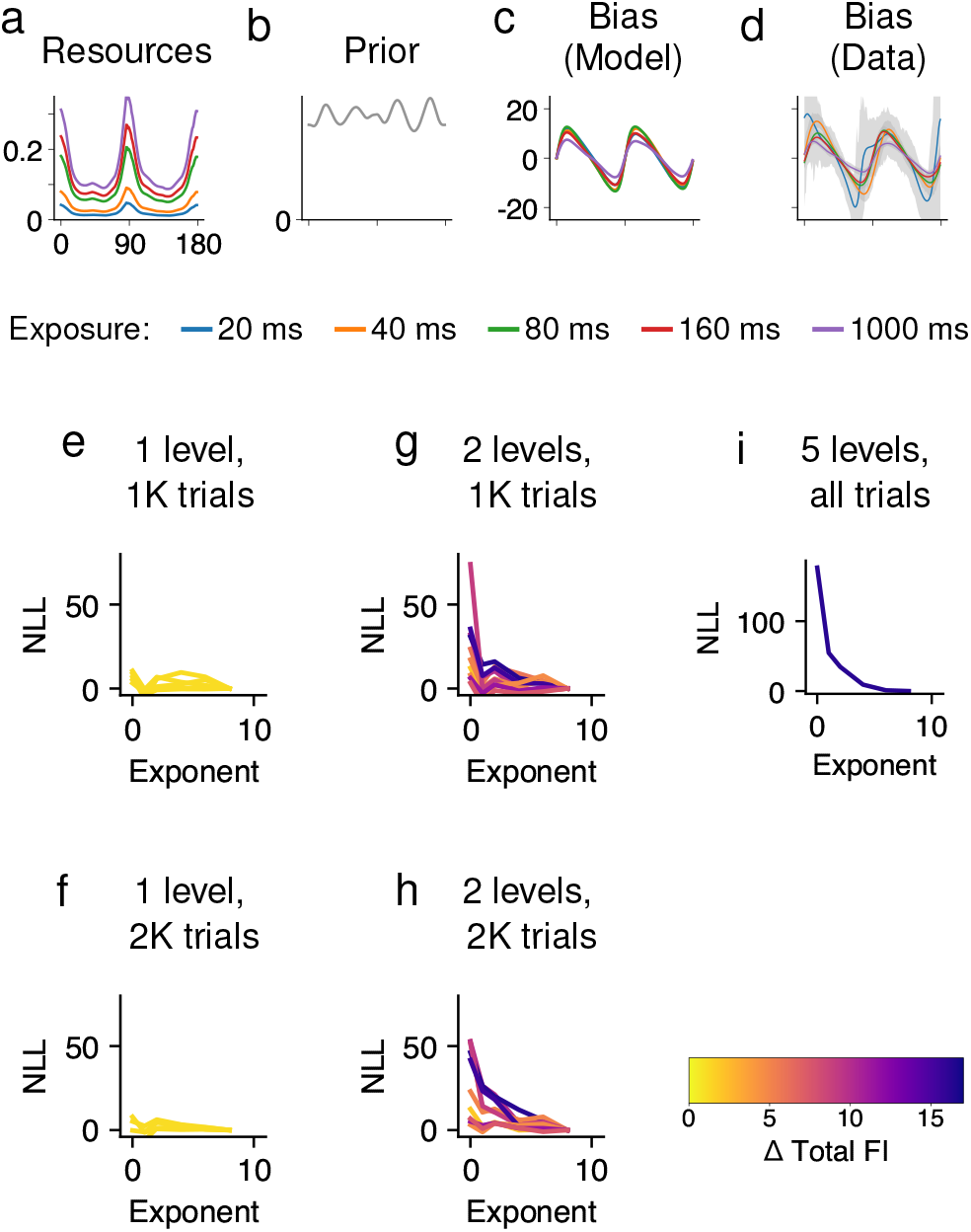
Application to experimental data: recovering the prior and loss function from human estimates of visual orientation. Model identification from human estimation responses for orientation of Gabor arrays (collected by (35)). (a,b): Neural resources and the prior inferred from the whole dataset that included data from 5 levels of sensory noise. Fitting recovers periodic resources and a close-to-uniform prior. (c,d) The best-fit model explains the biases observed in the data. (e-h) Comparing on subsampled datasets shows that the loss function is hard to identify when only a single noise level is available (yellow color), but become easier to identify when two noise levels are combined, at least when these correspond to noise levels with substantially different total FI (dark color). When evaluated on the whole data (9,936 trials), model fit favors higher loss function exponents (e.g., *p* =8).

According to our theory, we expect that the identifiablity of the model would increase when combining data from multiple conditions. To test this key theoretical prediction, we fit the model to the combined data from two noise levels together (a total of 10 different combinations). In general, we found that the loss function becomes more identifiable (Fig. 7b), at least when combining data from noise levels with very different total FI. With 2000 total trials, the difference between the best-fitting exponent and worst-fitting exponent of the *L*_*p*_ loss function can be larger than 20. This difference is much larger than the value obtained from fitting data from a single noise level using the same number of trials. Furthermore, using all data (10k trials) from all 5 conditions combined, we find the results clearly favor a large exponent in the *L*_*p*_ loss function (6), different from the popular squared error loss function considered in prior studies. Finally, the results from combining all data are consistent with the general patterns from combining two noise levels, further supporting that two noise levels might be in principle sufficient for identifiability.

Together, these empirical results support our theory, and suggest that it is beneficial to collect behavioral data under multiple noise levels experimentally.

#### Application 3: Difficulty in model identification from time interval data

We further examined a dataset from Remington et al. (38). In each trial of this experiment, a time interval of a certain duration was presented to the subject, who then reported the perceived time interval. The time intervals used were sampled from a bounded range. This task is typical in the study of perception of scalar variables. Similar paradigms were used in other studies (37, 41, 42). Bayesian models have been developed for this task, often under the assumptions that the stimulus encoding satisfied Weber’s law and the prior is Gaussian, log-normal, or uniform in the stimulus range. We previously found that Bayesian model with a log-normal prior outperforms models assuming Gaussian or uniform priors (6). As our theoretical analysis above suggests that this particular class of models may suffer from the issue of non-identifiability, we seek to test this prediction empirically.

Fitting the model to the data collected in one condition, we find that the data are well described by a unimodal prior resembling a lognormal prior (Fig. 8a-f). However, the loss function is difficult to identify. The NLL corresponding to models with different combinations of prior and loss functions are virtually identical based on these data (Fig. 8g). These results suggest that while the experimental data are well explained by Bayesian inference based on a log-normal prior operated on a neural representation consistent with Weber’s law, which is consistent with previous results (6), the precise parameter for the log-normal prior and the loss function cannot be identified from these data.

**Figure 8:**
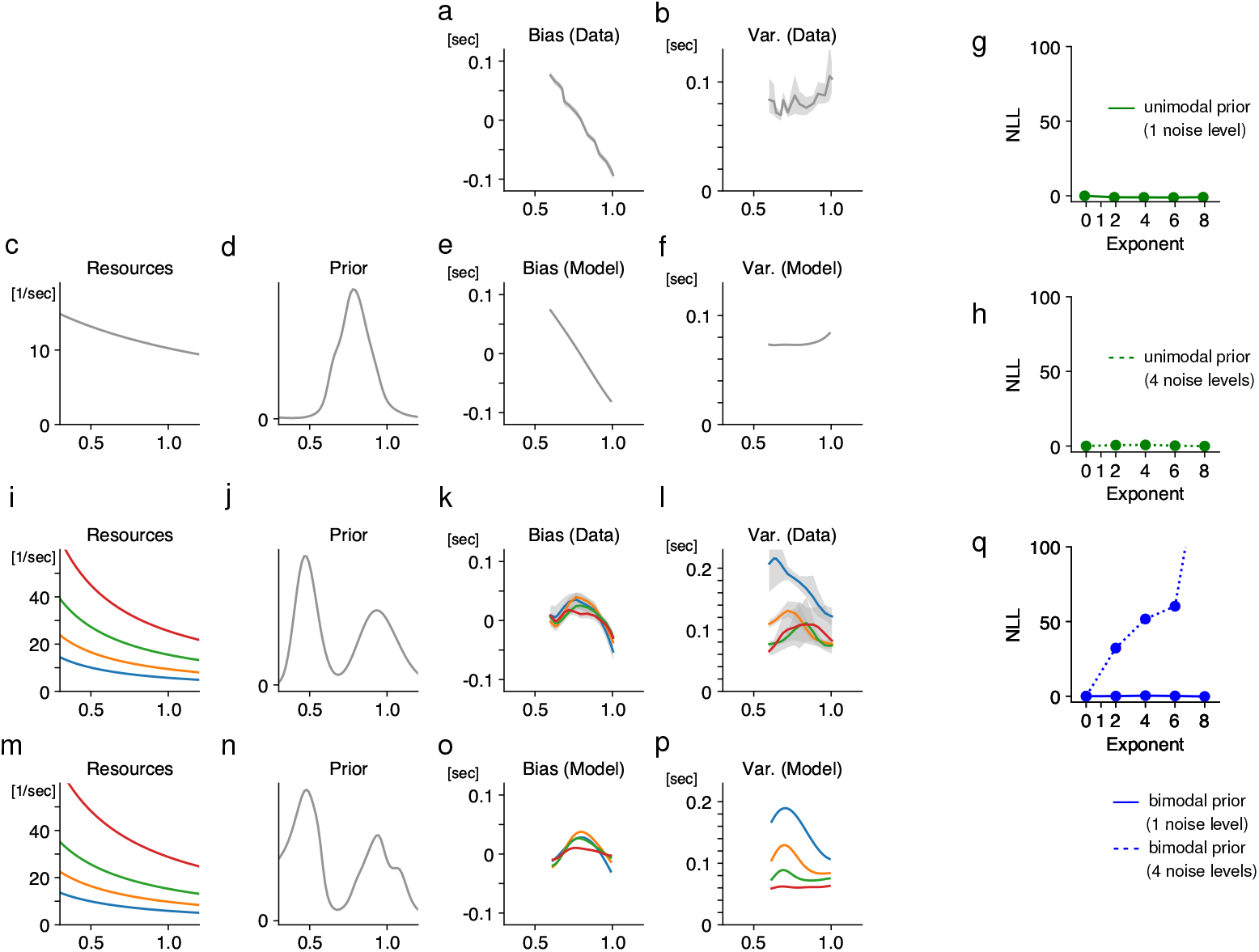
Application to experimental data: estimation of time intervals. (a-h) Analysis of the data in (38). (38) collected data at one level of sensory noise; all stimuli were from a bounded interval. Behavioral data showing a bias into the interval (a), and an increasing variability (b). This behavior is well explained in terms of an encoding consistent with Weber’s law (c) and a unimodal prior centered in the interval (d). We show a fit at *p* =0, which reproduces the bias and variability (e,f). However, even when presupposing a Weber’s law-based encoding based on prior work, the loss function cannot be identified, as the same quality of fit is achieved across loss functions (g). As the combination of a Weber’s law-based encoding and a lognormal prior is in the exceptional set Ω, the theory suggests that even an experimental design varying the sensory noise might not remedy this. Indeed, when we next simulated data from four noise levels in a model with the same Weber’s law-based encoding and a lognormal prior, the loss function remained unidentifiable (h). (i-q) Simulation results showing that the non-identifiability may be resolved using adaptation. We simulated data from four noise levels in a model with the same Weber’s law-based encoding (i) but a bimodal prior (j), corresponding to inducing a bimodal short-term prior in an adaptation phase preceding the modeled trials. We assume the ground truth loss function is *L*_0_ loss (*p* =0). The resulting simulated dataset shows biases and variability (k,l) clearly different from those produced by the unimodal prior (a,b). Our theory predicts that the overall model should typically leave the exceptional set Ω when changing the prior, making the model identifiable. Indeed, in this simulated dataset, the loss function can be uniquely identified from the dataset in (k,l) (dotted blue line in q); at *p* = 0, the prior is recovered (n) and the model fits bias and variability (o,p). In contrast, model fit results for another synthetic dataset (q) with only one level of noise (solid line) cannot single out a loss function, as predicted by our theory.

We further generated a simulated version of the dataset where the number of noise levels was increased to four (Fig. 8h). For this dataset with multiple noise levels, the model remains non-identifiable. This is expected according to our theory, since the combination of Weber’s law encoding and a lognormal prior is in the exceptional set of unidentifiable models.

If one would like to identify the loss function and the precise prior used by the subjects in this task, how could one resolve this? By our theoretical analysis, a model with the same encoding but a different prior will almost certainly be outside the exceptional set of unidentifiable models. We thus simulated responses from a hypothetical version of the dataset where subjects would have undergone adaptation to a *bimodal* prior (Fig. 8i-p) without changing the loss function. Indeed, this simulated dataset makes the loss function well identifiable when multiple noise levels are available (Fig. 8i-q), in precise agreement with our theory. These analyses provides a prediction that can be directly tested in future experiments.

## 3 Discussion

Although prior work inferred components of Bayesian observer models from behavioral data, the extent to which these models can be recovered remains poorly understood. A few studies that investigated the identifiability of Bayesian models only focused on a particular set of models under restrictive conditions (15, 13). By systematically investigating the identifiability of a general family of Bayesian models of perception and cognition, our results fill an important gap. Despite their generality, our results can be further extended to even broader settings. Below, we discuss three such directions.

First, our analytical results on identifiablity based on measurements at multiple levels require the encoding function to be shared across multiple noise levels. While this seems reasonable, for certain experimental manipulation, e.g.. changing the contrast of the stimulus, it is possible that the coding resource (or the discrimination threshold) for the stimulus may scale disproportionally. Thus, it would be useful to understand whether similar results hold when relaxing this assumption. We performed simulations to investigate this question, and find that identifiability may be preserved even in models where the encoding varies by noise level (SI Appendix, S5.3).

Second, while our results mainly address the role of sensory noise, experimentally one can also manipulate stimulus noise. Consider experiments studying the visual orientation perception (36, 12). One can use a stimulus that consists of multiple small Gabor patches with their orientations sampled from a Gaussian distribution. By varying the width of this Gaussian distribution, one can control the magnitude of the stimulus noise (36, 12). Unlike the sensory noise, stimulus noise is external, and cannot be reduced by neural processing. It is of interest to understand whether stimulus noise may benefit model identification. Theoretically, we find that having behavioral responses with non-zero stimulus noise condition in addition to those with zero-stimulus noise (i.e., pure sensory noise) makes most models identifiable, assuming that the encoding is not entirely uniform (SI Appendix, S5.2). Numerically, we also find that varying stimulus noise indeed boosts identifiability. However, when comparing the efficacy of varying the two types of noises (stimulus vs. sensory noises) in improving model identifiability, varying stimulus noise is generally less effective according to simulation results. Further investigation (SI Appendix S5.2) shows that stimulus noise reduces those components of the biases that are most revealing of the loss function while simultaneously broadening the response distribution, resulting in a less favorable signal-to-noise ratio in identifying the bias from data.

Third, while our results focused on estimation tasks, they may be generalized to another important paradigm in psychophysics, i.e., two-alternative-forced choice tasks (2-AFC). In these tasks, the observers need to compare the test stimulus with a reference to make a binary choice on every trial. To model the task, one needs to specify a decision rule. Two theoretical accounts of such tasks are common: either subjects decode 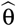 and compare it to the reference, or they directly make a binary decision based on the posterior density *P*(θ *> x*). It turns out that the latter corresponds to the settings where the loss function is already known. If both the test stimulus and the reference stimulus are subject to nonnegligible amounts of sensory noise, two levels of sensory noise for test stimuli are needed to identify the model if the loss function is known. We also find that three levels are sufficient to obtain the same asymptotic guarantees as two levels were under the estimation paradigm (SI Appendix, Section S5.1.1). If the sensory noise of the reference stimulus is negligible, the number of noise levels required for these guarantees reduces by one, matching the estimation paradigm. However, numerical simulations suggest that the 2-AFC paradigm is less efficient than the estimation paradigm in identifying models (SI Appendix, Section S5.1.2).

In this paper, we have taken the perspective that it is desirable to quantitatively recover all modeling components from the data. Empirically, the desirable notion of identiability may depend on the question under study. For instance, there may be experiments in which we are primarily interested in understanding how the coding resources are allocated along a particular stimulus dimension, but not the prior distribution or loss function. Also, for cases where the primary goal is to understand the general shape of the prior (i.e., where does the prior peak at), it may be unnecessary to precisely recover the prior. For these cases, identifiablity may be achieved more easily.

Prior work (15, 13) on the recovery of Bayesian models painted a mixed picture, suggesting that the Bayesian framework has substantial intrinsic degeneracy, though without clear theoretical understanding. Several key differences exist between their work and ours. First, (15, 13) focused on specific functional forms for encoding and prior that accommodate certain types of scalar stimuli. Specifically, (15) assume a specific logarithmic form for the encoding and represent the prior as a mixture of Gaussians, whereas we allow both priors and encodings to be arbitrary smooth functions. (15) further considered loss functions that are mixtures of two inverted Gaussians, which enabled them to obtain fast numerical solutions for the expected loss, whereas we instead consider the family of *L*_*p*_ losses.

Our results, on the one hand, confirm the presence of degeneracy when a single level of sensory noise is considered, as in (15). On the other hand, we show that, importantly, this degeneracy can be largely avoided when varying the magnitude of sensory noise. (13) empirically studied identifiability for Bayesian models in the special case of logarithmic encoding and lognormal prior. They numerically found that combining observations from two noise levels alleviated some identifiability limitations, though only loss functions based on *L*_2_ loss (with an added action cost) were considered. Our findings reveal that this is no accident: For this model, the *L*_*p*_ exponent is fundamentally unidentifiable; but it can be identified with further strategies such as adaptation to a different prior.

In contrast, our findings reveal a substantially broader, and more optimistic, picture. We provide generic identifiability theorems without parametric assumptions on encoding or prior, and without assuming the exponent of *L*_*p*_ loss to be known in advance. All three components are usually identifiable separately on the basis of estimation data alone. Interestingly, our findings reveal the previously studied combination of logarithmic encoding and log-normal prior to be a highly special case with unusually high degeneracy, while providing strategies for overcoming it.

Our work has direct implications in the experimental work that seeks to reverse engineer the perceptual decision process. First, our results on the identifiability of the encoding (i.e., Fisher information) suggest that continuous estimation task is a viable paradigm for studying how the coding resources vary as the stimulus. Most previous studies used 2-AFC to estimate the discrimination threshold (the inverse of the coding resource) for a given stimulus value. 2-AFC requires a large number of trials to estimate the discrimination threshold, thus practically only the thresholds at a small number of discrete stimulus values can be measured. Our results suggest that continuous estimation tasks offer a more sample-efficient strategy for inferring how the discrimination threshold varies along a continuous stimulus dimension (43). Second, and crucially, our results provide direct guidance on what would be effective experimental designs if the goal is to infer the components of the Bayesian observers. Our results show that it is useful to have experimental conditions that manipulate the noise characteristics of the stimuli. While these manipulations were adopted in some experimental work, their role in model identification is not recognized. Our results show that collecting behavioral responses at multiple noise levels is in fact crucial for identification of Bayesian models, in particular when the loss function is known. One interesting question for future research would be how to optimize the experimental design for recovering the modeling components for a given number of trials. In general, a better understanding of the identifiability of the Bayesian models (as well as other models used in computational cognitive neuroscience) will facilitate the interpretation of existing data and inform the design of future experiments.

## Materials and Methods

### Model of Encoding and Decoding

We assume a general encoding and decoding model for one-dimensional stimulus variables. Specifically, we assume that the stimulus space *𝒳* is a circle or a bounded interval, the space of measurement (the neural encoding signal in the brain) is denoted as *ϒ*, and the map *F* : *𝒳 → ϒ* is a bijection. Formally, the neural encoding can be described by the following equation

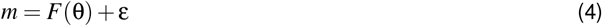

We assume, without loss of generality, that *ϒ* has volume 1. Given a particular loss function *𝓁* and an encoding model *m ∈ ϒ*, the Bayesian estimator 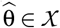 that minimizes the loss function is obtained as

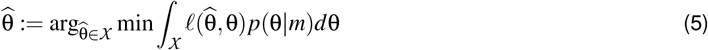

At a stimulus θ, the estimator 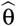 has bias 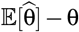.

The resource allocation is 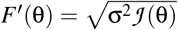, which is well-defined independently across magnitudes σ^2^ of the sensory noise.

#### Formalizing the Space of Models

As we consider the problem of identifying general observer models without fixed parametric forms for prior and encoding, the space of possible models is infinitely-dimensional. This idea can be conveniently formalized using standard concepts from functional analysis.

We take *𝒳* to be either a bounded interval or the unit circle. A function is continuous on the unit circle if it is continuous on [0, 2π)and additionally *f* (2π) = *f* (0).

Let *ℱ* (*𝒳*)be the set of real-valued functions on *𝒳* that are five times continuously differentiable, whose values are always strictly positive, and where ∫ *f* = 1. Both the prior *p*_*prior*_ and the resource allocation *F*′(θ)are elements of *ℱ* (*𝒳*).

We take *𝒫* ⊂ ℕ to be a finite set of exponents *p* for *L*_*p*_ loss functions; we assume *𝒫 ⊂* {0, 1, 2, 4, 6, …} for our theoretical results.

The space of encoding models is then the product space 𝔐 := ℱ × ℱ ×, 𝒫 whose elements are tuples of priors *p*_*prior*_, resource allocations *F*′, and loss function exponents *p*.

We quantify volume on the infinite-dimensional space 𝔐 by parameterizing both *p*_*prior*_ and *F*′ in terms of their logarithms, and placing a Gaussian process measure on these. Independently sampling prior, encoding, and loss function determines a notion of measure or volume on the space of models. See SI Appendix S2.1 for details.

### Identifiability Theorems

We consider the response distribution ℙ(*y*|*M*, θ, σ)determined by the model *M* ∈ 𝔐, θ ∈ *𝒳*, and a sensory noise magnitude σ *>* 0. We assume that these distributions are known at each stimulus θ, at one or more sensory noise levels. The noise magnitudes σ associated with the noise levels are assumed to be unknown, and inferred together with the other model components.

Our first theorem concerns recovery of the encoding, providing an explicit formula for the Fisher information in terms of the overlap in response distributions:

**Theorem 1** (Recovering Encoding from Response Distribution). *In the absence of motor noise, the encoding F and the sensory noise magnitude can be recovered from the response distribution at a single level of sensory noise, with estimation error exponentially small in* 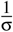:

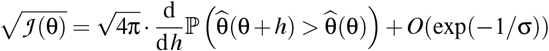

*where the derivative is evaluated at* 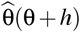 *and* 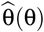 *are independent response samples at stimuli* θ +*h and* θ, *respectively*.

The proof is in SI Appendix S2.4. Numerical approximation of this formula allows straightforward estimation, independent of the loss function, as we illustrate in SI Appendix, Figure S3.

Our next theorem concerns recovery of the prior conditional on the loss function. It guarantees recovery of the prior when the loss function is known:

**Theorem 2** (One Level of Sensory Noise). *Let* σ *>* 0 *be the SD of sensory noise, and p ∈ 𝒫. Assume the response distribution* 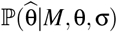 *is given at each* θ, *under a ground-truth model ⟨F*′, *p*_*prior*_, *p⟩* ∈ *𝔐. There is a functional* Φ_*p*_ *mapping these distributions to* 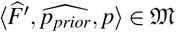 *such that*

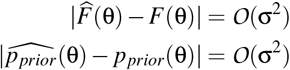

*as* σ *→* 0. *Constants in O*(*·*)*depend on F and p*_*prior*_, *but not* θ.

We next state our identifiability theorem for the more general setup where encoding, prior, and loss function are all unknown a priori. We use Ω to denote the exceptional set of unidentifiable models.

**Theorem 3** (Two Levels of Sensory Noise). *There is a subset* Ω *⊂ 𝔐 of volume* 0 *such that the following holds. Let* 0 *<* σ_1_ *<* σ_2_ *be two levels of sensory noise. Assume the response distribution* 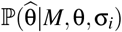 *are given at each* θ *and for both* σ_1_, σ_2_, *under a ground-truth model M* = *⟨F*′, *p*_*prior*_, *p⟩ ∈ 𝔐. There is a functional* Φ *mapping the collection of these distributions to* 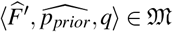 *such that—provided M ≠* Ω*—we have*

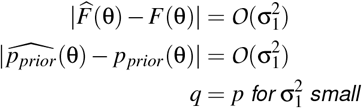

*in the limit where* σ_1_, σ_2_ *→* 0, *provided:*

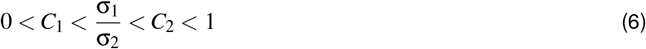

*for some C*_1_,*C*_2_. *Constants in the O*(*·*) *expressions depend on C*_1_,*C*_2_, *and on F and p*_*prior*_, *but not on* θ.

This theorem provides in-principle identification of the loss function in all models outside of Ω. Importantly, this theorem does not presuppose that the magnitudes σ_1_, σ_2_ are known, only that they are small, and that their ratio is bounded away from both 0 and 1. In practice, as the mapping between stimulus noise magnitudes σ and manipulations such as exposure durations may not be known in advance, it may not be feasible to design an experiment to have two optimal noise levels. A practical approach is thus to obtain observations at a larger number of noise levels across different scales; our numerical results show that this is generally successful in recovering the loss function.

**Theorem 4** (Role of Motor Noise). *Theorem 3 is not affected by adding symmetric isotropic motor noise of variance* τ^2^, *and by guessed responses appearing at a rate* 0 *<* γ *<* 1, *in the limit* τ, σ_1_, σ_2_ *→* 0, *while maintaining* 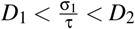 *for some* 0 *< D*_1_ *< D*_2_ *<* ∞.

The proofs of Theorems 2–4, provided in SI Appendix S2.5 and S2.6, recover model components using the Method of Moments, by matching observed moments of the response distribution to truncated Taylor expansions of the moments in powers of the noise magnitude, with coefficients reflecting the model components *F*′, *p*_*prior*_, *p*. This strategy largely recovers the ground-truth model up to an error indicating deviation from the low-noise regime. Our simulation experiments indicate that conclusions remain valid at finite noise, and when only a finite sample dataset is available.

### Fitting Procedure

We use the fitting procedure introduced in (6), which, given a loss function, jointly fits encoding, prior, motor and sensory noise magnitudes, and guessing rate, by maximizing the data’s likelihood under the model. Prior and encoding are specified on a discretized grid over *𝒳*. Following (6), its size is 180 in the models in Figures 1–7, and 200 in the model in Figure 8.

As in (6), the fitting procedure includes regularization encouraging smooth fits for prior and encoding. For synthetic data in Figures 2–7, we used the same regularization strength as (6) used for the data collected by (35). In Figure 8, we used the same regularization strength as in the fit to the original data from (6).

Note that this method is applicable even outside of the small-noise regime, allowing us to verify the generalizability of the theoretical conclusions beyond it.

### Simulations

Following (6), simulated datasets include Gaussian (or von Mises) motor noise and guesses. For all experiments on circular stimulus spaces, motor noise magnitude and guessing rate are taken from the fit to the data of (35) as obtained by (6). We also use sensory magnitudes from the same fit; focusing on the four noise levels corresponding to exposure durations of 40ms, 80ms, 160ms, 1000ms, as we find four levels to largely be sufficient for identifiability.

The numbers of trials in simulated data are as indicated in Fig. 2, 10K in Fig. 3, 10K in Fig. 4, and as indicated in Fig. 5.

For experiments on circular stimulus spaces, we focus on the orientation domain, with stimulus space [0°, 180°].

As in (6), we randomly partition each dataset into 10 folds. We fit models on 9 and evaluate negative log-likelihood (NLL) on one fold. We plot Δ NLL relative to the NLL achieved by the ground-truth loss function exponent.

For Fig. 4, we randomized models as follows. We sampled *F*′ and *p*_*prior*_ each by parameterizing their logarithms in terms of the first four terms of their Fourier series, with coefficients uniformly in [*−*0.5, 0.5].

For Fig. 5, we considered (i) uniform prior and periodic resource assignment 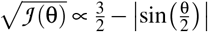 (stylized from the fit to the data collected by (35)), (ii) matched periodic prior and encoding, (iii) periodic prior and uniform encoding, (iv) a periodic encoding and a shifted prior 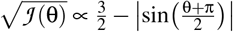. The same periodic encoding was used in Figs. 2 and 3.

In Fig. 6, we show results fitted at *p* = 8, selected based on quality of fit on the full dataset. Fit of the encoding is highly consistent even when fitting at other exponents (SI Appendix, Figure S33).

In Fig. 7, we averaged the NLL over five random seeds for downsampling the dataset.

In Fig. 8, parameters for the unimodal model are taken from the fit to the behavioral data, which is from (6). Parameters for the bimodal model are identical except for the prior. In Fig. 8, all fits presuppose a logarithmic encoding consistent with Weber’s law, as in the analysis of the dataset in (6). Corresponding model fit statistics for freely fitted encodings (SI Appendix, Fig. S34) show that the bimodal model is identifiable even without assuming a specific encoding.

### Experimental data

The experimental data used in this paper are published in ref (35) and (38). Data collection and analysis were not performed blind to the conditions of the experiments.

Orientation perception (35): 49 subjects adjusted a blue strip in order to reproduce the orientation of a Gabor patch. Stimuli were uniformly distributed on most trials, but had orientation 0, 45, 90, 135 degrees on a specific subset of trials; we followed (6) in excluding the latter. Depending on the trial, exposure duration was 20, 40, 80, 160, or 1000 ms. On 10% of trials, no stimulus was shown; we excluded these. There were 2,208 trials at 20ms, 40ms, 80ms, 160ms each, and 1,104 trials at 1000ms. In total, 9,936 trials were included.

Time intervals (38): 15 subjects reproduced time intervals in a Ready-Set-Go Task in Experiments 1 and 2. Sessions differed in whether subjects reproduced the time interval itself (the Identity condition) or reproduced it scaled by factors of 0.75 or 1.5; we only included data in the identity condition. In total, 8,999 trials were included.

## Acknowledgements

This research uses data from a number of previous published studies. We would like to thank the authors of these studies for sharing their data or making their data publicly available. We thank Alan Stocker and Pascal Mamassian for helpful discussions. X.X.W. is supported by a Sloan Research Fellowship from the Alfred P. Sloan Foundation. M.H. and E.W. are supported by startup funding from Saarland University.

## Author Contributions

M.H. and X.X.W. conceived and designed the research. M.H. and X.X.W. developed the theoretical framework. M.H. and E.W. performed the theoretical, numerical, and data analyses, with inputs from X.X.W.. M.H. and X.X.W. interpreted the results and wrote the paper.

## Competing Interests

The authors declare no competing interests.

## Notes

### Competing Interest Statement

The authors have declared no competing interest.

## References

[1] Konrad P Körding and Daniel M Wolpert. Bayesian integration in sensorimotor learning. Nature, 427(6971):244–247, 2004.

[2] Y. Weiss, Eero P. Simoncelli, and Edward H. Adelson. Motion illusions as optimal percepts. Nature Neuroscience, 5:598–604, 2002.

[3] Alan A. Stocker and Eero P. Simoncelli. Noise characteristics and prior expectations in human visual speed perception. Nature Neuroscience, 9:578–585, 2006.

[4] Wilson S Geisler. Contributions of ideal observer theory to vision research. Vision research, 51(7):771–781, 2011.

[5] Xue-Xin Wei and A. Stocker. A bayesian observer model constrained by efficient coding can explain ‘anti-bayesian’ percepts. Nature Neuroscience, 18:1509–1517, 2015.

[6] Michael Hahn and Xue-Xin Wei. A unifying theory explains seemingly contradictory biases in perceptual estimation. Nature Neuroscience, 27(4):793–804, 2024.

[7] Joshua B Tenenbaum, Thomas L Griffiths, and Charles Kemp. Theory-based bayesian models of inductive learning and reasoning. Trends in cognitive sciences, 10(7):309–318, 2006.

[8] Thomas L Griffiths and Joshua B Tenenbaum. Optimal predictions in everyday cognition. Psychological science, 17(9):767–773, 2006.

[9] David C Knill and Alexandre Pouget. The bayesian brain: the role of uncertainty in neural coding and computation. TRENDS in Neurosciences, 27(12):712–719, 2004.

[10] David C. Knill and Whitman Richards, editors. Perception as Bayesian Inference. 1996.

[11] Alan A Stocker and Eero P Simoncelli. Noise characteristics and prior expectations in human visual speed perception. Nature neuroscience, 9(4):578–585, 2006.

[12] Ahna Reza Girshick, Michael S. Landy, and Eero P. Simoncelli. Cardinal rules: Visual orientation perception reflects knowledge of environmental statistics. Nature neuroscience, 14:926 – 932, 2011.

[13] Dominik Straub, Tobias F Niehues, Jan Peters, and Constantin A Rothkopf. Inverse decision-making using neural amortized bayesian actors. In The Thirteenth International Conference on Learning Representations, 2025.

[14] Tyler S Manning, Benjamin N Naecker, Iona R McLean, Bas Rokers, Jonathan W Pillow, and Emily A Cooper. A general framework for inferring bayesian ideal observer models from psychophysical data. eneuro, 10(1), 2023.

[15] Luigi Acerbi, Wei Ji Ma, and Sethu Vijayakumar. A framework for testing identifiability of bayesian models of perception. Advances in neural information processing systems, 27, 2014.

[16] Konrad Paul Körding and Daniel M Wolpert. The loss function of sensorimotor learning. Proceedings of the National Academy of Sciences, 101(26):9839–9842, 2004.

[17] Rafael Polanía, Michael Woodford, and Christian C. Ruff. Efficient coding of subjective value. Nature neuroscience, 22:134 – 142, 2018.

[18] Miguel Barretto-García, Gilles de Hollander, Marcus Grueschow, Rafael Polanía, Michael Woodford, and Christian C Ruff. Individual risk attitudes arise from noise in neurocognitive magnitude representations. Nature Human Behaviour, 7(9):1551–1567, 2023.

[19] Pascal Mamassian and Michael S Landy. It’s that time again. Nature neuroscience, 13(8):914–916, 2010.

[20] David C Knill and Jeffrey A Saunders. Do humans optimally integrate stereo and texture information for judgments of surface slant? Vision research, 43(24):2539–2558, 2003.

[21] Johannes Burge, Charless C Fowlkes, and Martin S Banks. Natural-scene statistics predict how the figure– ground cue of convexity affects human depth perception. Journal of Neuroscience, 30(21):7269–7280, 2010.

[22] Julia Trommershäuser, Laurence T Maloney, and Michael S Landy. Statistical decision theory and the selection of rapid, goal-directed movements. JOSA A, 20(7):1419–1433, 2003.

[23] Jeffrey S Bowers and Colin J Davis. Bayesian just-so stories in psychology and neuroscience. Psychological bulletin, 138(3):389, 2012.

[24] Thomas L Griffiths, Nick Chater, Dennis Norris, and Alexandre Pouget. How the bayesians got their beliefs (and what those beliefs actually are):comment on bowers and davis (2012). 2012.

[25] Ronald Aylmer Fisher. Theory of statistical estimation. In Mathematical proceedings of the Cambridge philosophical society, volume 22, pages 700–725. Cambridge University Press, 1925.

[26] Erich L Lehmann and George Casella. Theory of point estimation. Springer Science & Business Media, 2006.

[27] Eric Walter. Identifiability of parametric models. Elsevier, 2014.

[28] Tjalling C Koopmans. Identification problems in economic model construction. Econometrica, Journal of the Econometric Society, pages 125–144, 1949.

[29] Joseph P Romano and Azeem M Shaikh. Inference for identifiable parameters in partially identified econometric models. Journal of Statistical Planning and Inference, 138(9):2786–2807, 2008.

[30] Alejandro F Villaverde, Antonio Barreiro, and Antonis Papachristodoulou. Structural identifiability of dynamic systems biology models. PLoS computational biology, 12(10):e1005153, 2016.

[31] Franz-Georg Wieland, Adrian L Hauber, Marcus Rosenblatt, Christian Tönsing, and Jens Timmer. On structural and practical identifiability. Current Opinion in Systems Biology, 25:60–69, 2021.

[32] Xue-Xin Wei and Michael Hahn. The identifiability of bayesian models of perceptual decision. Journal of Vision (Vision Sciences Society Annual Meeting Abstract), 24(10):572–572, 2024.

[33] S. Laughlin. A simple coding procedure enhances a neuron’s information capacity. Zeitschrift für Naturforschung C, 36:910 – 912, 1981.

[34] Anthony J Bell and Terrence J Sejnowski. An information-maximization approach to blind separation and blind deconvolution. Neural computation, 7(6):1129–1159, 1995.

[35] Vincent de Gardelle, Sid Kouider, and Jérôme Sackur. An oblique illusion modulated by visibility: non-monotonic sensory integration in orientation processing. Journal of vision, 10 10:6, 2010.

[36] Alessandro Tomassini, Michael J. Morgan, and Joshua A. Solomon. Orientation uncertainty reduces perceived obliquity. Vision Research, 50:541–547, 2010.

[37] Mehrdad Jazayeri and Michael N. Shadlen. Temporal context calibrates interval timing. Nature neuroscience, 13:1020–1026, 2010.

[38] Evan D. Remington, Tiffany V Parks, and Mehrdad Jazayeri. Late bayesian inference in mental transformations. Nature Communications, 9, 2018.

[39] Frederike H. Petzschner, Stefan Glasauer, and Klaas Enno Stephan. A bayesian perspective on magnitude estimation. Trends in Cognitive Sciences, 19:285–293, 2015.

[40] Stuart Appelle. Perception and discrimination as a function of stimulus orientation: the” oblique effect” in man and animals. Psychological bulletin, 78(4):266, 1972.

[41] Neil W Roach, Paul V McGraw, David J Whitaker, and James Heron. Generalization of prior information for rapid bayesian time estimation. Proceedings of the National Academy of Sciences, 114(2):412–417, 2017.

[42] Yang Xiang, Thomas Graeber, Benjamin Enke, and Samuel J. Gershman. Confidence and central tendency in perceptual judgment. Attention, perception & psychophysics, 2021.

[43] Jean-Paul Noel, Ling-Qi Zhang, Alan A Stocker, and Dora E Angelaki. Individuals with autism spectrum disorder have altered visual encoding capacity. PLoS biology, 19(5):e3001215, 2021.

