## Supplementary material for "Identifiability of Bayesian Models of Cognition": SI Appendix

### Identifiability of Bayesian Models of Cognition
### Supporting Information

Michael Hahn
Entang Wang
Xue-Xin Wei

June 25, 2025

**S6 Applications to Experimental Data** **74**

### S1 Analytically Solvable Models

Here, we show the optimality of our results by considering analytically tractable models.

#### S1.1 Logarithmic Encoding, Log-Normal Prior (Loss Function and Prior Confounded)

We consider a model with logarithmic encoding and log-normal prior, a simple model for scalar stimulus variables satisfying Weber's Law, where the log-normal prior represents a stimulus distribution concentrated in an interval [16, 5]. Remarkably, we will show that this model family[1] is entirely contained in the exceptional set $\Omega$ studied in Theorem 2, and, indeed, *no number of sensory noise levels* will be sufficient for disentangling prior and loss function in this model family. An important implication is thus that $\Omega$, even though it is very small in a mathematically precise sense, may still contain such very natural models. We discuss strategies for recovering identifiability in Section S2.7. Define the log-normal density as

$$LogNormal(M,V)(\theta) := \frac{1}{\theta}\exp\left(-\frac{(\log\theta - M)^2}{2V}\right) \tag{1}$$

Then set:

$$\text{(Prior)} \qquad p_{prior}(\theta) \propto LogNormal(\log\mu, \sigma^2) := \frac{1}{\theta}\exp\left(-\frac{(\log\theta - \log\mu)^2}{2\sigma^2}\right)$$

$$\text{(Encoding)} \qquad F(\theta) = \log\theta \qquad (\text{hence, } F'(\theta) = \frac{1}{\theta})$$

on the stimulus space $\mathcal{X} = (0,\infty)$ and the corresponding sensory space $\mathcal{Y} = \mathbb{R}$. Note that this space is infinite, unlike the other setups we consider; we comment on this below. We now show that, for such models, the location parameter $\mu$ is confounded with the loss function:

**Theorem S1.** *Given $p \in \mathbb{N}$, define $\tilde{p}$ as in Equation 36. Let $\mu, \sigma^2 > 0$. The family of models*

$$\{\langle F'(\theta) = \frac{1}{\theta}, p_{prior} = LogNormal\left(\log\mu - \frac{\tilde{p}}{2}\sigma^2, \sigma^2\right), p\rangle : p = 0,1,2,4,\dots\} \tag{2}$$

*defines the same response distribution for any $p$ and any sensory noise magnitude $t$. Hence, no amount of trials and sensory noise levels can distinguish between these models.*

*Proof.* For the model

$$\langle F'(\theta) = \frac{1}{\theta}, p_{prior} = LogNormal\left(\log\mu - \frac{\tilde{p}}{2}\sigma^2, \sigma^2\right), p\rangle \tag{3}$$

the encoding likelihood transformed into stimulus space is

$$p(m|\theta) \propto LogNormal(\log\theta, t) := \frac{1}{m}\exp\left(-\frac{(m - \log\theta)^2}{2t}\right). \tag{4}$$

[1] Strictly speaking, restrictions of this model to a finite stimulus space interval.

Hence, the posterior is

$$p(\theta|m) = LogNormal(M,V)(\theta) \tag{5}$$

with parameters

$$V := \frac{1}{\frac{1}{\sigma^2} + \frac{1}{t}} \qquad\qquad M := V \cdot \left( \frac{\log m}{t} + \frac{\log\mu - \tilde{p}\sigma^2/2}{\sigma^2} \right)$$

The $L_p$ estimator is

$$\begin{aligned}
\exp\left(M + \frac{\tilde{p}-1}{2}V\right) &= \exp\left(V \cdot \left[\left(\frac{\log m}{t} + \frac{\log\mu - \tilde{p}\sigma^2/2}{\sigma^2}\right) + \frac{\tilde{p}-1}{2}\right]\right) \\
&= \exp\left(V \cdot \left[\frac{\log m}{t} + \frac{\log\mu}{\sigma^2} - \frac{\tilde{p}}{2} + \frac{\tilde{p}-1}{2}\right]\right) \\
&= \exp\left(V \cdot \left[\frac{\log m}{t} + \frac{\log\mu}{\sigma^2} - \frac{1}{2}\right]\right)
\end{aligned}$$

which is independent of $\tilde{p}$. For $p = 0,1,2$, the formula for the $L_p$ estimator follows from the standard formulas for mode, median, and mean of the lognormal distribution. For higher even exponents, this follows from Lemma S3 below.

□

We note that the transformation of the prior in the family (2) is consistent with Eq. 3 in the main paper: multiplying a log-normal density with a power of θ amounts to a shift of the density's location parameter:

**Lemma S2.**

$$G(\theta) := p_{prior}(\theta)F'(\theta)^{\frac{p-p'}{2}} \propto LogNormal\left(\log\mu - \frac{p-p'}{2}\sigma^2, \sigma^2\right)$$

*Proof.* Define

$$\delta = \frac{p-p'}{2}.$$

Then, we can rewrite $G(\theta)$ as

$$G(\theta) = \frac{1}{\theta^{1+\delta}} \exp\left(-\frac{(\log\theta - \log\mu)^2}{2\sigma^2}\right) = \frac{1}{\theta}\exp\left[-\frac{(\log\theta - \log\mu)^2}{2\sigma^2} - \delta\log\theta\right].$$

Notice that $\delta\log\theta = \frac{p-p'}{2}\log\theta$. Thus, the exponent becomes

$$-\frac{(\log\theta - \log\mu)^2}{2\sigma^2} - \delta\log\theta = -\frac{1}{2\sigma^2}\left[(\log\theta - \log\mu)^2 + 2\delta\sigma^2\log\theta\right].$$

Expanding the square, combining the linear terms in $\log\theta$, and completing the square for the quadratic in $\log\theta$, we have:

$$\begin{aligned}
(\log\theta - \log\mu)^2 + 2\delta\sigma^2\log\theta &= (\log\theta)^2 - 2\log\mu\log\theta + (\log\mu)^2 + 2\delta\sigma^2\log\theta \\
&= (\log\theta)^2 - 2\left(\log\mu - \delta\sigma^2\right)\log\theta + (\log\mu)^2 \\
&= \left(\log\theta - \left(\log\mu - \delta\sigma^2\right)\right)^2 - \left(\log\mu - \delta\sigma^2\right)^2 + (\log\mu)^2.
\end{aligned}$$

Thus, up to a multiplicative constant (which absorbs the θ-independent terms), we obtain

$$G(\theta) \propto \frac{1}{\theta}\exp\left[-\frac{\left(\log\theta - \left(\log\mu - \delta\sigma^2\right)\right)^2}{2\sigma^2}\right] = LogNormal\left(\log\mu - \frac{p-p'}{2}\sigma^2, \sigma^2\right).$$

□

It remains to show:

**Lemma S3.** *Let $X$ be a lognormal random variable with $\ln X \sim N(\mu, \sigma^2)$, and let $p \geq 2$ be an even integer. Then the unique minimizer of*

$$f(y) = \mathbb{E}\big[(X-y)^p\big]$$

*over $y \in \mathbb{R}$ is*

$$y^* = \exp\Big(\mu + \frac{p-1}{2}\sigma^2\Big).$$

This generalizes the standard formula for the mean and median of lognormal random variables; if one replaces $p=0$ with $\tilde{p} = -1$, it also covers the mode.

*Proof.* Due to strong convexity,

$$\mathbb{E}\big[(X-y)^p\big] \tag{6}$$

has a unique stationary point, where it attains its minimum. Setting $f'(y) = 0$ yields

$$\mathbb{E}\big[(X-y)^{p-1}\big] = 0. \tag{7}$$

We now write $X$ in its lognormal form:

$$X = e^{\mu+\sigma Z}, \quad \text{with } Z \sim N(0,1).$$

Substituting the ansatz

$$y = \exp\Big(\mu + c\,\sigma^2\Big),$$

(for some constant $c$ to be determined) into Eq. 7, we obtain

$$\mathbb{E}\Big[\Big(e^{\mu+\sigma Z} - \exp\big(\mu + c\,\sigma^2\big)\Big)^{p-1}\Big] = 0.$$

Factor out $e^{\mu}$:

$$\mathbb{E}\Big[\Big(e^{\sigma Z} - \exp\big(c\,\sigma^2\big)\Big)^{p-1}\Big] = 0.$$

Denote $M = \exp\big(c\,\sigma^2\big)$ so that the condition becomes

$$\mathbb{E}_Z\left[\Big(e^{\sigma z} - M\Big)^{p-1}\right] = 0.$$

Since $p-1$ is odd (because $p$ is even), we expand the integrand using the binomial theorem:

$$\Big(e^{\sigma z} - M\Big)^{p-1} = \sum_{k=0}^{p-1} \binom{p-1}{k} (-M)^{p-1-k} e^{k\sigma z}.$$

Taking expectation term-by-term[2], we have

$$\sum_{k=0}^{p-1} \binom{p-1}{k} (-M)^{p-1-k} \exp\Big(\frac{1}{2}k^2\sigma^2\Big) = 0.$$

Replacing $M$ by $\exp\big(c\,\sigma^2\big)$, this sum becomes

$$\sum_{k=0}^{p-1} \binom{p-1}{k} (-1)^{p-1-k} \exp\Big(c\,\sigma^2(p-1-k) + \frac{1}{2}k^2\sigma^2\Big) = 0.$$

[2]Recalling that

$$\mathbb{E}[e^{k\sigma Z}] = \exp\Big(\frac{1}{2}k^2\sigma^2\Big)$$

by the moment generating function of the normal distribution.

Now set

$$c = \frac{p-1}{2},$$

The key idea is now to show that the terms in the sum pair up and cancel. For each $k$ there is a corresponding term with $k' = (p-1) - k$. The exponent becomes, upon substituting $c = \frac{p-1}{2}$,

$$\Phi(k) := c\sigma^2(p-1-k) + \frac{1}{2}k^2\sigma^2 = \frac{p-1}{2} \cdot \sigma^2(p-1-k) + \frac{1}{2}k^2\sigma^2 = \frac{\sigma^2}{2}\Big((p-1)^2 - (p-1)k + k^2\Big).$$

This expression is symmetric in $k$ and $p-1-k$:

$$\begin{aligned}
\Phi(p-1-k) = \frac{\sigma^2}{2}\Big((p-1)^2 - (p-1)(p-1-k) + (p-1-k)^2\Big) &= \frac{\sigma^2}{2}\Big((p-1)(k) + (p-1)^2 - 2(p-1)k + k^2\Big) \\
&= \frac{\sigma^2}{2}\Big((p-1)^2 - (p-1)(k) + k^2\Big) \\
&= \Phi(k)
\end{aligned}$$

and the alternating signs $(-1)^{p-1-k}$ (recall that $p$ is even) ensure that the contributions of paired terms cancel. Thus, the sum is zero, and Eq. 7 is satisfied. Therefore, the choice

$$y^* = \exp\Big(\mu + \frac{p-1}{2}\sigma^2\Big)$$

minimizes the function $f(y)$. □

We note that the models defined in this section have an infinite sensory space, with infinite precision as $\theta \to 0$ and an infinite aggregate amount of coding resources, expressed by the infinite volume of $\mathcal{Y}$. In reality, one may however expect human subjects to have a finite coding capacity; indeed, infinite resources are not feasible in a finite population code with nonzero noise. Indeed, when fitting such models on behavioral data, the sensory space is generally truncated [16]; in the related domain of log-odds encodings of probabilities, [29] even found finite truncation of the sensory space to be important for fitting behavioral data. In such cases, deviations from the infinite sensory noise space might be beneficial for identifiability by leading to deviations from the theoretically unidentifiable model.

#### S1.2 Gaussian Prior and Uniform Encoding (Prior Identifiable, Loss Function Unidentifiable)

Traditional Bayesian models of cognition have sometimes considered a uniform encoding and Gaussian prior. In such a model, the loss function is not identifiable, but the prior can be identified uniquely nonetheless. This is a unique special case and can only happen for a uniform encoding, as this is the only situation where the bias in the small-noise regime is invariant to changing the loss function.

Likelihood:

$$p(m|\theta) \propto \exp\left(-\frac{(m-\theta)^2}{2\sigma^2}\right) \tag{8}$$

Prior:

$$p(\theta) \propto \exp\left(-\frac{\theta^2}{2\tau^2}\right) \tag{9}$$

Posterior:

$$p(\theta|m) \propto \exp\left(-\frac{1}{2}\frac{\left(\theta - \frac{m\tau^2}{\sigma^2+\tau^2}\right)^2}{\frac{\sigma^2\tau^2}{\sigma^2+\tau^2}}\right). \tag{10}$$

This is symmetric in $\theta$; hence, in this special case, all $L_p$ estimators provide the same result. Hence, the loss function is not identifiable in this setup. However, the prior is uniquely identifiable nonetheless. In situations where the prior (but not the loss function) is of primary interest, situations with a uniform encoding may thus be particularly advantageous.

##### S1.3 Asymmetric Prior and Uniform Encoding (Prior Identifiable at 1 Level)

If the prior is asymmetric or multimodal, but the encoding is uniform, the prior is always identifiable, because the transformation from Formula 3 in the main paper leaves the prior unchanged. The loss function will also be identifiable barring special cases. This is because the posterior will be asymmetric in general, and different $L_p$ loss functions will generally pick out different points on an asymmetric posterior. The bias and variability have the form

$$\begin{aligned} var(\theta) =& \sigma^2 + O(\sigma^4) \\ bias(\theta) =& \sigma^2 (\log p_{prior})' + O(\sigma^4) \end{aligned}$$

Even with observations from a single level of sensory noise, the prior is identifiable with error $O(\sigma^2)$ (because the encoding is uniform), but the loss function cannot be recovered from this.

### S2 Identifiability Theorems

#### S2.1 Formalization of Model Space and Measure

##### S2.1.1 Regularity Assumptions

We will make the following assumptions, which are satisfied in most Bayesian observer models constructed in the literature.

1. $\mathcal{X}$ is a closed interval (interval space) or a circle (circular space)
2. $\mathcal{Y}$ has the same topology as $\mathcal{X}$; without loss of generality, we assume its size is 1.
3. $F'$ and $p_{prior}$ are five times continuously differentiable.
4. $F'$ is nowhere zero or infinite
5. $p_{prior}$ is nowhere zero or infinite

**Note on Differentiability** We note that we expect five-times continuous differentiability for Theorem 3; in fact, for the prior, three orders of differentiability would be sufficient, but we assume the same order for simplicity. We also note that fewer orders of differentiability are required for Theorem 2. Because the density of the prior distribution is almost always assumed to be arbitrarily smooth in Bayesian observer models developed in the literature, the assumption of five-times continuous differentiability is rather mild.

In our numerical experiments, we use some models (naturalistic periodic encoding/prior) whose components are not necessarily even once differentiable, without apparent detriment to identifiability. We note that such functions can be approximated very closely for all practical purposes with such smooth functions, i.e., up to very small approximation errors that are unlikely to matter practically, our guarantees extend to such functions.

Importantly, we believe that real-world models will be smooth for almost all stimulus points, with isolated points of reduced smoothness (such as the peaks in these periodic functions). Our identifiability guarantees apply locally: around any point, they allow identifying encoding and prior from the local changes in variability and bias, and they furthermore identify the loss function using the behavior locally around any point where prior or encoding vary, because they rely on checking the second-order component $D_M$ around any stimulus. Hence, our guarantees extend to encodings and priors that have isolated points of reduced smoothness. A formalization in terms of more general function spaces would be feasible, but we believe that our treatment in terms of smooth functions strikes a good balance between generality and complexity.

**Note on Boundaries** When $\mathcal{X}$ is an interval, Bayesian decoding shows special behavior for stimuli whose distance from the boundary is on the order of $\sim \frac{1}{\sqrt{\mathcal{J}}}$, as formalized in Theorem 3 of Hahn and Wei [5]. Our identifiability theory applies to points in the interior $(0,1)$, where this boundary effect vanishes for sufficiently small noise. As encoding and prior are continuous, this is sufficient to reveal them on the whole interval $[0,1]$. Our simulations in Section S4.3 take the boundary effect into account. We expect that the boundary effect, as it is loss-function dependent, might provide additional identifiability benefit.

###### S2.1.2 Model of Encoding-Decoding Cascade

We assume the following standard encoding-decoding cascade, spelled out here for reference. We are given a stimulus (measurement) space $\mathcal{X}$, assumed to be a circle or a bounded closed interval. The encoding $F$ establishes a smooth bijection from $\mathcal{X}$ to the sensory (internal) space $\mathcal{Y}$. A stimulus $\theta \in \mathcal{X}$ is encoded as

$$m = F(\theta) + \varepsilon \tag{11}$$

where $\varepsilon \sim \mathcal{N}(0, \sigma^2)$ denotes sensory noise. If $\mathcal{X}$ (and thus $\mathcal{Y}$) is a bounded interval, sensory noise is truncated to $\mathcal{Y}$. If $\mathcal{X}$, $\varepsilon$ is instead von Mises with concentration parameter $1/\sigma^2$. The encoding $m$ is then decoded into the Bayesian estimate

$$\widehat{\theta} = \arg_{\widehat{\theta} \in \mathcal{X}} \min \int \ell(\hat{\theta}, \theta) P(\theta|m) d\theta \tag{12}$$

where $P(\theta|m)$ is the posterior, and $\ell$ is an $L_p$ loss function. With probability $\gamma \in (0,1)$, the response is uniform on $\mathcal{X}$ (guess). With probability $1-\gamma$, the response is $\widehat{\theta} + \varepsilon_{Motor}$, where $\varepsilon_{Motor} \sim \mathcal{N}(0, \tau^2)$ (or analogously von Mises if $\mathcal{X}$ is a circle).

###### S2.1.3 Formalizing the Space of Models

As we consider the problem of identifying general observer models without fixed parametric forms for prior and encoding, the space of possible models is infinitely-dimensional. This idea can be conveniently formalized using standard concepts from functional analysis.

We take $\mathcal{X}$ is either a bounded interval $[0,1]$ or the circle, formally the space $\mathbb{T}^1$ obtained from the interval $[0, 2\pi)$ by identifying $2\pi$ with 0. A function is continuous on $\mathbb{T}^1$ if it is continuous on $[0, 2\pi)$ and additionally $f(2\pi) = f(0)$.

**Definition S4.** *Let $\mathcal{F}(\mathcal{X})$ the set of real-valued functions $f : \mathcal{X} \rightarrow \mathbb{R}$ that are five times continuously differentiable, whose values are always strictly positive, and where $\int_{\mathcal{X}} f = 1$.*

Both the prior $p_{prior}(\theta)$, and the resource allocation $F'(\theta)$ are of this form. Also, $\mathcal{P} \subset \{0, 1, 2, 4, 6, \dots\}$ is the set of loss function exponents, which we take to be finite. The precise set is not important for our theoretical results. In particular, we include the Posterior Mean ($p = 2$; e.g. [6]), Median ($p = 1$), MAP ($p = 0$; e.g. [22]) estimators, and higher even-order $L_p$ estimators such as $p = 4$ or $p = 8$, found to provide better fit in orientation and direction perception [5].

Overall, we define the space of model as the Cartesian product of two copies of $\mathcal{F}(\mathcal{X})$, with the set of exponents:

**Definition S5.**

$$\mathfrak{M} := \mathcal{F}(\mathcal{X}) \times \mathcal{F}(\mathcal{X}) \times \mathcal{P} \tag{13}$$

*where the first component describes $F'$, the second component describes $p_{prior}$, and the third component describes the loss function.*

###### S2.1.4 Measure on the Space of Models.

In order to quantify the volume occupied by unidentifiable models, we aim to define a measure on $\mathfrak{M}$. As $\mathcal{F}(\mathcal{X})$ is infinite-dimensional, we formalize this using Gaussian process constructions that permit representing any function, with higher weight placed on smoother functions but no other constraints. This measure will assign the weight of 1 to the overall space, allowing a simple probabilistic interpretation in terms of randomly choosing encodings, priors, and loss functions:

**Informal Assumption S6.** *Intuitively, $\mu$ describes the probability that one encounters a given model $\langle F', p_{prior}, p \rangle$ by independently sampling $F'$ and $p_{prior} \in \mathcal{F}(\mathcal{X})$ (both as random smooth functions), and $p \in \mathcal{P}$.*

For any (formally, any measurable) subset $A \subseteq \mathfrak{M}$, $\mu(A)$ describes the probability of obtaining a model in this set. In particular, for the set $\Omega$ of possibly unidentifiable models (Theorem 3), $\mu(\Omega)$ describes the probability that a randomly constructed model may be unidentifiable. By saying that $\Omega$ has measure zero ($\mu(\Omega) = 0$), we say that the chance that one encounters an unidentifiable model when randomly constructing prior, encoding, and loss is extremely small. We note that a key assumption here is that prior and encoding are constructed independently. We examine the situation where they are dependent (e.g., due to efficient coding) in Section S2.7; $\Omega$ also has zero measure under such settings.

**Formal Definition** We now make Assumption S6 formal, in Assumptions S7 and S8. As the elements of $\mathcal{F}(\mathcal{X})$ are constrained to be positive and integrate to 1, it is most convenient to parameterize them in terms of their logarithms.[3] We parameterize the elements of $\mathcal{F}(\mathcal{X})$ in terms of their logarithms, with a normalization term subtracted for uniqueness:

$$f \mapsto L(f) := \text{ where } L(f)(\theta) = \log f(\theta) - \log f(0)$$

If $f \in \mathcal{F}(\mathcal{X})$, then $L(f)$ is a general $k$-times differentiable function with $L(f)(0) = 0$. [4] We then define a measure on $f \in \mathcal{F}(\mathcal{X})$ by placing a standard Gaussian process on $L(f)$. Formally:

**Assumption S7.** *We consider a measure on $\mathcal{F}(\mathcal{X})$ under which $L(f)$ and its first through fifth derivatives ($L(f)$, $L(f)^{(1)}$, ..., $L(f)^{(5)}$) are jointly Gaussian, with non-degenerate covariance. That is, for each finite collection $x_1, \dots, x_t \in \mathcal{X}$, the vector $(L(f)^{(r)}(x_s) : s = 1, \dots, t; r = 0, \dots, 5)$ is Gaussian with a nondegenerate covariance matrix.*

In both $\mathbb{T}$ and $[0,1]$, there are standard constructions using orthonormal basis functions for the space of functions $L(f)$, with Gaussian coefficients grounded in the Karhunen-Loève decomposition, a probabilistic analogue of the Fourier series [7, 10, 15]. For $\mathcal{X} = \mathbb{T}$, this corresponds to a Fourier series with random coefficients.[5] On an interval stimulus space $\mathcal{X} = [0,1]$, a representation using Brownian motion is available. We sample initial values $y(0), y'(0), \dots, y^{(m)}(0)$ from a Gaussian, and place a Brownian motion prior on $f(\theta) := y^{(m)}(\theta) - y^{(m)}(0)$. An explicit series representation with Gaussian coefficients can be obtained from the Karhunen-Loeve representation of Brownian motion. In both cases, any continuous function $f$[6] with $f(0) = 0$ can be represented for some choice of the random coefficients in the series representations; the coefficients are scaled to ensure that $f$ is five times differentiable almost surely.

The precise definition of the Gaussian measures on $\mathcal{F}(\mathcal{X})$ is not important for our results. What is, however, important is the assumption of non-degeneracy. This informally means that $L(f)$ and its derivatives at some stimulus $\theta$ cannot uniquely determine the corresponding values at some other stimulus $\theta'$.

**Assumption S8.** *We define a measure $\mu$ on $\mathfrak{M}$ as the product measure of the measure from Assumption S7 on the $\mathcal{F}(\mathcal{X})$ for the first two components, and the uniform measure on $\mathcal{P}$ for the third component.*

[3] In fact, this corresponds to the parameterization used in the implemented fitting procedure.

[4] This is a 1-to-1 correspondence with inverse:

$$f \mapsto softmax(f) \text{ where } softmax(f)(\theta) = \frac{\exp(f(\theta))}{\int \exp(f(s))ds}$$

[5] To define a Gaussian process over $m$-times continuously differentiable functions on $\mathbb{T}$, set

$$f = \sum_{n=1}^{\infty} \left( \frac{X_n}{n^{m+1}} \cos(nx) + \frac{Y_n}{n^{m+1}} \sin(nx) \right) \tag{14}$$

where $X_n, Y_n \sim \mathcal{N}(0,1)$.

[6] Formally, even every element of $L^2(\mathcal{X})$.

#### S2.2 Background on Bias and Variability

Here, we provide background from [5] relevant to Theorems 1 and 2. As described in the main paper, it is convenient to represent the MAP estimator using $-1$ instead of 0 as a convention (so the expression for the bias matches the pattern for the other exponents). Formally,

$$\tilde{p} := \begin{cases} p & \text{if } p > 0 \\ -1 & \text{if } p = 0 \end{cases} \tag{15}$$

Then, the bias is given as (Theorem 1 of [5]):

$$bias(\theta) = \frac{1}{\mathcal{J}(\theta)} \frac{d}{d\theta} \log p_{prior}(\theta) + \frac{\tilde{p}+2}{4} \frac{d}{d\theta} \frac{1}{\mathcal{J}(\theta)} + O(\sigma^4) \tag{16}$$

or equivalently

$$bias(\theta) = \frac{\sigma^2}{F'(\theta)^2} \frac{d}{d\theta} \log p_{prior}(\theta) + \frac{\tilde{p}+2}{4} \frac{d}{d\theta} \frac{\sigma^2}{F'(\theta)^2} + O(\sigma^4) \tag{17}$$

Furthermore, the variance of the response is given as (see Lemma S24):

$$var(\theta) = \tau^2 + \frac{1}{\mathcal{J}(\theta)} + O(\sigma^4) \tag{18}$$

or equivalently

$$var(\theta) = \tau^2 + \frac{\sigma^2}{F'(\theta)^2} + O(\sigma^4) \tag{19}$$

where $\tau^2$ is the variance of motor noise.

#### S2.3 Confoundedness in Low-Noise Regime

Here, we show that models can be confounded in the low-noise regime when their priors are linked according to a specific transformation, using the results from Section S2.2. Recall

$$\tilde{p} := \begin{cases} p & \text{if } p > 0 \\ -1 & \text{if } p = 0 \end{cases} \tag{20}$$

Then formally:

**Theorem S9.** *Let*

$$M_1 := \langle F_1', p_{prior}^{(1)}, p_1 \rangle$$
$$M_2 := \langle F_2', p_{prior}^{(2)}, p_2 \rangle$$

*be two models in $\mathfrak{M}$. In the absence of motor noise ($\tau = 0$), the following are equivalent:*

1. *$F_1 = F_2$, with identical Fisher Information $\mathcal{J} = \frac{F_1^2}{\sigma^2} = \frac{F_2^2}{\sigma^2}$, and the priors are linked as follows:*

$$p_{prior}^{(1)} \propto p_{prior}^{(2)} \mathcal{J}^{\frac{\tilde{p}_1 - \tilde{p}_2}{4}}, \tag{21}$$

2. *The response bias and variance of $M_1$ and $M_2$ differ only by $O(\sigma^4)$ when $\sigma$ is small.*

The second property intuitively indicates that, when $\sigma$ is small, $M_1$ and $M_2$ are essentially indistinguishable, as bias and variance scale wirh $\sigma^2$, which is much larger than $\sigma^4$ when $\sigma$ is small. We note that, in the notation used in Section S2.6, this condition amounts to stating that $\mathcal{A}_{M_1} \equiv \mathcal{A}_{M_2}$ and $\mathcal{C}_{M_1} \equiv \mathcal{C}_{M_2}$. We illustrate the theorem with a family of models satisfying this property in Figure S12.

*Proof.* We first show $1 \Rightarrow 2$. By (19), the two models generate the same variability up to $\sigma^4$ if $F_1 = F_2 =: F$. Further, based on (17), they generate the biases:

$$\begin{aligned}
bias_1(\theta) =& \frac{\sigma^2}{F'^2}\frac{d}{d\theta}\log p^{(1)}_{prior} + \frac{\tilde{p}_1+2}{4}\frac{d}{d\theta}\frac{\sigma^2}{F'^2} + O(\sigma^4)\\
=& \frac{\sigma^2}{F'^2}\frac{d}{d\theta}\log\left(p^{(2)}_{prior}\mathcal{J}^{\frac{\tilde{p}_1-\tilde{p}_2}{4}}\right) + \frac{\tilde{p}_1+2}{4}\frac{d}{d\theta}\frac{\sigma^2}{F'^2} + O(\sigma^4)\\
=& \frac{\sigma^2}{F'^2}\frac{d}{d\theta}\log\left(p^{(2)}_{prior}\right) + \frac{\tilde{p}_1-\tilde{p}_2}{2}\frac{\sigma^2}{F'(\theta)^2}\frac{d}{d\theta}\frac{F''(\theta)}{F'(\theta)} + \frac{\tilde{p}_1+2}{4}\frac{d}{d\theta}\frac{\sigma^2}{F'^2} + O(\sigma^4)\\
=& \frac{\sigma^2}{F'^2}\frac{d}{d\theta}\log p^{(2)}_{prior} + \sigma^2\frac{\tilde{p}_1-\tilde{p}_2}{2}\frac{F''(\theta)}{F'(\theta)^3} - \sigma^2\frac{\tilde{p}_1+2}{2}\frac{F''(\theta)}{F'(\theta)^3} + O(\sigma^4)\\
=& \frac{\sigma^2}{F'^2}\frac{d}{d\theta}\log p^{(2)}_{prior} + \frac{\tilde{p}_2+2}{4}\frac{d}{d\theta}\frac{\sigma^2}{F'^2} + O(\sigma^4)\\
=& bias_2(\theta) + O(\sigma^4)
\end{aligned}$$

To show $2 \Rightarrow 1$, we first note that the models can only generate the same variability if $F_1 = F_2$, again by (19). Second, retracing the computation of the bias above shows that the only way for $bias_1(\theta)$ and $bias_2(\theta)$ to differ only by $O(\sigma^4)$ is if the priors satisfy the specified transformation. □

#### S2.4 Proof of Theorem 1

**Theorem S10** (Recovering Encoding from Response Distribution)**.** *In the absence of motor noise, the encoding F and the sensory noise magnitude can be recovered from the response distribution at a single level of sensory noise, with estimation error exponentially small in* $\frac{1}{\sigma}$.

$$\sqrt{\mathcal{J}(\theta)} = \sqrt{4\pi}\cdot\frac{\mathrm{d}}{\mathrm{d}\theta}\mathbb{P}(\widehat{\theta}(\theta+h) \geq \widehat{\theta}(\theta)) + O(\exp(-1/\sigma))$$

We exemplify the application to a simulated dataset in Figure S3.

*Proof.* Let $\widehat{\theta}(\theta)$ be the response based on stimulus $\theta$ as a random variable, with randomness coming from the sensory encoding noise. That is,

$$\widehat{\theta}(\theta) = f(F^{-1}(F(\theta)+\varepsilon)) \tag{22}$$

where $f$ is the Decoding Function (Definition S11) and $\varepsilon \sim N(0,\sigma^2)$. We consider the observable quantity

$$\mathbb{P}\left(\widehat{\theta}(\theta_1) \geq \widehat{\theta}(\theta_2)\right) \tag{23}$$

where sensory noise is applied independently to the two encodings (as is assumed the case when these observations come from different trials). This, in the absence of motor noise, equals

$$\mathbb{P}\left(F(\theta_1)+\varepsilon_1 \geq F(\theta_2)+\varepsilon_2\right) \tag{24}$$

due to the monotonicity of both $F$ and the decoding function (Lemma S12), where $\varepsilon_1, \varepsilon_2$ describe independent sensory noise. We first consider the setting where $\mathcal{X}$ is the entire real line $\mathbb{R}$, so that $\varepsilon_1, \varepsilon_2$ are unconstrained Gaussians, $\varepsilon_1, \varepsilon_2 \sim N(0,\sigma^2)$. Then, the above equals

$$\mathbb{P}\left(\widehat{\theta}(\theta_1) \geq \widehat{\theta}(\theta_2)\right) = \mathbb{P}\left(F(\theta_1)-F(\theta_2) > \varepsilon_2-\varepsilon_1\right) = \Phi\left(\frac{F(\theta_1)-F(\theta_2)}{\sqrt{2\sigma^2}}\right) \tag{25}$$

where $\varepsilon_2 - \varepsilon_1 \sim N(0, 2\sigma^2)$, and $\Phi$ is the cumulative density function of the normal distribution. Now taking

$$\begin{aligned}\frac{d}{d\theta}\mathbb{P}(\widehat{\theta}(\theta+h) \geq \widehat{\theta}(\theta)) =& \frac{d}{d\theta}\mathbb{P}(F(\theta+h) - F(\theta) \geq \varepsilon_2 - \varepsilon_1) \\ =& \frac{d}{d\theta}\Phi\left(\frac{F(\theta+h)-F(\theta)}{\sqrt{2\sigma^2}}\right) \\ =& F'(\theta)\frac{1}{\sqrt{2\pi}}\frac{1}{\sqrt{2\sigma^2}}\end{aligned}$$

where the last step follows since $\Phi'(0) = \frac{1}{\sqrt{2\pi}}$. Integrating this and normalizing the sensory space volume then allows us to obtain the noise magnitude and the encoding separately. We now consider the more general case where $\mathcal{X}$ is a bounded interval or a circle. In this case, the density function of $\sqrt{2\sigma^2}(\varepsilon_2 - \varepsilon_1)$ at 0 is still equal to a standard Gaussian up to error exponentially small in $\frac{1}{\sigma}$. $\square$

We now define and discuss the Decoding Function $f$ used in the proof above:

**Definition S11** (Decoding Function)**.** *Given a model $M \in \mathfrak{M}$ and a sensory noise level $\sigma^2 > 0$, the* decoding function *$f : \mathcal{X} \to \mathcal{X}$ maps a point $\theta$ to the result of decoding $\widehat{\theta}$ from the encoding $F(\theta)$. It is defined by minimization of the loss function:*

$$f(\theta) := \arg_{\widehat{\theta}}\min \int_{\mathcal{X}} \ell(\widehat{\theta}, x) P_{post}(x|m = F(\theta))dx \tag{26}$$

We establish an important property of the decoding function, key to establishing Theorem 1:

**Lemma S12.** *Let $f$ be the Decoding Function. If $\mathcal{X}$ is an interval, then $f$ is nondecreasing.*

*Proof.* We start with the case where $\ell$ represents $L_p$ loss for $p \geq 1$. Let $P_{lik}(m|x)$ ($x \in \mathcal{X}$, $m \in \mathcal{Y}$) be the likelihood expressed in the sensory space. Let $P_{post}(x|m)$ ($x \in \mathcal{X}$, $m \in \mathcal{Y}$) be the posterior. By Bayes' Rule,

$$P_{post}(x|m = F(\theta)) = \frac{P_{prior}(x)P_{lik}(F(\theta)|x)}{\int P_{lik}(F(\theta)|s)ds} \tag{27}$$

where the denominator is independent of $x$. We write

$$f(\theta) := \arg_{\widehat{\theta}}\min \int_{\mathcal{X}} \ell(\widehat{\theta}, x) P_{post}(x|m = F(\theta))dx \tag{28}$$

We may set

$$\mathcal{L}(\widehat{\theta}, \theta) = -\int_{\mathcal{X}} \ell(\widehat{\theta}, x) P_{post}(x|F(\theta))dx \tag{29}$$

The monotonicity properties of such solutions to optimization problems are studied in the field of Comparative Statics [25]. An early result of this type for Bayesian estimators is Theorem 3 in Karlin and Rubin [8]. Here, we provide a simple and self-contained proof of the lemma grounded in Topkis' Theorem [24, 25, 11], which states that $f$ is nondecreasing provided $\mathcal{L}$ is *supermodular*, which means that, whenever $h_1 \leq h_2$ and $\theta_1 \leq \theta_2$, we have

$$\mathcal{L}(h_2, \theta_2) - \mathcal{L}(h_1, \theta_2) \geq \mathcal{L}(h_2, \theta_1) - \mathcal{L}(h_1, \theta_1) \tag{30}$$

Fixing any two $h_1 \leq h_2$, define

$$D(x) = \ell(h_1, x) - \ell(h_2, x) \tag{31}$$

which is nondecreasing in $x$ for $p \geq 1$. For $\theta_2 > \theta_1$,

$$\mathcal{L}(h_2, \theta_2) - \mathcal{L}(h_1, \theta_2) = \int D(x) P_{post}(x|\theta_2)dx \geq^{(\dagger)} \int D(x) P_{post}(x|\theta_1)dx = \mathcal{L}(h_2, \theta_1) - \mathcal{L}(h_1, \theta_1)$$

proving that $\mathcal{L}$ is supermodular. It remains to justify $(\dagger)$. We first examine

$$\frac{P_{post}(x|\theta_2)}{P_{post}(x|\theta_1)} = \frac{P_{lik}(F(\theta_2)|x)}{P_{lik}(F(\theta_1)|x)} \frac{\int P_{lik}(F(\theta_2)|s)ds}{\int P_{lik}(F(\theta_1)|s)ds} \tag{32}$$

The second multiplicand is independent of $x$. The first multiplicand is increasing in $x$:

$$\log \frac{P_{lik}(F(\theta_2)|x)}{P_{lik}(F(\theta_1)|x)} = \frac{(F(\theta_1)-x)^2-(F(\theta_2)-x)^2}{2t} = \frac{1}{2t}\left(F(\theta_1)^2 - F(\theta_2)^2 + 2x \cdot (F(\theta_2)-F(\theta_1))\right)$$

which is increasing in $x$ because $F(\theta_2) > F(\theta_1)$. Hence, the posterior exhibits monotone likelihood ratios, in the sense that, for each $\theta_2 > \theta_1$,

$$\frac{P_{post}(x|\theta_2)}{P_{post}(x|\theta_1)} \text{ is increasing in } x \tag{33}$$

This property entails stochastic dominance (e.g., Theorem 1.C.1 in [20]; Theorem 1 in [9]): That is, if $\Psi_1, \Psi_2$ are the cumulative distribution functions of the two posteriors $P_{post}(x|\theta_1)$, $P_{post}(x|\theta_2)$, then we have:

$$\Psi_1(x) \geq \Psi_2(x), \quad \forall x \tag{34}$$

Because $D$ is nondecreasing, (†) follows using an equivalent characterization of stochastic dominance in terms of integrals of nondecreasing functions (e.g., Eq. 1.A.7 in [20]):

$$\int D(x)P_{post}(x|\theta_2)dx = \int D(X)d\Psi_2(X) \geq \int D(X)d\Psi_1(X) = \int D(x)P_{post}(x|\theta_1)dx$$

This concludes the proof when $p \geq 1$. Now at $p = 0$, we instead apply Topkis' theorem to the log of the unnormalized posterior:

$$\mathcal{L}(\widehat{\theta}, \theta) = \log p_{prior}(\widehat{\theta}) p_{lik}(F(\theta)|\widehat{\theta}) \tag{35}$$

where $f(\theta) = \arg\max_{\widehat{\theta}} \mathcal{L}(\widehat{\theta}, \theta)$. As above, we need to show that $\mathcal{L}$ is supermodular; this is now easy due to Topkis' characterization that $\mathcal{L}$ is supermodular if and only if $\partial_{\widehat{\theta}}\partial_\theta \mathcal{L} \geq 0$ everywhere [24, 25]. We check:

$$\begin{aligned}
\partial_{\widehat{\theta}}\partial_\theta \mathcal{L}(\widehat{\theta}, \theta) =& \partial_{\widehat{\theta}}\partial_\theta \left[\log p_{prior}(\widehat{\theta}) p_{lik}(F(\theta)|\widehat{\theta})\right] \\
=& \partial_{\widehat{\theta}}\partial_\theta \left[\log p_{lik}(F(\theta)|\widehat{\theta})\right] \\
=& -\frac{1}{2t}\partial_{\widehat{\theta}}\partial_\theta \left[(F(\theta)-F(\widehat{\theta}))^2\right] \\
=& -\frac{1}{t}\partial_{\widehat{\theta}} \left[(F(\theta)-F(\widehat{\theta}))F'(\theta)\right] \\
=& \frac{1}{t}F'(\widehat{\theta})F'(\theta)
\end{aligned}$$

which is positive.

□

#### S2.5 Proof of Theorem 2

Let $\mathcal{P}$ be the (finite) set of admissible loss function exponents (e.g., $0, 1, 2, 4, 6, 8, 10$). As discussed in Section S2.2, it is convenient to represent the MAP estimator using $-1$ instead of $0$ as a convention (so the expression for the bias matches the pattern for the other exponents). Formally,

$$\tilde{p} := \begin{cases} p & \text{if } p > 0 \\ -1 & \text{if } p = 0 \end{cases} \tag{36}$$

We now prove Theorem 2:

**Theorem S13** (Restated from Theorem 2: One Level of Sensory Noise)**.** *Let* $\sigma > 0$ *be the SD of sensory noise, and* $p \in \mathcal{P}$*. Assume the response distribution* $P(\widehat{\theta}|M,\theta,\sigma)$ *is given at each* $\theta$*, under a ground-truth model* $\langle F', p_{prior}, p\rangle \in \mathfrak{M}$*. There is a function* $\Phi_p$ *mapping these distributions to* $\langle \widehat{F}, \widehat{p_{prior}}, p\rangle \in \mathfrak{M}$ *such that*

$$|\widehat{F}(\theta) - F(\theta)| = O(\sigma^2)$$
$$|\widehat{p_{prior}}(\theta) - p_{prior}(\theta)| = O(\sigma^2)$$

*as* $\sigma \to 0$*. Constants in* $O(\cdot)$ *depend on* $F$ *and* $p_{prior}$*, but not* $\theta$*.*

We prove a slightly more general statement that includes an estimate of the (potentially incorrectly) estimated prior when the loss function might be potentially misspecified; Theorem 2 follows in the special case when the loss functions match:

**Theorem S14** (Generalization of Theorem 2)**.** *Let* $\sigma > 0$ *be the SD of sensory noise. Assume the response distribution* $P(\widehat{\theta}|M,\theta,\sigma)$ *is given at each* $\theta$*, under a ground-truth model* $\langle F', p_{prior}, p\rangle \in \mathfrak{M}$*. For each exponent* $q \geq 0$*, there is a function* $\Phi_q$ *mapping these distributions to* $\langle \widehat{F}, \widehat{p_{prior}}, q\rangle \in \mathfrak{M}$ *such that*

$$|\widehat{F}(\theta) - F(\theta)| = O(\sigma^2)$$
$$|\widehat{p_{prior}}(\theta) - U \cdot (F')^{\frac{\tilde{q}-\tilde{p}}{2}} \cdot p_{prior}(\theta)| = O(\sigma^2)$$

*as* $\sigma \to 0$*, for some constant*

$$U = \frac{1}{\int (F'(\theta))^{\frac{q-p}{2}} \cdot p_{prior}(\theta) d\theta} \tag{37}$$

*The identified model has bias and variance equal to b and v up to error* $O(\sigma^2)$*. Constants in* $O(\cdot)$ *depend on* $F$ *and* $p_{prior}$*, but not* $\theta$*.*

We note that Theorem 2 follows in the special case where $p = q$.

*Proof of Theorem S14.* We obtain the encoding $\widehat{F}$ using Lemma S10.[7] We define $\Phi_q$ by setting

$$\mathcal{E}(\theta) := \frac{(bias(\theta) - \frac{q+2}{4} v'(\theta))}{v(\theta)}$$
$$\widehat{p_{prior}}(\theta) \propto \exp\left(\int_0^\theta \mathcal{E}(\theta') d\theta'\right)$$

Since the fist two moments $bias(\theta)$ (bias) and $var(\theta)$ (variance) of the response distribution $P(\widehat{\theta}|M,\theta,\sigma)$ are given at each $\theta$, and the sensory noise obeys Gaussian distribution $\mathcal{N}(0,\sigma^2)$, we have (Section S2.2 and Lemma S23):

$$var(\theta) = \mathcal{A}(\theta)\sigma^2 + \mathcal{B}(\theta)\sigma^4 + O(\sigma^6)$$
$$bias(\theta) = \mathcal{C}(\theta)\sigma^2 + \mathcal{D}(\theta)\sigma^4 + O(\sigma^6)$$

where $\mathcal{A}(\theta) = \frac{1}{F'(\theta)^2}$.We will write $t$ for the sensory noise variance $\sigma^2$ to make the notation easier.

Then we prove $\widehat{p_{prior}}(\theta) = U \cdot (F')^{\frac{q-p}{2}} \cdot p_{prior}(\theta) + O(t)$. According to the definition of $\mathcal{E}(\theta), var(\theta), bias(\theta)$, we have:

$$\mathcal{E}(\theta) = \frac{(bias(\theta) - \frac{\tilde{q}+2}{4} v'(\theta))}{var(\theta)} = \frac{[\mathcal{C}(\theta)t + O(t^2)] - \frac{\tilde{q}+2}{4}[\mathcal{A}'(\theta) + O(t^2)]}{\mathcal{A}(\theta)t + O(t^2)} = \frac{\mathcal{C}(\theta)t - \frac{\tilde{q}+2}{4}\mathcal{A}'(\theta)t + O(t^2)}{\mathcal{A}(\theta)t + O(t^2)} = \frac{\mathcal{C}(\theta) - \frac{\tilde{q}+2}{4}\mathcal{A}'(\theta)}{\mathcal{A}(\theta)} + O(t)$$

[7] An alternative approach, based on only moments but with error $O(t)$, could proceed as in the proof of Theorem 3.

As discussed in Section S2.2, we have $\mathcal{C}(\theta) = \frac{(\log p_{prior})'}{F'^2} - \frac{p+2}{2} \cdot \frac{F''}{F'^3}$. Thus, we can plug in the expression of $\mathcal{C}(\theta)$ into the definition of $\widehat{p_{prior}}(\theta)$ and let $\alpha$ be the constant coefficient. Then we have:

$$
\begin{aligned}
\widehat{p_{prior}}(\theta) &= \alpha \cdot \exp\left(\int_0^\theta \mathcal{E}(\theta')d\theta'\right) \\
&= \alpha \cdot exp\left(\int_0^\theta \frac{C(\theta') - \frac{\tilde{q}+2}{4}\mathcal{A}'(\theta)}{\mathcal{A}(\theta')} d\theta' + O(t)\right) \\
&= \alpha \cdot \exp\left(\int_0^\theta [(\log p_{prior})' + \frac{\tilde{q}-\tilde{p}}{2}\frac{F''}{F'}]d\theta' + O(t)\right) \\
&= \alpha \cdot \exp\left(\int_0^\theta [\log p_{prior} + \frac{\tilde{q}-\tilde{p}}{2}\log F']' d\theta' + O(t)\right) \\
&= \alpha \cdot \exp\left(\int_0^\theta \left(\log\left(p_{prior} \cdot F'^{\frac{\tilde{q}-\tilde{p}}{2}}\right)\right)' d\theta' + O(t)\right) \\
&= \alpha \cdot \exp\left(\log\left(p_{prior}(\theta) \cdot F'(\theta)^{\frac{\tilde{q}-\tilde{p}}{2}}\right) - \log\left(p_{prior}(0) \cdot F'(0)^{\frac{\tilde{q}-\tilde{p}}{2}}\right) + O(t)\right) \\
&= \alpha \cdot \frac{p_{prior}(\theta) \cdot F'(\theta)^{\frac{\tilde{q}-\tilde{p}}{2}}}{p_{prior}(0) \cdot F'(0)^{\frac{\tilde{q}-\tilde{p}}{2}}} \cdot e^{O(t)} \\
&= \alpha \cdot \frac{p_{prior}(\theta) \cdot F'(\theta)^{\frac{\tilde{q}-\tilde{p}}{2}}}{p_{prior}(0) \cdot F'(0)^{\frac{\tilde{q}-\tilde{p}}{2}}} \cdot (1 + O(t)) \\
&= \alpha \cdot \frac{p_{prior}(\theta) \cdot F'(\theta)^{\frac{\tilde{q}-\tilde{p}}{2}}}{p_{prior}(0) \cdot F'(0)^{\frac{\tilde{q}-\tilde{p}}{2}}} + O(t)
\end{aligned}
$$

where

$$
\begin{aligned}
&\alpha \cdot \frac{1}{p_{prior}(0) \cdot F'(0)^{\frac{\tilde{q}-\tilde{p}}{2}}} \cdot \int_0^{\theta_{max}} p_{prior}(\theta') \cdot F'(\theta')^{\frac{\tilde{q}-\tilde{p}}{2}} d\theta' = 1 \\
&\alpha = \frac{p_{prior}(0) \cdot F'(0)^{\frac{\tilde{q}-\tilde{p}}{2}}}{\int_0^{\theta_{max}} p_{prior}(\theta') \cdot F'(\theta')^{\frac{\tilde{q}-\tilde{p}}{2}} d\theta'}
\end{aligned}
$$

Thus, $\widehat{p_{prior}}(\theta) = U \cdot (F')^{\frac{\tilde{q}-\tilde{p}}{2}} \cdot p_{prior}(\theta) + O(\sigma^2)$ is proved.

□

#### S2.6 Proof of Theorems 3 and 4

Given a model $M \in \mathfrak{M}$, we consider the Taylor expansion of the variability and bias of the response, given a stimulus $\theta$, in terms of the squared magnitude of sensory noise shown in Lemma S23, and refining the decomposition reviewed in Section S2.2:[8]

$$
\begin{aligned}
var(\theta) &= \tau^2 + \mathcal{A}_M(\theta)\sigma^2 + \mathcal{B}_M(\theta)\sigma^4 + O(\sigma^6) \\
bias(\theta) &= \mathcal{C}_M(\theta)\sigma^2 + \mathcal{D}_M(\theta)\sigma^4 + O(\sigma^6)
\end{aligned}
$$

[8]We justify in Section S2.6.3 that these coefficients exist, that is, that the bias is twice differentiable in $\sigma^2$ at the limit point $\sigma^2 = 0$ when $F$ and $p_{prior}$ are sufficiently regular.

where $\tau^2$ is the motor noise variance, and $\sigma^2$ is the sensory noise variance. This refines the decomposition reviewed in Section S2.2, by setting

$$\mathcal{A}_M(\theta) := \frac{1}{F'^2}\frac{d}{d\theta}\log p_{prior}(\theta) + \frac{\tilde{p}+2}{4}\frac{d}{d\theta}\frac{1}{F'(\theta)^2}$$
$$\mathcal{C}_M(\theta) := \frac{1}{F'(\theta)^2}$$

See Section S2.6.3 for explicit expressions for $\mathcal{B}_M, \mathcal{D}_M$.

##### S2.6.1 Definition of Exceptional Set $\Omega$

We define $\Omega$ in terms of models $M_1$ where there is some model $M_2$ distinct from it such that the coefficients $\mathcal{A}_M, \mathcal{B}_M, \mathcal{C}_M, \mathcal{D}_M$ are identical for $M_1$ and $M_2$ – that is, where the response bias and variance are identical up to a higher-order residual of order $O(\sigma^6)$. We also include models where the first-order bias component $\mathcal{C}_M$ vanishes entirely, i.e., where the bias is zero everywhere in the low-noise regime.[9] Then:

**Definition S15.** *We define $\Omega \subset \mathfrak{M}$ as:*

$$\Omega := \{M : \mathcal{C}_M \equiv 0\} \cup \bigcup_{F, p_{prior}} \bigcup_{p,p' \in \mathcal{P}} \{M_1 \in \mathfrak{M} : \exists M_2 \in \mathfrak{M} : M_2 \neq M_1; \mathcal{A}_{M_1} \equiv \mathcal{A}_{M_2}; \mathcal{B}_{M_1} \equiv \mathcal{B}_{M_2}; \mathcal{C}_{M_1} \equiv \mathcal{C}_{M_2}; \mathcal{D}_{M_1} \equiv \mathcal{D}_{M_2}\}$$

It will be useful to define $\Omega$ more explicitly in terms of (i) identical encoding, (ii) prior transformed based on the loss function difference, (iii) $\mathcal{D}_M$ identical for both models. Recall the definition of $\tilde{p}$ in Eq. 36.

**Lemma S16.** *We have:*

$$\Omega = \{M : \mathcal{C}_M \equiv 0\} \cup \bigcup_{F, p_{prior}} \bigcup_{p,p' \in \mathcal{P}} \{M_1 := \langle F', p_{prior}, p\rangle : \exists p' \neq p, C \in \mathbb{R}, \mathcal{D}_{M_1} \equiv \mathcal{D}_{M_2} \text{ where } M_2 := \langle F', C \cdot (F')^{\frac{\tilde{p}'-\tilde{p}}{2}} p_{prior}, p'\rangle\}$$

*Proof.* First, when $\mathcal{A}_{M_1} \equiv \mathcal{A}_{M_2}$ and $\mathcal{C}_{M_1} \equiv \mathcal{C}_{M_2}$, then the form $M_1 = \langle F', p_{prior}, p\rangle$, $M_2 = \langle F', C \cdot (F')^{\frac{\tilde{p}'-\tilde{p}}{2}} p_{prior}, p'\rangle$ follows from the expressions for $\mathcal{A}$ and $\mathcal{C}$ (Lemma S24). It remains to prove the other direction, i.e., that when $M_2 := \langle F', C \cdot (F')^{\frac{\tilde{p}'-\tilde{p}}{2}} p_{prior}, p'\rangle$, then $\mathcal{A}_{M_1} \equiv \mathcal{A}_{M_2}, \mathcal{B}_{M_1} \equiv \mathcal{B}_{M_2}, \mathcal{C}_{M_1} \equiv \mathcal{C}_{M_2}$ follow. Since the encoding functions $F$ of $M_1$ and $M_2$ are identical, $\mathcal{A}_{M_1} \equiv \mathcal{A}_{M_2}$ holds obviously ($\mathcal{A}$ only depends on $F$). Now, $\mathcal{C}_{M_1} \equiv \mathcal{C}_{M_2}$ follows from the expression ():

$$\mathcal{C}_M(\theta) = \frac{1}{F'^2}\frac{d}{d\theta}\log p_{prior}(\theta) + \frac{\tilde{p}+2}{4}\frac{d}{d\theta}\frac{1}{F'(\theta)^2} \tag{38}$$

or equivalently

$$\mathcal{C}_M(\theta) = \frac{1}{F'^2}\frac{d}{d\theta}\log p_{prior}(\theta) - \frac{\tilde{p}+2}{2}\frac{F''(\theta)}{F'(\theta)^3}$$

Next we prove that $\mathcal{B}_{M_1} \equiv \mathcal{B}_{M_2}$. According to Lemma S24, we have

$$\mathcal{B}_M(\theta) = \frac{2g(\theta)}{F'^2(\theta)} + \frac{F''(\theta)}{2F'^6(\theta)}$$

where $g_M = \frac{d}{d\theta}\mathcal{C}_{dec,M}$. Since $\mathcal{C}_{M_1} = \mathcal{C}_{M_2}$, we also have $\mathcal{C}_{dec,M_1} = \mathcal{C}_{dec,M_2}$. Hence, it follows that $\mathcal{B}_{M_1} = \mathcal{B}_{M_2}$. Overall, we have shown the equivalence of the two expressions for $\Omega$. □

[9]This is for technical reasons, as the proof of Lemma S22 presupposes that $\mathcal{C}$ is nonzero for some stimuli. In practice, as soon as prior or encoding are nonuniform, the only way for $\mathcal{C}_M$ to vanish everywhere is for $F$ and $p_{prior}$ to be aligned in a very specific way to ensure perfect cancellation of attraction and repulsion.

We conclude:

**Corollary S17.** *$\Omega$ has volume zero under the measure $\mu$: $\mu(\Omega) = 0$.*

*Proof of Lemma S17.* We condition on $F$ and $p \in \mathcal{P}$, and show that the set of $L(p_{prior})$ such that $\langle F', p_{prior}, p\rangle \in \Omega$ has measure zero; the result then follows from the Fubini-Tonelli theorem. We first note that, given $F$ and $p$, the condition $\mathcal{C}_M \equiv 0$ directly defines $L(p_{prior})$, leading to zero measure. Now we consider the second component: For each $p' \neq p$, we investigate the condition

$$\mathcal{D}_{M_1} = \mathcal{D}_{M_2} \tag{39}$$

where

$$M_2 := \langle F', U \cdot (F')^{\frac{p'-p}{2}} p_{prior}, p\rangle \tag{40}$$

Considering the form of $\mathcal{D}$ (Equation 64), the highest derivatives contribute a single term at $p, p' \geq 2$, and – after simplification – the condition $\mathcal{D}_{M_1} = \mathcal{D}_{M_2}$ can be written as:

$$\frac{d^3}{d\theta^3} q_{prior}(m) = f\left((q_{prior})(m), \frac{d}{d\theta} q_{prior}(m), \frac{d^2}{d\theta^2} q_{prior}(m)\right) \tag{41}$$

for some continuous function $f$ (depending on $F$, $p$, $p'$). By the Picard-Lindelöf theorem, any initial value for $q_{prior}$ and its first three derivatives at $m = 0$ uniquely determines the corresponding values at sufficiently small $m > 0$. Thus, the set of $L(p_{prior})$ solving the equation $\mathcal{D}_{M_1} = \mathcal{D}_{M_2}$ has measure zero in $\mathcal{F}$ by Assumption S7. The union over different $p' \neq p$ still has zero measure. The proof is similar if $p$ or $p' < 2$.

□

##### S2.6.2 Proof of Theorem

We jointly prove Theorems 3 and 4. Recall that Theorem 4 extends Theorem 3 by allowing nonzero motor noise. For reference, we first restate Theorem 3:

**Theorem S18** (Restated from Theorem 3)**.** *There is a subset $\Omega \subset \mathfrak{M}$ of volume 0 such that the following holds. Let $0 < \sigma_1 < \sigma_2$ be two levels of sensory noise. Assume the fist two moments of the response distribution $\mathbb{P}(\widehat{\theta}|M, \theta, \sigma_i)$ are given at each $\theta$ and for both $\sigma_1, \sigma_2$, under a ground-truth model $M = \langle F', p_{prior}, p\rangle \in \mathfrak{M}$. There is a functional $\Phi$ mapping the collection of these distributions to $\langle \widehat{F}', \widehat{p_{prior}}, q\rangle \in \mathfrak{M}$ such that—provided $M \notin \Omega$—we have*

$$\widehat{F}(\theta) = F(\theta) + O(\sigma_1^2)$$
$$\widehat{p_{prior}}(\theta) = p_{prior}(\theta) + O(\sigma_1^2)$$
$$q = p \text{ for } \sigma_1^2 \text{ small}$$

*in the limit where $\sigma_1, \sigma_2 \to 0$, provided:*

$$0 < C_1 < \frac{\sigma_1}{\sigma_2} < C_2 < 1 \tag{42}$$

*for some $C_1, C_2$. Constants in the $O(\cdot)$ expressions depend on $C_1, C_2$, and on $F$ and $p_{prior}$, but not on $\theta$.*

and Theorem 4:

**Theorem S19.** *[Restated from Theorem 4] Theorem S18 is not affected by adding symmetric isotropic motor noise of variance $\tau^2$, and by guessed responses appearing at a rate $0 < \gamma < 1$, in the limit $\tau, \sigma_1, \sigma_2 \to 0$, while maintaining $D_1 < \frac{\sigma_1}{\tau} < D_2$ for some $0 < D_1 < D_2 < \infty$.*

Our proof applies directly to the overall result covering both Theorems 3 and 4:

**Theorem S20** (Restated from Theorems 3 and 4). *Consider the extended model with Gaussian motor noise and guessing. There is a subset $\Omega \subset \mathfrak{M}$ of volume 0 such that the following holds. Let $0 < \sigma_1 < \sigma_2$ be two levels of sensory noise. Let $\tau^2$ be the variance of motor noise. Assume the fist two moments of the response distribution $\mathbb{P}(\widehat{\theta}|M,\theta,\sigma_i)$ are given at each $\theta$ and for both $\sigma_1,\sigma_2$, under a ground-truth model $M = \langle F', p_{prior}, p\rangle \in \mathfrak{M}$. There is a functional $\Phi$ mapping the collection of these distributions to $\langle \widehat{F}', \widehat{p_{prior}}, q\rangle \in \mathfrak{M}$ such that—provided $M \notin \Omega$—we have*

$$\widehat{F}(\theta) = F(\theta) + O(\sigma_1^2)$$
$$\widehat{p_{prior}}(\theta) = p_{prior}(\theta) + O(\sigma_1^2)$$
$$q = p \text{ for } \sigma_1^2 \text{ small}$$

*in the limit where $\sigma_1, \sigma_2, \tau \to 0$, provided:*

$$1 < C_1 < \frac{\sigma_2}{\sigma_1} < C_2 < \infty$$
$$0 < C_3 < \frac{\tau}{\sigma_1} < C_4 < \infty$$

*for some $C_1,C_2,C_3,C_4$. Constants in the $O(\cdot)$ expressions depend on $C_1,C_2,C_3,C_4$, and on the regularity of $F$ and $p_{prior}$ and their derivatives, but not on $\theta$.*

We will use the following function norms:

**Definition S21.** *Let $f : X \to \mathbb{R}$ be a function, then*

$$\|f\|_\infty := \sup_{x\in X} |f(x)| \tag{43}$$

*Next, if $f$ is k-times differentiable, then we set*

$$\|f\|_{\infty,k} := \sum_{i=0}^{k} \|f^{(i)}\|_\infty \tag{44}$$

We proceed to proving the theorem, using Lemma S22:

*Proof of the theorem.* We first note that, in the limit of small sensory noise, the guessing rate $\gamma$ can be identified by investigating the rates at which stimuli $\theta$ elicit far-away responses, with an error exponentially small in $1/\sigma$. By deconvolving the response distributions, one can then obtain the raw response distributions before adding guesses; we can thus assume $\gamma = 0$ for the remainder of the proof. We define a parameter $t$ by setting:

$$\sigma_1^2 = at \qquad \sigma_2^2 = bt \qquad \tau^2 = \rho^2 t$$

where, to make this value definite, we take $a = 1$. By assumption, in the limit of interest,

$$t \to 0$$
$$0 < C_1 < \frac{b}{a} < C_2 < 1$$
$$0 < C_3 < \frac{\rho}{a} < C_4 < \infty$$

Now given the biases $b_1(\theta)$, $b_2(\theta)$ and variances $v_1(\theta)$, $v_2(\theta)$, we compute $\widehat{F}$, $\widehat{p_{prior}}_{p'}$, $\tilde{\mathcal{D}}_{p'}$ for each $p' \in \mathcal{P}$ as in Lemma S22, to form models

$$M_{p'} := \langle \widehat{F}', \widehat{p_{prior}}, p'\rangle \tag{45}$$

where $\widehat{p_{prior}}$ is fitted depending on $p'$. On the other hand, we define reference models

$$\hat{M}_{p'} := \langle F', U F'^{\frac{p'-p}{2}} p_{prior}, p'\rangle \tag{46}$$

(with $U$ a normalization constant to make the prior integrate to 1) where specifically $\hat{M}_p = M$. For each $p' \in \mathcal{P}$, we compute:

$$Err(p') := \|\tilde{\mathcal{D}}_{p'} - \mathcal{D}_{M_{p'}}\|_\infty =^{(\dagger)} \|\mathcal{D}_M - \mathcal{D}_{M_{p'}}\|_\infty + O(t)$$

where ($\dagger$) follows from $\|\tilde{\mathcal{D}}_{p'} - \mathcal{D}_M\|_\infty = O(t)$ (Lemma S22). Furthermore, due to the formula for $\mathcal{D}_M$ (Lemma S25) and the convergence guarantee for $\widehat{F}, \widehat{p_{prior}}$ given by Lemma S22, we have

$$\|\mathcal{D}_{M_{p'}} - \mathcal{D}_{\hat{M}_{p'}}\|_\infty = O(t) \tag{47}$$

Hence, when $p = p'$, then $Err_{p'} = O(t)$. On the other hand, consider $p' \neq p$. If the ground-truth model is not in $\Omega$, then $\mathcal{D}_{M_{p'}}$ converges in $\|\cdot\|_\infty$ to $\mathcal{D}_{\hat{M}_{p'}} \neq \mathcal{D}_M$ as $t \to 0$; hence, $Err(p') \not\to 0$ as $t \to 0$. Taken together, for sufficiently small $t$, and if the model is not in $\Omega$, $Err(p')$ will be minimized uniquely by the ground-truth exponent $p$. The result then follows. □

We are now ready to prove the main lemma used in the proof of Theorem 3:

**Lemma S22.** *Consider the model in the presence of motor noise $\varepsilon \sim N(0, \rho^2 t)$. Let $p' \in \mathcal{P}$. There is a function $\Phi_{p'}$ mapping biases and variances to a model $\langle \widehat{F}', \widehat{p_{prior}}, p' \rangle$, and a function $\tilde{\mathcal{D}} \in C(\mathcal{X})$ such that, given data from the model $\langle F', p_{prior}, p \rangle$ ($C_M(\theta)$ not constant zero), we have:*

1. $\|\widehat{F}(\theta) - F(\theta)\|_{\infty,4} = O(t)$
2. $\|\frac{d}{d\theta} \log \widehat{p_{prior}} - \frac{d}{d\theta} \log p_{prior}\|_{\infty,2}$ = *O(t)*
3. $\|\tilde{\mathcal{D}}(\theta) - \mathcal{D}(\theta)\|_\infty = O(t)$ *where $\mathcal{D}$ is the second-order component of the bias of the model $\langle F', p_{prior}, p \rangle$*

*in the limit $t \to 0$, with constants depending on $a, b, \rho$, and the regularity of $F$, $p_{prior}$, and their derivatives.*

*Proof.* In the absence of motor noise, the existence of estimates $\widehat{F}$, $\widehat{p_{prior}}$ with the error estimates claimed already follows from the proof of Theorem 2. The key challenge is to show that, with two levels of sensory noise, (i) these estimates continue to hold in the presence of motor noise, (ii) the estimate for $\tilde{\mathcal{D}}$ also holds.

If the observed human response has Gaussian motor noise, we have for the response $\tilde{\theta}$:

$$\begin{aligned} m &= F(\theta) + \delta, \delta \sim N(0, t) \\ \widehat{\theta} &= f(m) + \varepsilon, \varepsilon \sim N(0, \rho^2 t) \end{aligned}$$

($\delta$ and $\varepsilon$ independent) where $f : \mathcal{Y} \to \mathcal{X}$ is the function computing the $L_p$ estimator on the basis of a neural encoding $m$. The bias and variance of human responses in two different levels of sensory noise are given in terms of the decomposition given by Lemma S24:

$$\begin{aligned} var_1 &= a\mathcal{A}t + \rho^2 t + a^2 \mathcal{B}t^2 + O(t^3) \\ var_2 &= b\mathcal{A}t + \rho^2 t + b^2 \mathcal{B}t^2 + O(t^3) \\ bias_1 &= aCt + a^2 \mathcal{D}t^2 + O(t^3) \\ bias_2 &= bCt + b^2 \mathcal{D}t^2 + O(t^3) \end{aligned} \tag{48}$$

Here, the left hand side (bias and variability) is considered known, the terms on the right hand side are not directly observed. The basic idea of the proof is that we have a system with four equations that are linear in the four unknowns $\mathcal{A}, \mathcal{B}, C, \mathcal{D}$, hence one may hope to identify these quantities. As in Theorem 2, one obtains $F$ and (conditional on $p$) $p_{prior}$ from $\mathcal{A}$ and $C$; $\mathcal{D}$ is now also provided.

**Illustration: Proof without Motor Noise** For illustrative purposes, we first prove the theorem in the absence of motor noise. In this setup, we can use Lemma S10 to obtain $F$, $at$, $bt$ to very high precision. We can also obtain the prior $\widehat{p_{prior}}$ as in Theorem 2. We then set

$$\tilde{\mathcal{D}}(\theta) := \frac{bias_1(\theta) - \frac{at}{bt} bias_2(\theta)}{at \cdot (at - bt)} = \mathcal{D}(\theta) + O(t)$$

concluding the proof. The situation is more complex in the presence of motor noise, as we need to first separate out the effect of motor noise in estimating sensory noise. The full proof approaches this challenge by first obtaining an estimate of $\frac{a}{b}$, $at$, $bt$ that control for the effect of motor noise except perhaps a higher-order residual.

**Full Proof with Motor Noise** We now proceed with the proof in the more general case of nonzero motor noise. We first estimate several quantities related to the noise magnitudes. First, we note, for any $\theta \in \mathcal{X}$,

$$(b-a)\mathcal{A}(\theta)t = var_2(\theta) - var_1(\theta) + O(t^2)$$
$$b\mathcal{A}t = \frac{var_2(\theta) - var_1(\theta) + O(t^2)}{1 - \frac{a}{b}}$$

We now define a first, coarse, estimator of $\frac{a}{b}$ based on the observed bias and variability. Let $\theta^* \in \mathcal{X}$ be such that

$$|bias_1(\theta^*)| = \max_\theta |bias_1(\theta)| \tag{49}$$

Under the assumption that $\mathcal{C}(\theta)$ is nonzero at least for some $\theta$, then, when $t$ is small, $\theta^*$ will satisfy $\mathcal{C}(\theta^*) \neq 0$. Then we define an estimate of $\frac{a}{b}$ based on the observed quantities:

$$\widehat{\left(\frac{a}{b}\right)} := \frac{bias_1(\theta^*)}{bias_2(\theta^*)} = \frac{a \cdot \mathcal{C}(\theta^*)t + O(t^2)}{b \cdot \mathcal{C}(\theta^*)t + O(t^2)} = \frac{a}{b} + O(t)$$

Based on this, we define:

$$\widehat{b\mathcal{A}t} := \frac{var_2 - var_1}{1 - \widehat{\frac{a}{b}}} = \frac{bias_2 \cdot var_2 - bias_2 \cdot var_1}{bias_2 - bias_1} = \frac{var_2 - var_1}{1 - \frac{a}{b} + O(t)} = b\mathcal{A}t + O(t^2)$$
$$\widehat{\rho^2 t} := var_2 - \widehat{b\mathcal{A}t} = \frac{bias_2 \cdot var_1 - bias_1 \cdot var_2}{bias_2 - bias_1} = \rho^2 t + O(t^2)$$

A first approach to concluding the proof might be to substitute $t' := bt$, insert $\widehat{(\frac{a}{b})}$ instead of $a$ and $\widehat{\rho^2 t}$ instead of $\rho^2 t$ into (48), so that this system of equations is now linear in the four remaining unknowns $\mathcal{A}t', \mathcal{B}t', \mathcal{C}t'^2, \mathcal{D}t'^2$. A linear system of four equations in four unknowns should now admit a unique solution, allowing us to extract $\mathcal{A}, \mathcal{B}, \mathcal{C}, \mathcal{D}$ up to the scalar parameter $t'$. While on the right track, this idea requires refinement. The reason is that the estimates $\widehat{\rho^2 t}$ and $\widehat{(\frac{a}{b})}$ have an estimation error that, upon inserting these quantities instead of $\rho^2 t$ and $a$ would uncontrollably contaminate the terms at order $t^2$, preventing extraction of $\mathcal{C}$ and $\mathcal{D}$. We solve this puzzle by first refining $\widehat{(\frac{a}{b})}$ to a more precise estimator whose estimation error is $O(t^2)$, based on fine-grained understanding of the second-order behavior of the variability; once this has been achieved, we can realize the approach outlined above.

We estimate $F$ from the variability while controlling for the addition of motor noise using the estimator $\widehat{\rho^2 t}$:[10]

$$\widehat{F}(\theta) := \frac{\int_0^\theta \frac{1}{\sqrt{var_1(\theta') - \widehat{\rho^2 t}}} d\theta'}{\int_0^{\theta_{\max}} \frac{1}{\sqrt{var_1(\theta') - \widehat{\rho^2 t}}} d\theta'} = \frac{\frac{1}{\sqrt{at}} \int_0^\theta \frac{1}{\sqrt{\mathcal{A}(\theta') + O(t)}} d\theta'}{\frac{1}{\sqrt{at}} \int_0^{\theta_{\max}} \frac{1}{\sqrt{\mathcal{A}(\theta') + O(t)}} d\theta'} = \frac{\int_0^\theta \frac{1}{\sqrt{\mathcal{A}(\theta') + O(t)}} d\theta'}{\int_0^{\theta_{\max}} \frac{1}{\sqrt{\mathcal{A}(\theta') + O(t)}} d\theta'}$$

[10] An essentially equivalent approach would be to deconvolve with the variance-$\widehat{\rho^2}$ motor distribution and apply Lemma S10.

As $A$ is bounded away from zero, we have

$$\widehat{F}(\theta) = \frac{\int_0^\theta \frac{1}{\sqrt{\mathcal{A}(\theta')}} d\theta'}{\int_0^{\theta_{\max}} \frac{1}{\sqrt{\mathcal{A}(\theta')}} d\theta'} + O(t) = \frac{\int_0^\theta F'(\theta') d\theta'}{\int_0^{\theta_{\max}} F'(\theta') d\theta'} + O(t) = F(\theta) + O(t)$$

The same derivation, without the integral in the numerator, shows

$$\widehat{F}'(\theta) = F'(\theta) + O(t)$$

We will also require the second and third derivatives of $F$ up to error $O(t)$; as discussed in Section S2.6.4, this can be achieved by differentiating $\widehat{F}'(\theta)$:

$$\widehat{F}''(\theta) = F''(\theta) + O(t)$$
$$\widehat{F}'''(\theta) = F'''(\theta) + O(t)$$

This establishes

$$\|\widehat{F}(\theta) - F(\theta)\|_{\infty,4} = O(t) \tag{50}$$

Based on this, set

$$\tilde{\mathcal{A}}(\theta) = \frac{1}{\left(\widehat{F}'(\theta)\right)^2} = \mathcal{A}(\theta) + O(t)$$

with constants depending on $\min_\theta F'(\theta)$. Further, set (where $\theta$ can be chosen arbitrarily; note that $\mathcal{A}$ is nowhere zero):

$$\widehat{at} := \frac{var_1(\theta) - \rho^2 t}{\tilde{\mathcal{A}}(\theta)} = \frac{a\mathcal{A}(\theta)t + O(t^2)}{\mathcal{A}(\theta) + O(t)}$$

Similarly, we set:

$$\widehat{bt} := \frac{var_2(\theta) - \rho^2 t}{\tilde{\mathcal{A}}(\theta)} + O(t^2) = bt + O(t^2)$$

where $\theta$ can be chosen arbitrarily. Next, we use $\frac{bias_1(\theta)}{\widehat{at}}$ to estimate $\tilde{\mathcal{C}}(\theta) = \mathcal{C}(\theta) + O(t)$:

$$\tilde{\mathcal{C}}(\theta) := \frac{bias_1(\theta)}{\widehat{at}} = \frac{bias_1(\theta)}{\frac{var_1(\theta)-\rho^2 t}{\tilde{\mathcal{A}}(\theta)}} = \frac{a\mathcal{C}(\theta)t + O(t^2)}{at + O(t^2)} = \mathcal{C}(\theta) + O(t)$$

Lemma S24 shows that $\mathcal{C}(\theta) = \frac{\frac{d\log P_{prior}(\theta)}{d\theta}}{F'^2(\theta)} - \frac{p+2}{2} \cdot \frac{F''(\theta)}{F'^3(\theta)}$. Thus, we can use the formula of $\mathcal{C}$ and the estimation $\tilde{\mathcal{C}}$ to derive an estimate of $\frac{d\widehat{\log P_{prior}}(\theta)}{d\theta} = \frac{d\log P_{prior}(\theta)}{d\theta} + O(t)$. First, (recall the definition of $\tilde{p}$ in Eq. 36):

$$\mathcal{C}(\theta) = \frac{\frac{d\log P_{prior}(\theta)}{d\theta}}{F'^2(\theta)} - \frac{\tilde{p}+2}{2} \cdot \frac{F''(\theta)}{F'(\theta)^3}$$
$$\frac{d\log P_{prior}}{d\theta} = [\mathcal{C} + \frac{\tilde{p}+2}{2} \cdot \frac{F''(\theta)}{F'(\theta)^3}] \cdot F'(\theta)^2 = \mathcal{C} \cdot F'(\theta)^2 + \frac{\tilde{p}+2}{2} \cdot \frac{F''(\theta)}{F'(\theta)}$$

Based on this, we estimate the derivative of the prior as follows (taking $\tilde{p}' = \tilde{p} + \Delta p$). For readability, here and later, we suppress the argument θ in the stimulus-dependent functions:

$$\begin{aligned}\frac{d\widehat{\log P_{prior}}}{d\theta} &= [\tilde{C} + \frac{\tilde{p}'+2}{2}\cdot\frac{\widehat{F}''}{\widehat{F}'^3}]\cdot\widehat{F}'^2 \\ &= (C+O(t))\cdot(F'+O(t)))^2 + \frac{\tilde{p}'+2}{2}\cdot\frac{F''+O(t)}{F'+O(t)} \\ &= C\cdot F'^2 + O(t) + \frac{\tilde{p}'+2}{2}\cdot[\frac{F''}{F'}+O(t)] \\ &= C\cdot F'^2 + \frac{\tilde{p}'+2}{2}\cdot\frac{F''}{F'} + O(t) \\ &= \frac{d\log P_{prior}}{d\theta} + \frac{\Delta P}{2}\cdot\frac{F''}{F'} + O(t)\end{aligned}$$

By differentiating, we establish

$$\left\|\log\widehat{p_{prior}}(\theta) - (\mathrm{U} + \log\left(F'\right)^{\frac{p'-p}{2}} + \log p_{prior}(\theta))\right\|_{\infty,3} = O(t) \tag{51}$$

for the scalar

$$U = \frac{1}{\int (F'(\theta))^{\frac{p'-p}{2}}\cdot p_{prior}(\theta)d\theta} \tag{52}$$

Then, based on the connection between the expressions for $\mathcal{B}$ and $C$, we can derive an estimation for $\mathcal{B}$. By Lemma S24,

$$\mathcal{B} = \frac{2g}{F'^2} + \frac{F''}{2F'^6}$$

where

$$g = \frac{d}{d\theta}C_{dec} = \frac{[(F')^2\cdot\frac{d^2 logP_{prior}}{d\theta^2} - 2F'F''\cdot\frac{d\log p_{prior}}{d\theta}]}{F'^4} - \frac{(\tilde{p}+1)(F'F''' - 3F''^2)}{2F'^4}$$

We can estimate

$$\begin{aligned}\tilde{C_{enc}} :&= \frac{1}{4}\frac{d}{d\theta}\tilde{\mathcal{A}} = \frac{1}{4}\frac{d}{d\theta}\frac{1}{F'^2} + O(t) = C_{enc} + O(t) \\ \tilde{g} :&= \frac{d}{d\theta}\left(\tilde{C} - \tilde{C_{enc}}\right) = \frac{d}{d\theta}C_{dec} + O(t)\end{aligned}$$

This permits us to derive an estimate of $\mathcal{B}$:

$$\tilde{\mathcal{B}} = \frac{2\tilde{g}}{\widehat{F}'^2} + \frac{\widehat{F}''}{2\widehat{F}'^6} = \frac{2g+O(t)}{F'^2+O(t)} + \frac{F''+O(t)}{2F'^6+O(t)} = \frac{2g}{F'^2} + \frac{F''}{2F'^6} + O(t) = \mathcal{B} + O(t)$$

Our aim is now to solve the simultaneous linear equations of $bias_1, bias_2$:

$$\begin{aligned}bias_1 &= aCt + a^2\mathcal{D}t^2 + O(t^3) \\ bias_2 &= bCt + b^2\mathcal{D}t^2 + O(t^3)\end{aligned}$$

in order to obtain an estimate of:

$$\mathcal{D} = \frac{bias_1 - \frac{a}{b}bias_2}{at\cdot(at-bt)} + \frac{O(t^3)}{at\cdot(at-bt)} = \frac{bias_1 - \frac{a}{b}bias_2}{at\cdot(at-bt)} + O(t)$$

To precisely estimate $\mathcal{D}$, we need to obtain a sufficiently precise estimate of $\frac{a}{b}$. We first need to separate $\rho^2 t$, as it otherwise would influence the estimation of $at$. We can differentiate the variance w.r.t. $\theta$, then we have by Lemma S24:

$$\frac{dvar_1}{d\theta} = a\frac{d\mathcal{A}}{d\theta}t + a^2\frac{d\mathcal{B}}{d\theta}t^2 + O(t^3)$$

Rearranging,

$$a\frac{d\mathcal{A}}{d\theta}t = \frac{dvar_1}{d\theta} - a^2\frac{d\mathcal{B}}{d\theta}t^2 + O(t^3)$$

Now by differentiating $\tilde{\mathcal{B}} = \mathcal{B} + O(t)$, and recalling Section S2.6.4,

$$\frac{d\tilde{\mathcal{B}}}{d\theta} = \frac{d\mathcal{B}}{d\theta} + O(t) \qquad\qquad \frac{d\tilde{\mathcal{A}}}{d\theta} = \frac{d\mathcal{A}}{d\theta} + O(t)$$

Further, define

$$\begin{aligned}
\widehat{a^2t^2} &:= \left(\widehat{at}\right)^2 = \left[at + O(t^2)\right]^2 = a^2t^2 + O(t^3) \\
\widehat{b^2t^2} &:= \left(\widehat{bt}\right)^2 = \left[bt + O(t^2)\right]^2 = b^2t^2 + O(t^3) \\
\mathfrak{A} &:= \frac{dvar_1(\theta^*)}{d\theta} - \widehat{a^2t^2}\frac{d\tilde{\mathcal{B}}}{d\theta} = a\frac{d\mathcal{A}(\theta^*)}{d\theta}t + O(t^3) \\
\mathfrak{B} &:= \frac{dvar_2(\theta^*)}{d\theta} - \widehat{b^2t^2}\frac{d\tilde{\mathcal{B}}}{d\theta} = b\frac{d\mathcal{A}(\theta^*)}{d\theta}t + O(t^3)
\end{aligned}$$

This now enables us to derive a precise estimate of $\frac{a}{b}$:

$$\overline{(\frac{a}{b})} = \frac{\mathfrak{A}}{\mathfrak{B}} = \frac{\frac{dvar_1(\theta^*)}{d\theta} - \widehat{a^2t^2}\frac{d\tilde{\mathcal{B}}}{d\theta}}{\frac{dvar_2(\theta^*)}{d\theta} - \widehat{b^2t^2}\frac{d\tilde{\mathcal{B}}}{d\theta}} = \frac{a\frac{d\mathcal{A}(\theta^*)}{d\theta}t + O(t^3)}{b\frac{d\mathcal{A}(\theta^*)}{d\theta}t + O(t^3)} = \frac{a}{b} + O(t^2)$$

where $\theta^*$ can be chosen arbitrarily. The important achievement here is a gap in the order of the error: in this error estimate, there is no term of order $O(t)$. Since we have $\overline{\frac{a}{b}} = \frac{a}{b} + O(t^2)$, we can use these conditions to estimate $\mathcal{D}$:

$$\begin{aligned}
\tilde{\mathcal{D}}(\theta) &:= \frac{bias_1(\theta) - \overline{\frac{a}{b}}bias_2(\theta)}{\widehat{at}\cdot(\widehat{at} - \widehat{bt})} \\
&= \frac{bias_1(\theta) - (\frac{a}{b} + O(t^2))bias_2(\theta)}{(at + O(t^2))\cdot(at - bt + O(t^2))} \\
&= \frac{bias_1(\theta) - \frac{a}{b}bias_2(\theta) + O(t^3)}{at\cdot(at - bt) + O(t^3)} \\
&= [bias_1(\theta) - \frac{a}{b}bias_2(\theta) + O(t^3)]\cdot\frac{1}{at\cdot(at-bt) + O(t^3)} \\
&= \frac{bias_1(\theta) - \frac{a}{b}bias_2(\theta) + O(t^3)}{at\cdot(at - bt)}\cdot\frac{1}{1 + O(t)} \\
&= \frac{bias_1(\theta) - \frac{a}{b}bias_2(\theta) + O(t^3)}{at\cdot(at - bt)}\cdot[1 + O(t)] \\
&= \frac{bias_1(\theta) - \frac{a}{b}bias_2(\theta)}{at\cdot(at - bt)} + O(t) \\
&= \mathcal{D} + O(t)
\end{aligned}$$

Taken together, we have defined estimates $\widehat{F}$, $\widehat{p_{prior}}$, $\tilde{\mathcal{D}}$ satisfying the claimed error estimates. □

##### S2.6.3 Auxiliary Calculations: Second-Order Expansion of Bias and Variance

As reviewed in Section S2.2, in the low-noise regime, the bias is given as (Theorem 1 of [5])

$$bias(\theta) = \frac{\sigma^2}{F'^2}\frac{d}{d\theta}\log p_{prior} + \frac{\tilde{p}+2}{4}\frac{d}{d\theta}\frac{\sigma^2}{F'^2} + O(\sigma^4) \tag{53}$$

The proof of Theorem 3 builds on strengthening this to the following expansion to the next order in the noise magnitude:

**Lemma S23.** *Let $M \in \mathfrak{M}$. Let $\tau$ be the magnitude of motor noise, and $\sigma$ the magnitude of sensory noise. Then, for $\sigma$ close to zero, the following expansion holds:*

$$\begin{aligned} var(\theta) =& \tau^2 + \mathcal{A}_M(\theta)\sigma^2 + \mathcal{B}_M(\theta)\sigma^4 + O(\sigma^6) \\ bias(\theta) =& \mathcal{C}_M(\theta)\sigma^2 + \mathcal{D}_M(\theta)\sigma^4 + O(\sigma^6) \end{aligned}$$

*where $O(\cdot)$ includes constants depending on $M$ but not $\theta$, and where $\mathcal{A}_M, \mathcal{B}_M, \mathcal{C}_M, \mathcal{D}_M$ are differentiable functions.*

We will directly show this expansion by providing explicit expressions for the coefficients $\mathcal{A}, \mathcal{B}, \mathcal{C}, \mathcal{D}$. First, the result cited above is equivalent to stating that

$$\mathcal{C}_M(\theta) = \frac{1}{F'^2}\frac{d}{d\theta}\log p_{prior}(\theta) + \frac{\tilde{p}+2}{4}\frac{d}{d\theta}\frac{1}{F'(\theta)^2} \tag{54}$$

We tackle $\mathcal{A}, \mathcal{B}$ in Lemma S24, and $\mathcal{D}$ in Lemma S25. Throughout, we will require the Decoding Function $f$ introduced in Definition S11.

The decoding function also has an expansion in powers of $\sigma$:

$$f(\theta) = \theta + \mathcal{C}_{dec,M}(\theta)\sigma^2 + \mathcal{D}_{dec,M}(\theta)\sigma^4 + O(\sigma^6) \tag{55}$$

where the expression for $\mathcal{C}_{dec,M}(\theta)$ was likewise obtained in [5]:

$$\mathcal{C}_{dec,M}(\theta) = \mathcal{C}_M(\theta) - \frac{1}{4}\frac{d}{d\theta}\frac{1}{F'^2}$$

and $\mathcal{D}_{dec,M}$ will be obtained explicitly in Lemma S25. We first show:

**Lemma S24.** *Let $M \in \mathfrak{M}$. Then, in the decomposition*

$$var(\theta) = \tau^2 + \mathcal{A}_M(\theta)\sigma^2 + \mathcal{B}_M(\theta)\sigma^4 + O(\sigma^6) \tag{56}$$

*we have*

$$\begin{aligned} \mathcal{A}_M(\theta) &= \frac{1}{F'(\theta)^2} \\ \mathcal{B}_M(\theta) &= \frac{2g(\theta)}{F'^2(\theta)} + \frac{F''^2(\theta)}{2F'^6(\theta)} \end{aligned}$$

*where*

$$g = \frac{d}{d\theta}\mathcal{C}_{dec,M} = \frac{[(F')^2 \cdot \frac{d^2 logP_{prior}}{d\theta^2} - 2F'F'' \cdot \frac{dlogP_{prior}}{d\theta}]}{F'^4} - \frac{(p+1)(F'F''' - 3F''^2)}{2F'^4}$$

*Proof.* Recall the decoding function $f$ (Definition S11). Applying a Taylor Expansion to $\widehat{\theta}$ at $m_0 = F(\theta)$, we find:

$$\begin{aligned} \widehat{\theta} &= f(F^{-1}(m)) \\ &= f(F^{-1}(F(\theta))) + \frac{\partial f}{\partial\theta}\cdot[F^{-1}(m) - F^{-1}(F(\theta))] + \frac{1}{2}\cdot\frac{\partial^2 f}{\partial\theta^2}\cdot[F^{-1}(m) - F^{-1}(F(\theta))]^2 + O(F^{-1}(m)^3) \\ &= f(\theta) + \frac{\partial f}{\partial\theta}\cdot[F^{-1}(m) - \theta] + \frac{1}{2}\cdot\frac{\partial^2 f}{\partial\theta^2}\cdot[F^{-1}(m) - \theta]^2 + O\left(F^{-1}(m)^3\right) \end{aligned}$$

The variance of the estimate $\widehat{\theta}$, conditioned on the true stimulus $\theta$, can be expressed as:

$$\begin{aligned}
var(\widehat{\theta}) =& var\left(f(F^{-1}(m))\right) \\
=& var\left(f(\theta)+\frac{\partial f}{\partial\theta}\cdot F^{-1}(m)+\frac{1}{2}\cdot\frac{\partial^2 f}{\partial\theta^2}\cdot[F^{-1}(m)]^2+O\left(F^{-1}(m)^3\right)\right) \\
=& var\left(\frac{\partial f}{\partial\theta}\cdot F^{-1}(m)+\frac{1}{2}\cdot\frac{\partial^2 f}{\partial\theta^2}\cdot\left[F^{-1}(m)\right]^2\right)+O(\sigma^6) \\
=& \left(\frac{\partial f}{\partial\theta}\right)^2\cdot var\left(F^{-1}(m)\right)+\left(\frac{1}{2}\cdot\frac{\partial^2 f}{\partial\theta^2}\right)^2\cdot var\left([F^{-1}(m)]^2\right) \\
&+2\left(\frac{\partial f}{\partial\theta}\right)\cdot\left(\frac{1}{2}\cdot\frac{\partial^2 f}{\partial\theta^2}\right)\cdot cov\left(F^{-1}(m),[F^{-1}(m)]^2\right)+O\left(\sigma^6\right)
\end{aligned}$$

where we dropped $f(\theta)$ as it is a constant.

Thus, we need to calculate $var\left(F^{-1}(m)\right)$, $\frac{\partial f}{\partial\theta}$, $\frac{\partial^2 f}{\partial\theta^2}$, $var\left([F^{-1}(m)]^2\right)$, $cov\left(F^{-1}(m),[F^{-1}(m)]^2\right)$.

In computing variances and covariances of encoding-related quantities, we will assume, without loss of generality, $\theta=0$ by shifting the stimulus space.

- We first study the variance of the encoding, $var(F^{-1}(m))$, where $m=F(\theta)+\delta$:

$$\begin{aligned}
F^{-1}(m) &= F^{-1}(F(\theta)+\delta) \\
&= F^{-1}(F(\theta))+\frac{d}{d\theta}F^{-1}(F((\theta))\cdot\delta+\frac{1}{2}\cdot\frac{d^2}{d\theta^2}F^{-1}(F(\theta))\cdot\delta^2+O(\delta^3) \\
&= \theta+\frac{1}{F'(\theta)}\cdot\delta+\frac{1}{2}\cdot\frac{d(\frac{1}{F'(\theta)})}{dF(\theta)}\cdot\delta^2+O(\delta^3)
\end{aligned}$$

Taking the variance on both sides,

$$\begin{aligned}
var(F^{-1}(m)) &= var(\theta+\frac{1}{F'(\theta)}\cdot\delta+O(\delta^2)) \\
&= var\left(\theta+\frac{1}{F'(\theta)}\cdot\delta+\frac{1}{2}\cdot\frac{d(\frac{1}{F'(\theta)})}{dF(\theta)}\cdot\delta^2+O(\delta^3)\right)
\end{aligned}$$

With the substitution $y=F(\theta)$, we have

$$\frac{d(\frac{1}{F'(\theta)})}{dF(\theta)}=\frac{d(\frac{d\theta}{dy})}{dy}=\frac{d(\frac{1}{\frac{dy}{d\theta}})}{dy}=\frac{d(\frac{1}{\frac{dy}{d\theta}})}{d\theta}\cdot\frac{d\theta}{dy}=\frac{d(y')^{-1}}{d\theta}\cdot\frac{1}{y'}=\frac{\frac{dy'}{d\theta}}{-y'^2}\cdot\frac{1}{y'}=-\frac{y''}{y'^3}.$$

Inserting into the above,

$$\begin{aligned}
var(F^{-1}(m)) &= var\left(\theta+\frac{1}{F'(\theta)}\cdot\delta+\frac{1}{2}\cdot\frac{d(\frac{1}{F'(\theta)})}{dF(\theta)}\cdot\delta^2+O(\delta^3)\right) \\
&= var(\theta)+var\left(\frac{1}{F'}\cdot\delta\right)+var\left(\frac{1}{2}\cdot\left(-\frac{F''}{F'^3}\right)\cdot\delta^2\right)+var\left(O(\delta^3)\right) \\
&= 0+\frac{\sigma^2}{F'^2}+var\left(\frac{1}{2}\cdot\left(-\frac{F''}{F'^3}\right)\cdot\delta^2\right)+O(\sigma^6) \\
&= \frac{\sigma^2}{F'^2}+\frac{1}{4}\cdot\frac{F''^2}{F'^6}\cdot var(\delta^2)+O(\sigma^6)
\end{aligned}$$

since $var(\delta)=\mathbb{E}(\delta^2)-\mathbb{E}(\delta)^2=\mathbb{E}(\delta^2)=\sigma^2, var\left(\delta^2\right)=\mathbb{E}(\delta^4)-[\mathbb{E}(\delta^2)]^2=\mathbb{E}(\delta^4)-\sigma^4$.

Additionally, by the definition of $\delta$, we have $\mathbb{E}(\delta^4) = 3\sigma^4$. Thus, $var(\delta^2) = 3\sigma^4 - \sigma^4 = 2\sigma^4$.

In summary,

$$\begin{aligned} var(F^{-1}(m)) &= \frac{\sigma^2}{F'^2} + \frac{1}{4} \cdot \frac{F''^2}{F'^6} \cdot var(\delta^2) + O(\sigma^6) \\ &= \frac{\sigma^2}{F'^2} + \frac{1}{4} \cdot \frac{F''^2}{F'^6} \cdot 2\sigma^4 + O(\sigma^6) \\ &= \frac{\sigma^2}{F'^2} + \frac{F''^2}{2F'^6} \cdot \sigma^4 + O(\sigma^6) \end{aligned}$$

- Next, we tackle $\frac{\partial f}{\partial \theta}$, the first derivative of the decoding function $f$. We have

$$\frac{df}{d\theta} = 1 + g(\theta)\sigma^2 + O(\sigma^4) \tag{57}$$

where

$$g(\theta) = \frac{d}{d\theta} C_{dec}(\theta) = \frac{[(F')^2 \cdot \frac{d^2 \log P_{prior}}{d\theta^2} - 2F'F'' \cdot \frac{d \log P_{prior}}{d\theta}]}{F'^4} - \frac{(p+1)(F'F''' - 3F''^2)}{2F'^4} \tag{58}$$

- The second derivative $\frac{\partial^2 f}{\partial \theta^2}$ is obtained by differentiating our previous result once more.

$$\frac{d^2 f}{d\theta^2} = \frac{d(\frac{df}{d\theta})}{d\theta} = \sigma^2 \frac{d^2}{d\theta^2} C_{dec} + O(\sigma^4)$$

Evaluating an explicit formula is straightforward, but it will be sufficient to note that this term has order $O(\sigma^2)$.

- The variance of the squared encoding, $var([F^{-1}(m)]^2)$, is computed as follows.

We expand $F^{-1}(m) = \theta + \frac{1}{F'} \cdot \delta + \frac{1}{2} \cdot (-\frac{F''}{F'^3}) \cdot \delta^2 + O(\delta^3)$.

Thus,

$$\begin{aligned} F^{-1}(m)^2 &= \left(\theta + \frac{1}{F'} \cdot \delta + \frac{1}{2} \cdot (-\frac{F''}{F'^3}) \cdot \delta^2 + O(\delta^3)\right)^2 \\ &= \theta^2 + \left(\frac{\delta}{F'}\right)^2 + \left(-\frac{F''}{2F'^3} \cdot \delta^2\right)^2 + 2\theta \cdot \frac{\delta}{F'} + \theta \cdot \left(-\frac{F''}{F'^3} \cdot \delta^2\right) + \frac{\delta}{F'} \cdot \left(-\frac{F''}{F'^3}\right) \cdot \delta^2 + O(\delta^3) \\ &= \theta^2 + \left(\frac{\delta}{F'}\right)^2 + 2\theta \cdot \frac{\delta}{F'} + \theta \cdot \left(-\frac{F''}{F'^3} \cdot \delta^2\right) + O(\delta^3) \\ &= \left(\frac{\delta}{F'}\right)^2 + O(\delta^3) \end{aligned}$$

at $\theta = 0$. Taking the variance, we find:

$$var\left(F^{-1}(m)^2\right) = var\left(\theta^2 + \left(\frac{\delta}{F'}\right)^2 + 2\theta \cdot \frac{\delta}{F'} + \theta \cdot \left(-\frac{F''}{F'^3} \cdot \delta^2\right) + O(\delta^3)\right) = var\left(\left(\frac{\delta}{F'}\right)^2 + O(\delta^3)\right). \tag{59}$$

Since $\mathbb{E}(\delta) = 0$, $cov(\delta, \delta^2) = 0$, we obtain

$$var(F^{-1}(m)^2) = \frac{2\sigma^4}{F'^4} + O(\sigma^6) \tag{60}$$

- We now investigate $cov\left(F^{-1}(m),[F^{-1}(m)]^2\right)$.
  By definition, $cov(F^{-1}(m),[F^{-1}(m)]^2) = \mathbb{E}(F^{-1}(m)\cdot[F^{-1}(m)]^2) - \mathbb{E}(F^{-1}(m))\mathbb{E}([F^{-1}(m)]^2)$
  From the above we know $F^{-1}(m) = \theta + \frac{1}{F'}\cdot\delta + \frac{1}{2}\cdot\left(-\frac{F''}{F'^3}\right)\cdot\delta^2 + O(\delta^3)$. Let $\alpha(\theta) = \frac{1}{F'}, \beta(\theta) = -\frac{F''}{2F'^3}$, we have $F^{-1}(m) = \theta + \alpha\cdot\delta + \beta\cdot\delta^2 + O(\delta^3)$.
  Thus, again at $\theta = 0$,

$$\begin{aligned}
\mathbb{E}(F^{-1}(m)) &= \theta + \alpha\mathbb{E}(\delta) + \beta\mathbb{E}(\delta^2) + \mathbb{E}(O(\delta^3)) \\
&= \beta\sigma^2 + O(\sigma^4) \\
\mathbb{E}([F^{-1}(m)]^2) &= \theta^2 + 2\theta\alpha\mathbb{E}(\delta) + \alpha^2\mathbb{E}(\delta^2) + 2\theta\beta\mathbb{E}(\delta^2) + \mathbb{E}(O(\delta^3)) \\
&= \alpha^2\sigma^2 + O(\sigma^4) \\
\mathbb{E}([F^{-1}(m)]^3) &= \alpha^3\mathbb{E}(\delta^3) + 3\alpha^2\beta\mathbb{E}(\delta^4) + \mathbb{E}(O(\delta^5)) \\
&= 9\alpha^2\beta\sigma^4 + O(\sigma^6) \\
cov(F^{-1}(m),[F^{-1}(m)]^2) &= \mathbb{E}([F^{-1}(m)^3]) - \mathbb{E}(F^{-1}(m))\mathbb{E}([F^{-1}(m)]^2) \\
&= 9\alpha^2\beta\sigma^4 - \alpha^2\beta\sigma^4 + O(\sigma^6) \\
&= \frac{-4F''}{(F')^5}\sigma^4 + O(\sigma^6)
\end{aligned}$$

  Importantly, this is $O(\sigma^4)$.

Putting all of these results together, we find:

$$\begin{aligned}
var(\widehat{\theta}) = &\left(\underbrace{\frac{\partial f}{\partial\theta}}_{1+\sigma^2 C'+O(\sigma^4)}\right)^2 \cdot \underbrace{var(F^{-1}(m))}_{\frac{\sigma^2}{F'^2}+\frac{F''^2}{2F'^6}\cdot\sigma^4+O(\sigma^6)} + \underbrace{\left(\frac{1}{2}\cdot\frac{\partial^2 f}{\partial\theta^2}\right)^2 \cdot var\left([F^{-1}(m)]^2\right)}_{O(\sigma^6)} \\
&\underbrace{+2\left(\frac{\partial f}{\partial\theta}\right)\cdot\left(\frac{1}{2}\cdot\frac{\partial^2 f}{\partial\theta^2}\right)\cdot cov\left(F^{-1}(m),[F^{-1}(m)]^2\right)}_{O(\sigma^6)} + O(\sigma^6) \\
&= \left(1+\sigma^2 C'\right)^2\cdot\left(\frac{\sigma^2}{F'^2} + \frac{F''^2}{2F'^6}\cdot\sigma^4\right) + O(\sigma^6) \\
&= \sigma^2\underbrace{\frac{1}{F'^2}}_{\mathcal{A}(\theta)} + \underbrace{\left(\frac{2g(\theta)}{F'^2} + \frac{F''^2}{2F'^6}\right)}_{\mathcal{B}(\theta)}\cdot\sigma^4 + O(\sigma^6)
\end{aligned}$$

where $g$ is given by (58).

□

**Lemma S25.** *Assume that $M = \langle F', p_{prior}, p\rangle \in \mathfrak{M}$. Let $f$ be the function mapping $F^{-1}(m)$ to the estimate $\widehat{\theta}$. Then, for the expansion*

$$\begin{aligned}
f(\theta) &= \mathcal{C}_{dec,M}(\theta)\sigma^2 + \mathcal{D}_{dec,M}(\theta)\sigma^4 + O(\sigma^6) \\
bias(\theta) &= \mathcal{C}_M(\theta)t + \mathcal{D}_M(\theta)t^2 + O(t^3),
\end{aligned}$$

*we have*

$$\mathcal{D}_M(\theta) = \mathcal{D}_{dec,M}(\theta) + \mathcal{C}_{enc,M}(\theta)\mathcal{C}'_{dec,M}(\theta) + \mathcal{D}_{enc,M}(\theta) + \frac{1}{2}\mathcal{A}_M(\theta)\mathcal{C}''_{dec,M}(\theta) \tag{61}$$

*and the coefficient $\mathcal{D}_{dec,M}$ has the following properties:*

1. *Across $p = 2, 4, 6, \ldots$, it is polynomial in $p$ with degree at most 4. The coefficients of this polynomial are combined products of these factors and their powers*

$$(\log p_{prior})'(\theta), \quad (\log p_{prior})''(\theta), \quad (\log p_{prior})'''(\theta), \quad F'(\theta), \quad F''(\theta), \quad F'''(\theta), \quad F''''(\theta), \quad \frac{1}{F'(\theta)}, \quad \frac{1}{p_{prior}(\theta)}$$

.

2. *At $p = 0$, the expression $\mathcal{D}_{\langle F', p_{prior}, p\rangle}(\theta)$ is a polynomial in*

$$(\log p_{prior})'(\theta), \quad (\log p_{prior})''(\theta), \quad F'(\theta), \quad F''(\theta), \quad F'''(\theta), \quad F''''(\theta), \quad \frac{1}{F'(\theta)}$$

3. *At $p = 1$, the expression $\mathcal{D}_{\langle F', p_{prior}, p\rangle}(\theta)$ is a polynomial in*

$$(\log p_{prior})'(\theta), \quad F'(\theta), \quad F''(\theta), \quad F'''(\theta), \quad F''''(\theta), \quad \frac{1}{F'(\theta)}$$

*Proof.* The expression for $\mathcal{C}_{dec,M}$ was already shown by Hahn and Wei [5].

First, recall that the bias consists of encoding bias and decoding bias [5], i.e.

$$\mathbb{E}[\hat{\theta}] - \theta = \underbrace{E[F^{-1}(m)] - \theta}_{Encoding\ Bias} + \underbrace{E[\hat{\theta}] - E[F^{-1}(m)]}_{Decoding\ Bias}.$$

First, we treat the encoding bias, which is independent of the loss function. We use a Taylor expansion to expand $F^{-1}(m)$ at $m = F(\theta)$,

$$F^{-1}(m) = \theta + \frac{\delta}{F'} - \frac{F''}{2F'^3} \cdot \delta^2 + \frac{F^{-1^{(3)}}}{3!} \cdot \delta^3 + \frac{F^{-1^{(4)}}}{4!} \cdot \delta^4 + O(\delta^5)$$

Taking expectations over $\delta$,

$$\begin{aligned}
\mathbb{E}[F^{-1}(m)] &= \theta + \frac{E[\delta]}{F'} - \frac{F''}{2F'^3} E[\delta^2] + \frac{F^{-1^{(3)}}}{3!} E[\delta^3] + \frac{F^{-1^{(4)}}}{4!} E[\delta^4] + \frac{F^{-1^{(5)}}}{5!} E[\delta^5] + \mathbb{E}[O(\delta^6)] \\
&= \theta - \frac{F''}{2F'^3} t + \frac{F^{-1^{(4)}}}{4!} 3t^2 + O(\delta^6)
\end{aligned}$$

where we have used

$$\mathbb{E}[\delta] = 0, E[\delta^2] = t, E[\delta^3] = 0, E[\delta^4] = 3t^2, E[\delta^5] = 0, E[\delta^6] = 15t^3$$

Further, we find

$$\begin{aligned}
\frac{d^3 F^{-1}}{dm^3} &= -\frac{F'''}{F'^4} + 3\frac{F''^2}{F'^5} \\
\frac{d^4 F^{-1}}{dm^4} &= \frac{-F^{(4)} F'^2 + 10F'F''F''' - 15F''^3}{F'^7}
\end{aligned}$$

Inserting into the above, we find:

$$\mathbb{E}[F^{-1}(m)] - \theta = -\frac{F''}{2F'^3} \cdot t + \frac{-F^{(4)}F'^2 + 10F'F''F''' - 15F''^3}{8F'^7} t^2 + O(t^3) \tag{62}$$

Now let $f$ be the decoding function mapping $F^{-1}(m)$ to the estimate $\widehat{\theta}$. Then the bias is

$$\begin{aligned}
\mathbb{E}[\widehat{\theta}] - \theta =& \mathbb{E}_{m|\theta}[f(F^{-1}(m))] - \theta \\
=& \mathbb{E}_{m|\theta}[f(\theta) + (F^{-1}(m) - \theta)f'(\theta) + \frac{1}{2}(F^{-1}(m) - \theta)^2 f''(\theta) + h.o.t.] - \theta
\end{aligned}$$

Using the expansion (and suppressing the $M$ subscript where unambiguous),

$$\begin{aligned}
f(\theta) =& \theta + t\mathcal{C}_{dec}(\theta) + t^2\mathcal{D}_{dec}(\theta) + O(t^3) \\
\mathbb{E}_{m|\theta}[F^{-1}(m)] =& \theta + t\mathcal{C}_{enc}(\theta) + t^2\mathcal{D}_{enc}(\theta) + O(t^3)
\end{aligned}$$

we find that the bias equals – taking, without loss of generality, $\theta = 0$ for simplicity:

$$\begin{aligned}
=& t\mathcal{C}_{dec}(\theta) + t^2\mathcal{D}_{dec}(\theta) + \mathbb{E}_{m|\theta}[(F^{-1}(m))](1 + t\mathcal{C}'_{dec}(\theta)) + \frac{1}{2}\mathbb{E}_{m|\theta}[(F^{-1}(m))^2](t\mathcal{C}''_{dec}(\theta)) + O(t^3)] \\
=& t\mathcal{C}_{dec}(\theta) + t^2\mathcal{D}_{dec}(\theta) + (t\mathcal{C}_{enc}(\theta) + t^2\mathcal{D}_{enc}(\theta))(1 + t\mathcal{C}'_{dec}(\theta)) + \frac{1}{2}t\mathcal{A}(\theta)(t\mathcal{C}''_{dec}(\theta)) + O(t^3)] \\
=& t[\mathcal{C}_{enc}(\theta) + \mathcal{C}_{dec}(\theta)] \\
&+ t^2\left[\mathcal{D}_{dec}(\theta) + \mathcal{C}_{enc}(\theta)\mathcal{C}'_{dec}(\theta) + \mathcal{D}_{enc}(\theta) + \frac{1}{2}\mathcal{A}(\theta)\mathcal{C}''_{dec}(\theta)\right] + O(t^3)]
\end{aligned}$$

In particular,

$$\mathcal{D}(\theta) = \mathcal{D}_{dec}(\theta) + \mathcal{C}_{enc}(\theta)\mathcal{C}'_{dec}(\theta) + \mathcal{D}_{enc}(\theta) + \frac{1}{2}\mathcal{A}(\theta)\mathcal{C}''_{dec}(\theta) \tag{63}$$

It remains to calculate $\mathcal{D}_{dec}(\theta)$. We need to distinguish the decoding bias by the loss function exponent $p$.

- $p = 2, 4, 6, ...$

  We follow the same approach as in Hahn and Wei [5], but carry out the calculations to the next order in $t$. As done there, we first define

$$q_{prior}(m) := \frac{P_{prior}(m)}{\sqrt{\mathcal{S}(F^{-1}(m))}}$$

  Let $r := \widehat{\theta} - F^{-1}(m)$. To solve for $\widehat{\theta}$, we expand – taking without loss of generality $\theta, F(\theta) = 0$ here:

$$\begin{aligned}
r =& Rt + St^2 + O(t^3) \\
F^{-1}(x) =& m + Mx + Nx^2 + Px^3 + Qx^4 + O(x^5) \\
q_{prior}(x) =& q_{prior}(m) + Tx + Ux^2 + Vx^3 + O(x^4)
\end{aligned}$$

  and then set

$$\begin{aligned}
0 =& \int (r - F^{-1}(x))^{2q-1} q_{prior}(x) \frac{1}{\sqrt{2\pi t}} \exp\left(-\frac{x^2}{2t}\right) dx \\
=& \int (Rt + St^2 + Mx + Nx^2 + Px^3 + Qx^4)^{2q-1} \left(q_{prior}(m) + Tx + Ux^2 + Vx^3\right) \frac{1}{\sqrt{2\pi t}} \exp\left(-\frac{x^2}{2t}\right) dx \\
=& \sum \binom{2q-1}{n_1 n_2 n_3 n_4 n_5 n_6} R^{n_1} S^{n_2} M^{n_3} N^{n_4} P^{n_5} Q^{n_6} t^{n_1+2n_2} \int x^{n_3+2n_4+3n_5+4n_6} \frac{1}{\sqrt{t}} \left(q_{prior}(m) + Tx + Ux^2 + Vx^3\right) \exp\left(-\frac{x^2}{2t}\right) dx
\end{aligned}$$

where the sum runs over all $n_1+n_2+n_3+n_4+n_5+n_6=2q-1$. We know that each increase in $n_1$ will contribute to one increase of $t$'s order. Similarly, $n_2\sim 2, n_3\sim \frac{1}{2}, n_4\sim 1, n_5\sim \frac{3}{2}, n_6\sim 2$. Therefore, we can find all the $t^{q+1}$ terms in the equation. The sum of all these terms should be 0, i.e.

$$0=$$
$$[\frac{(2q-1)!}{(2q-2)!1!}SM^{2q-2}q_{prior}(m)(2q-3)!!+\frac{(2q-1)!}{(2q-2)!1!}QM^{2q-2}q_{prior}(m)(2q-3)!!+\frac{(2q-1)!}{(2q-4)!2!}RM^{2q-4}N^2q_{prior}(m)(2q-1)!!$$
$$+\frac{(2q-1)!}{(2q-4)!3!}R^3M^{2q-4}q_{prior}(m)(2q-5)!!+\frac{(2q-1)!}{(2q-4)!2!}R^2M^{2q-4}Nq_{prior}(m)(2q-3)!!+\frac{(2q-1)!}{(2q-4)!3!}M^{2q-4}N^3q_{prior}(m)(2q+1)$$
$$+\frac{(2q-1)!}{(2q-3)!2!}R^2M^{2q-3}T(2q-3)!!+\frac{(2q-1)!}{(2q-3)!2!}M^{2q-3}N^2T(2q+1)!!+\frac{(2q-1)!}{(2q-3)!}RM^{2q-3}NT(2q-1)!!$$
$$+\frac{(2q-1)!}{(2q-2)!}M^{2q-2}PT(2q+1)!!+\frac{(2q-1)!}{(2q-2)!}RM^{2q-2}U(2q-1)!!+\frac{(2q-1)!}{(2q-2)!}M^{2q-2}NU(2q+1)!!+$$
$$\frac{(2q-1)!}{(2q-1)!}M^{2q-1}V(2q+1)!!]\cdot t^{q+1}+O(t^{q+2})$$

We divide these terms by $(2q-1)M^{2q-2}q_{prior}(m)t^{q+1}(2q-3)!!$ to get

$$0=S+Q+\frac{(2q-3)(2q-2)(2q-1)}{2}RM^{-2}N^2+\frac{(2q-2)}{2}R^2M^{-1}Tq^{-1}_{prior}(m)$$
$$+\frac{(2q-2)(2q-1)(2q+1)}{2}A^{-1}N^2Tq^{-1}_{prior}(m)+(2q-1)RUq^{-1}_{prior}(m)+(2q+1)MVq^{-1}_{prior}(m)$$
$$+\frac{2q-2}{6}R^3M^{-2}+\frac{(2q-3)(2q-2)}{2}R^2M^{-2}N+\frac{(2q-3)(2q-2)(2q-1)(2q+1)}{6}M^{-2}N^3$$
$$+(2q-2)(2q-1)RM^{-1}NTq^{-1}_{prior}(m)+(2q-1)(2q+1)PTq^{-1}_{prior}(m)+(2q-1)(2q+1)NUq^{-1}_{prior}(m)+O(t)$$

Hence, the desired second-order component of the decoding bias is:

$$S=-[Q+\frac{(2q-3)(2q-2)(2q-1)}{2}RM^{-2}N^2+\frac{(2q-2)}{2}R^2M^{-1}Tq^{-1}_{prior}(m)$$
$$+\frac{(2q-2)(2q-1)(2q+1)}{2}M^{-1}N^2Tq^{-1}_{prior}(m)+(2q-1)RUq^{-1}_{prior}(m)+(2q+1)MVq^{-1}_{prior}(m)$$
$$+\frac{2q-2}{6}R^3M^{-2}+\frac{(2q-3)(2q-2)}{2}R^2M^{-2}N+\frac{(2q-3)(2q-2)(2q-1)(2q+1)}{6}M^{-2}N^3$$
$$+(2q-2)(2q-1)RM^{-1}NTq^{-1}_{prior}(m)+(2q-1)(2q+1)PTq^{-1}_{prior}(m)+(2q-1)(2q+1)NUq^{-1}_{prior}(m)]+O(t) \quad (64)$$

This $S$ is equal to the desired $\mathcal{D}_{dec}$.

- $p=0$

  As in [22, 5], we obtain the MAP estimator by considering a stationary point of the log-posterior, but expand to the second order:

$$\frac{d}{d\theta}\log P(\widehat{\theta}|m)=0$$
$$\begin{aligned}0&=t\frac{d}{d\theta}\log P(\widehat{\theta}|m)\\&=t\frac{d}{d\theta}\log P(m|\widehat{\theta})+t\frac{d}{d\theta}\log P_{prior}(\widehat{\theta})\\&=-(\frac{d}{d\theta}F(\widehat{\theta}))\cdot(F(\widehat{\theta})-m)+t\frac{d}{d\theta}\log P_{prior}(\widehat{\theta})\\&=-\mathcal{S}\cdot\widehat{\theta}-\frac{1}{2}F'F''\cdot\widehat{\theta}^2+t\frac{d}{d\theta}\log P_{prior}(\widehat{\theta})+O(\widehat{\theta}^3)\end{aligned}$$

$$= -\mathcal{S}\cdot\widehat{\theta} - \frac{1}{2}F'F''\cdot\widehat{\theta}^2 + t\frac{d}{d\theta}\log P_{prior}(0) + t\widehat{\theta}\frac{d^2}{d\theta^2}\log P_{prior}(0) + O(\widehat{\theta}^3)$$

$$\mathcal{S}\cdot\widehat{\theta} = -\frac{1}{2}F'F''\cdot\widehat{\theta}^2 + t\frac{d}{d\theta}\log P_{prior}(0) + t\widehat{\theta}\frac{d^2}{d\theta^2}\log P_{prior}(0) + O(\widehat{\theta}^3)$$

We thus obtain

$$\widehat{\theta} = -\frac{F''}{2F'}\cdot\widehat{\theta}^2 + t\frac{1}{\mathcal{S}}\frac{d}{d\theta}\log P_{prior}(0) + t\frac{1}{\mathcal{S}}\widehat{\theta}\frac{d^2}{d\theta^2}\log P_{prior}(0) + O(\widehat{\theta}^3)$$

Since $\widehat{\theta}$ is of order $O(t)$, we let $\widehat{\theta} = Pt + Qt^2 + O(t^3)$ and plug back in the equation. By comparing the coefficients of the same order of $t$, we can get the refined bias:

$$\begin{aligned}
\widehat{\theta} &= Pt + Qt^2 + O(t^3)\\
&= -\frac{F''}{2F'}\cdot(Pt + Qt^2 + O(t^3))^2 + t\frac{1}{\mathcal{S}}\frac{d}{d\theta}\log P_{prior}(0) + t\frac{1}{\mathcal{S}}(Pt + Qt^2 + O(t^3))\frac{d^2}{d\theta^2}\log P_{prior}(0) + O(t^3)\\
&= -\frac{F''}{2F'}\cdot P^2t^2 + t\frac{1}{\mathcal{S}}\frac{d}{d\theta}\log P_{prior}(0) + Pt^2\frac{1}{\mathcal{S}}\frac{d^2}{d\theta^2}\log P_{prior}(0) + O(t^3)
\end{aligned}$$

We obtain the terms of the expansion:

$$\begin{aligned}
P &= \frac{1}{\mathcal{S}}\frac{d}{d\theta}\log P_{prior}(0)\\
Q &= -\frac{F''}{2F'}\cdot P^2 + P\frac{1}{\mathcal{S}}\frac{d^2}{d\theta^2}\log P_{prior}(0)\\
&= -\frac{F''}{2F'}\cdot\frac{1}{\mathcal{S}^2}\cdot\left[\frac{d}{d\theta}\log P_{prior}(0)\right]^2 + \frac{1}{\mathcal{S}^2}\frac{d}{d\theta}\log P_{prior}(0)\frac{d^2}{d\theta^2}\log P_{prior}(0)
\end{aligned}$$

The second-order term $Q$ here is $\mathcal{D}_{dec}$.

- $p = 1$

  Recall that

$$r = \widehat{\theta} - F^{-1}(m)$$
$$F^{-1}(x) = m + Mx + Nx^2 + O(x^3)$$

According to the previous proof in [5], we have:

$$\begin{aligned}
0 &= 2q_{prior}(m)\left(\frac{r}{M\sqrt{t}} - \left(\frac{r}{M\sqrt{t}}\right)^3\cdot\frac{1}{12} + \left(\frac{r}{M\sqrt{t}}\right)^5\cdot\frac{1}{5\cdot16\cdot2!} + \ldots\right)\\
&+ 2\frac{d}{dm}q_{prior}(m)\sqrt{t}\\
&- 2\frac{d}{dm}q_{prior}(m)\sqrt{t}\left(\frac{r}{M\sqrt{t}})^2\cdot\frac{1}{2} - \left(\frac{r}{M\sqrt{t}}\right)^4\cdot\frac{1}{16} + \left(\frac{r}{M\sqrt{t}}\right)^6\cdot\frac{1}{6\cdot16\cdot2!} + \ldots\right)\\
&= \frac{2q_{prior(m)}\cdot r}{M} - \frac{q_{prior}(m)\cdot r^3}{6M^3t} + 2\frac{d}{dm}q_{prior}(m)t - \frac{d}{dm}q_{prior}(m)\cdot\frac{r^2}{M^2} + O(t^3)
\end{aligned}$$

Let $r = Rt + St^2 + O(t^3)$ and compare the coefficients of the same order of t, we can get the value of $S$:

$$
\begin{aligned}
0 = & \frac{2q_{prior}(m) \cdot (Rt + St^2 + O(t^3))}{M} - \frac{q_{prior}(m)(R^3t^3 + O(t^4))}{6M^3t} + 2\frac{d}{dm}q_{prior}(m)t \\
& - \frac{d}{dm}q_{prior}(m) \cdot \frac{R^2t^2 + O(t^3)}{M^2} + O(t^3) \\
= & \frac{2q_{prior}(m) \cdot (Rt + St^2)}{M} - \frac{q_{prior}(m)R^3t^2}{6M^3} + 2\frac{d}{dm}q_{prior}(m)t - \frac{d}{dm}q_{prior}(m) \cdot \frac{R^2t^2}{M^2} + O(t^3)
\end{aligned}
$$

Then we collect all the second order $t$ terms, the sum of these terms should be 0, i.e.

$$
0 = \frac{2q_{prior}(m)S}{M} - \frac{q_{prior}(m)R^3}{6M^3} - \frac{d}{dm}q_{prior}(m) \cdot \frac{R^2}{M^2}
$$

The lowest-order term is

$$
R = \frac{1}{\mathcal{S}}\frac{d}{d\theta}\log\frac{P_{prior}(\theta)}{\sqrt{\mathcal{S}(\theta)}}
$$

Now by solving this equation and using $R$ to express $S$, we have

$$
\begin{aligned}
S = & \frac{R^3}{12M^2} + \frac{R^2}{2M} \cdot \frac{d}{dm}\log q_{prior}(m) \\
= & \frac{R^3}{12M^2} + \frac{R^2}{2M} \cdot \frac{1}{\sqrt{\mathcal{S}}} \cdot \frac{d}{d\theta}\log\frac{P_{prior}(\theta)}{\sqrt{\mathcal{S}}}
\end{aligned}
$$

Inserting the expression for $R$, we obtain:

$$
\begin{aligned}
S = & \left(\frac{1}{\mathcal{S}}\frac{d}{d\theta}\log\frac{P_{prior}(\theta)}{\sqrt{\mathcal{S}(\theta)}}\right)^3 \frac{1}{12M^2} + \left(\frac{1}{\mathcal{S}}\frac{d}{d\theta}\log\frac{P_{prior}(\theta)}{\sqrt{\mathcal{S}(\theta)}}\right)^2 \frac{1}{2M} \cdot \frac{1}{\sqrt{\mathcal{S}}} \cdot \frac{d}{d\theta}\log\frac{P_{prior}(\theta)}{\sqrt{\mathcal{S}}} \\
= & \left(\frac{d}{d\theta}\log\frac{P_{prior}(\theta)}{\sqrt{\mathcal{S}(\theta)}}\right)^3 \cdot \left[\frac{1}{\mathcal{S}}^3 \cdot \frac{1}{12M^2} + \left(\frac{1}{\mathcal{S}}\right)^2 \frac{1}{2M} \cdot \frac{1}{\sqrt{\mathcal{S}}}\right]
\end{aligned}
$$

which provides the desired $\mathcal{D}_{dec}$.

□

##### S2.6.4 Derivative of Remainder

Derivations in Hahn and Wei [5] and in Section S2.6.3 guarantee approximations of bias and variability up to error $O(\sigma^6)$. In the proof of Theorem 3, a somewhat stronger approximation is needed, in the sense that that proof relies on using the derivatives of bias and variability to estimate various properties of the model. Formally, to show that this is valid, we need to show the following. Consider bias and variability, as functions of the stimulus $\theta$ and the noise magnitude $t = \sigma^2$:

$$
\begin{aligned}
var(\theta) &= t\mathcal{A}(\theta) + t^2\mathcal{B}(\theta) + O(t^3) \\
bias(\theta) &= t\mathcal{C}(\theta) + t^2\mathcal{D}(\theta) + O(t^3)
\end{aligned}
\tag{65}
$$

we want to show that this approximation is well-behaved under taking derivatives:

$$
\begin{aligned}
\frac{d}{d\theta}var(\theta) &= t\frac{d}{d\theta}\mathcal{A}(\theta) + t^2\frac{d}{d\theta}\mathcal{B}(\theta) + O(t^3) \\
\frac{d}{d\theta}bias(\theta) &= t\frac{d}{d\theta}\mathcal{C}(\theta) + t^2\frac{d}{d\theta}\mathcal{D}(\theta) + O(t^3)
\end{aligned}
\tag{66}
$$

and similarly for higher derivatives as far as is needed. This might a priori be unobvious because the $O(t^3)$ term is dependent on θ, so its derivatives need not a priori be $O(t^3)$. More explicitly, we write

$$\begin{aligned} var(\theta) &= t\mathcal{A}(\theta) + t^2\mathcal{B}(\theta) + t^3 S(t,\theta) \\ bias(\theta) &= t\mathcal{C}(\theta) + t^2\mathcal{D}(\theta) + t^3 R(t,\theta) \end{aligned} \tag{67}$$

where we have explicitly written out the remainders $R,S$, which are bounded as $t \to 0$. In order to conclude (66), we need to show that derivatives $\frac{d^k}{d\theta^k}R(t,\theta)$, $\frac{d^k}{d\theta^k}S(t,\theta)$ exist and are bounded as $t \to 0$. The key to this lies in showing this property for the decoding function mapping a neural encoding $m$ to the estimate $\widehat{\theta}$. We set

$$f(t,z,y) := \frac{1}{\sqrt{2\pi t}} \int_{-\infty}^{\infty} Q(x,y) \exp\Big(-\frac{(x-z)^2}{2t}\Big)\, dx,$$

with $t,z \in [0,1]$ and

$$Q(x,y) := (y - F^{-1}(x))^{p-1} q_{prior}(x) \tag{68}$$

As shown in Hahn and Wei [5, SI Appendix, Proof of Theorem 1] (also taken up in our Section S2.6.3), the estimator $\widehat{\theta}$, as a function of $t$ and θ, is obtained as the implicit function solving

$$f(t, F(\theta), \widehat{\theta}) = 0 \tag{69}$$

We want to understand the regularity of $\widehat{\theta}$ as a function jointly for $t$ and θ, using the Implicit Function Theorem. A priori, due to division by $t$, $f$ might potentially be unsmooth at $t \to 0$, impeding applicability of that theorem. To analyze the regularity of $f$ as $t \to 0$, we perform the change of variables, rewriting the variable $x$ explicitly in relation to the neural encoding $z$:

$$x = z + \sqrt{t}\, u,$$

so that

$$dx = \sqrt{t}\, du.$$

Substituting into the definition of $f$ gives:

$$\begin{aligned} f(t,z,y) &= \frac{1}{\sqrt{2\pi t}} \int_{-\infty}^{\infty} Q(z + \sqrt{t}\, u, y) \exp\Big(-\frac{(z + \sqrt{t}\, u - z)^2}{2t}\Big) \sqrt{t}\, du \\ &= \int_{-\infty}^{\infty} Q(z + \sqrt{t}\, u, y) \phi(u)\, du, \end{aligned}$$

where

$$\phi(u) = \frac{1}{\sqrt{2\pi}} \exp\Big(-\frac{u^2}{2}\Big)$$

is the standard Gaussian density. We can expand $Q(z + \sqrt{t}\, u, y)$ in a Taylor approximation about $z$:

$$Q(z + \sqrt{t}\, u, y) = Q(z,y) + \sqrt{t}\, u\, Q_x(z,y) + \frac{t}{2} u^2\, Q_{xx}(z,y) + \frac{t^{3/2}}{6} u^3\, Q_{xxx}(z,y) + \cdots.$$

to as many orders as allowed by the regularity of $F^{-1}$ and $p_{prior}$. Because $\phi(u)$ is symmetric (i.e., $\int u\phi(u)\, du = 0$ and all odd moments vanish), the terms with odd powers of $u$ drop out when integrated. Thus, integrating term-by-term yields:

$$f(t,z,y) = Q(z,y) + \frac{t}{2} Q_{xx}(z,y) + \frac{t^2}{24} Q_{xxxx}(z,y) + \cdots$$

We further substitute $y = F^{-1}(z) + tR$:

$$f(t,z,F^{-1}(z)+tR) = Q(z,F^{-1}(z)+tR) + \frac{t}{2}Q_{xx}(z,F^{-1}(z)+tR) + \frac{t^2}{24}Q_{xxxx}(z,F^{-1}(z)+tR) + \cdots$$

Now performing a Taylor expansion in the $y$ direction, we obtain a polynomial in $R$, with coefficients that include different powers of $t$, with remainder bounded in terms of powers of both $t$ and $y$. If $p > 2$, many lower-order terms will be zero; we divide by as many powers of $t$ as needed to make the lowest-order term be of order $O(1)$. After this transformation, the resulting function will have a nonzero derivative w.r.t. $R$ at $t = 0$. As a consequence, by the Implicit Function Theorem, $\widehat{\theta}$ is jointly smooth in $t$ and $\theta$ to as many orders as allowed by the regularity of $F$ and $p_{prior}$. Now given the expansion

$$\widehat{\theta} = \theta + tR(\theta) + t^2 u(t,\theta) \tag{70}$$

where $u$ is bounded as $t \to \infty$, we want to show

$$\frac{d^k}{d\theta^k}\widehat{\theta} = \frac{d^k}{d\theta^k}\theta + t\frac{d^k}{d\theta^k}R(\theta) + O(t^2) \tag{71}$$

which is equivalent to saying that $\frac{d^k}{d\theta^k}u(t,\theta) = O(1)$ as $t \to 0$, for as many orders $k$ as required and allowed by $F$ and $p_{prior}$. To show this, we think of $u$ as an implicit function defined by

$$0 = g(t,\theta,u) := f(t,F(\theta),\theta + tR(\theta) + t^2 u) \tag{72}$$

Then the Implicit Function Theorem guarantees the existence of a smooth solution $u$ that has smooth (hence, bounded) derivatives in $\theta$ to the order allowed by $F$ and $p_{prior}$. This entails (71), and analogously for higher derivatives. Specifically, the proof of our Theorem 3 requires derivatives up to $F''''$ and $\frac{d^2}{d\theta^2}\log p_{prior}$ estimated by differentiating bias and variability up to three times; overall, we require up to the fourth derivatives of $F$ and $p_{prior}$. We note that, while in (71) we expanded $\widehat{\theta}$ only up to the first order, the same argument applies to an expansion to second order. As the overall mean response is a convolution of $\widehat{\theta}(F^{-1}(m))$ with a Gaussian distribution over $m$, the conclusion (71) is preserved for the expansion of the bias. An analogous argument applies to the variance. Overall, Eq. 66 follows. We note that the above reasoning applies to $p \geq 2$; a similar consideration applies to the MAP estimator, which is implicitly defined as a stationary point of the posterior and can likewise be handled by the implicit function theorem. For the L1 estimator, direct calculation via implicit differentiation can be performed.

#### S2.7 Restoring Identifiability via Adaptation

Our theoretical results (Theorems 1–4) reveal that full identification of prior, encoding, and loss function is usually possible when data from multiple levels of sensory noise is available; however, there is an exceptional set $\Omega$ of models where even multiple noise levels may not be sufficient, such that prior and loss function are confounded even when observations at multiple noise levels are available. The fundamental issue lies in the fact that, under such models, response data may be explained equally well by different loss functions.

How can one restore identifiability in such a situation? One avenue for recovering identifiability can be by inducing a short-term prior. While both prior and encoding may adapt to stimulus statistics, there is substantial evidence that they adapt at different timescales [2, 5], and can even be dissociated on a trial-by-trial basis [2]. It is thus quite likely that exposure to a nonuniform short-term prior that leads to different adaptation in $F$ and $p_{prior}$ will bring the encoding-decoding-process outside of the unidentifiable set $\Omega$, allowing unique identification of the loss function. Assuming that the loss function used by the subject remains the same before and after adaptation, one can then use this loss function to infer a prior even from data collected without adaptation.

Even if prior and encoding jointly adapted according to some fixed efficient coding rule (e.g., information maximization [26] or power-law encoding [12]), this would usually result in leaving $\Omega$, since the set of priors $p_{prior}$ where the encodings $F$ with are linked as $F' \propto p_{prior}^q$ itself has measure zero. Hence, one strategy to recover identifiability can be to induce different priors (and possibly encodings) across multiple experimental conditions. Under the assumption that the subjects still use the same loss function, one can expect the loss function to generally become identifiable.

We show this for two special cases (Theorems S26 and S27): when the encoding shows little or now adaptation (as suggested e.g. by the fit to the data of [3] in [5]), or when the encoding fully adapts (as assumed in work on efficient coding, [e.g. 26, 12]). This argument is quite general and also applies to more complex situations where prior and encoding adapt differently, even possibly at different timescales [2]. Importantly, this strategy *does not* rely on knowing the law of adaptation, it only relies on the presence of *some* systematic link adapting prior and encoding, which has an overwhelming chance of moving encoding and prior used by the participant out of the set $\Omega$.

Recall $L(\cdot)$, $\mathcal{F}(\mathcal{X})$, and $\mu$ from Sections S2.1.3–S2.1.4.

Our first result here concerns the situation where prior and encoding are exactly matched up to a power-law transformation, as suggested by theories of efficient coding [e.g. 26, 12]. Our result says that the measure of priors for which the resulting model will be in $\Omega$ is zero. That is, efficiently adapting prior and encoding to some environmental statistics is almost certain to result in identifiability:

**Theorem S26** (Full Adaptation of Prior and Encoding)**.** *Let $q > 0$. The set of priors $p_{prior} \in \mathcal{F}(X)$ such that*

$$\exists p \in \mathcal{P} : \langle F', p_{prior}, p \rangle \in \Omega, F' \propto p_{prior}^q \tag{73}$$

*has measure zero.*

*Proof.* Given a model

$$M := \langle F', p_{prior}, p \rangle \tag{74}$$

in $\Omega$ with $F' \propto p_{prior}^q$, there must be a model $M'$ with the same encoding, a different exponent $p'$, and an appropriately transformed prior, such that

$$\mathcal{D}_M = \mathcal{D}_{M'}$$

This is an ordinary differential equation in the function $L(p_{prior})$. By the same arguments as in the proof of Lemma S17, the solution set has measure zero. □

Our second result here investigates the case where the encoding stays constant, and the prior adapts to reflect some environmental statistics. Again, this will almost surely result in identifiability:

**Theorem S27** (Prior adapts, encoding stays constant)**.** *For any given encoding $F$, the set of priors $p_{prior}$ such that*

$$\exists p \in \mathcal{P} : \langle F', p_{prior}, p \rangle \in \Omega \tag{75}$$

*has measure zero.*

*Proof.* Similar to the previous theorem, for any fixed $F$, we obtain an ordinary differential equation for $p_{prior}$. The solution set has measure zero again. □

### S3 Implementation

#### S3.1 Fitting Procedure

We used the implementation of the fitting procedure from Hahn and Wei [5]. It fits the model parameters to maximize data likelihood with gradient descent. As described in Hahn and Wei [5], $F'$ and $p_{prior}$ are defined by assigning one unconstrained real number to each point on the discretized stimulus space grid $x_1, \dots, x_N$, and obtaining $F'(\theta), p_{prior}(\theta)$ via the softmax transform (matching the theoretical construction of $\mu$ on $\mathcal{F}(\mathcal{X})$ in Section S2.1.3). A key aspect of this implementation is that it allows the backpropagation algorithm to compute gradients of $\widehat{\theta}$ with respect to the posterior $P$.

The implementation of the $L_p$ ($p = 2, 4, 6, \dots$) estimators is unchanged. We add an implementation of the $L_1$ estimator (posterior median), explained in Section S3.2. For the MAP estimator, we performed two changes. As described in Hahn and Wei [5], the MAP estimator is obtained as the maximum of function $\tilde{P}$ obtained by smoothing the discretized posterior with a Gaussian function. First, for circular spaces, we used a von Mises density function instead of a Gaussian function. Second, on interval stimulus spaces, a challenge is that the maximum of the smoothed posterior might lie outside the stimulus space, in fact, it may lie at $\pm\infty$, and the need to compute $\frac{\partial\widehat{\theta}}{\partial P(x_i)}$ for gradient-based fitting precludes simply truncating it at the boundary (which would be a nondifferentiable operation). Hahn and Wei [5] solved this by adding a smooth function that is almost constant inside the space and rapidly decays to $-\infty$ outside it, but we found this to make numerical fitting challenging when there are datapoints close to the boundary. We instead simply set the discretized posterior at the boundary points to be zero ($P(x_0), P(x_N) := 0$), which naturally ensures that the maximum of the smoothed posterior $\widehat{P}$ will be attained at a finite $\widehat{\theta}$. This $\widehat{\theta}$ might lie slightly outside the stimulus space; though the motor likelihood is restricted to points inside the stimulus space.

On interval stimulus spaces, the procedure of Hahn and Wei [5] computes the normalization constant of the motor distribution (i.e., the integral of a Gaussian density over an interval) numerically by summing over a grid; imprecision of this computation however can lead to incorrect model likelihoods when motor variance is very small. We found this to cause problems (i.e., spuriously low losses) at $p = 0$ and (to a lesser extent) $p = 1$, but not larger exponents. Hence, at $p = 0, 1$ on interval stimulus spaces, we rounded the data to the grid points $x_1, \ldots, x_N$, and converted the motor likelihood to a discrete distribution over those, guaranteeing a correct normalization constant. No changes are needed on circular stimulus spaces, where the normalization constant is independent of $\widehat{\theta}$.

#### S3.2 $L_1$ estimator (Posterior Median)

Whereas Hahn and Wei [5] implemented the $L_p$ estimators at even exponents ($p = 0, 2, 4, 6, \ldots$), we extended their fitting procedure to cover the $L_1$ estimator, i.e., the Posterior Median, as it is a relatively popular choice. Here, we discuss its definition and implementation.

##### Interval Stimulus Space

We first discuss the setting where $\mathcal{X}$ is an interval. Given a neural encoding $m \in \mathcal{Y}$ and the corresponding posterior $P(x|m)$ over $\mathcal{X}$, the $L_1$ loss for an estimate $\theta \in \mathcal{X}$ is defined as

$$\int |x-\theta| P(x|m) dx \tag{76}$$

In the implementation of Hahn and Wei [5], we are given the discretized posterior $P(x_1|m), \ldots, P(x_N|m)$ based on the discretization $x_1, ..., x_N$ of the stimulus space $\mathcal{X}$ ($x_{i+1} - x_i = \Delta$).

Whereas $L_p$ losses at $p \geq 2$ can be straightforwardly operationalized by replacing the integral with a sum over the discretized points $x_i$, the situation is more complex at $p = 1$, where a discretized version of (76) results in the median of a discrete distribution, which (depending on its definition) either is not unique or does not smoothly depend on $P$. Hence, we interpolate $P(\cdot|m)$ as a function $\widehat{P}$ defined on all of $\mathcal{X}$, and then define the median on that distribution. To obtain a well-defined median, a very simple piecewise constant interpolation of $P$ is sufficient. Let $(\phi_i)_{i=1,\ldots,N}$ be a sequence of functions $\mathcal{X} \to \mathbb{R}$, such that we interpolate

$$\widehat{P}(x) := \sum_{i=1}^{N} P(x_i|m)\phi_i(x) \tag{77}$$

It is sufficient to take

$$\phi_i(x) = \begin{cases} 1 & \text{if } |x-x_i| < |\frac{\Delta}{2}| \\ 0 & \text{else} \end{cases} \tag{78}$$

Then $\widehat{P}$ is piecewise constant, and interpolates $P(\cdot|m)$. We use it to define the implemented $L_1$ estimator:

**Definition S28.** *We define the $L_1$ estimator (median) as*

$$\min_{\theta\in[0,1]} \int_{\mathcal{X}} |\theta - x| \widehat{P}(x) dx \tag{79}$$

To determine it, we solve for $\widehat{\theta}$ satisfying:

$$0 = \int_{\mathcal{X}} sign(\widehat{\theta} - x)\widehat{P}(x)dx, \qquad \text{or equivalently:} \qquad \int_{x<\theta} \widehat{P}(x)dx = \int_{x>\theta} \widehat{P}(x)dx \tag{80}$$

As $P(\cdot|m)$ and hence $\widehat{P}$ is always strictly positive[11], there will be a unique solution $x_a$. Let $x_a$ be one of $x_1, \dots, x_N$. Set

$$A = P(x_a) \qquad C = \sum_{j:x_j<x_a} P(x_j) \qquad D = \sum_{j:x_j>x_a} P(x_j)$$

We now want to choose $\alpha \in [-\frac{1}{2}, \frac{1}{2}]$ such that $\widehat{\theta} = x_a + \alpha\Delta$. Then, $\widehat{\theta}$ partitions the patch $[x_a - \frac{\Delta}{2}, x_a + \frac{\Delta}{2}]$ with length $\Delta$ into sub-patches of length $(\frac{1}{2} + \alpha)\Delta$ (on the left) and $(\frac{1}{2} - \alpha)\Delta$ (on the right). The overall mass on the left of $\widehat{\theta}$ is

$$\int_{x<\theta} sign(\widehat{\theta} - x)\widehat{P}(x)dx = \Delta D + \left(\frac{1}{2} + \alpha\right)\Delta A \tag{81}$$

The overall mass on the right of $\widehat{\theta}$ is

$$\int_{x>\theta} sign(\widehat{\theta} - x)\widehat{P}(x)dx = \Delta C + \left(\frac{1}{2} - \alpha\right)\Delta A \tag{82}$$

Setting these two equal results in

$$\frac{C-D}{2A} = \alpha \tag{83}$$

We try all $x_a$'s, and choose the one that ensures $\alpha \in [-\frac{1}{2}, \frac{1}{2}]$ while minimizing (79). An important feature of the fitting procedure of Hahn and Wei [5] is that allows backpropagation of gradients through $\widehat{\theta}$ to the posterior $P(x|m)$, allowing full gradient-based fitting. In order to backpropagate through $\widehat{\theta}$, we need its derivatives w.r.t. $P(x_i)$. We can simply differentiate Eq. 83.

$$\frac{\partial \alpha}{\partial P(x_i)} = \begin{cases} \frac{1}{2A} & x_i - x_a > 0 \\ -\frac{1}{2A} & x_i - x_a < 0 \\ \frac{D-C}{2A^2} & i = a \\ -\frac{D-C}{2A^2} & i = b \end{cases}$$

Then

$$\frac{\partial \widehat{\theta}}{\partial P(x_i)} = \frac{\partial \alpha}{\partial P(x_i)} \cdot \Delta \tag{84}$$

**Circular Stimulus Space**

We now consider the case where $\mathcal{X}$ is circular. For the standard definition of the circular median see, we refer to Otieno and Anderson-Cook [14]. Essentially, the circular median is defined as a point partitioning the circle into two semicircles that carry equal mass of $P$. There are always at least two antipodal points satisfying this definition; the one closer to most mass of $P$ is chosen.

As before, we interpolate the posterior via a piecewise constant function. Let $(\phi_i)_{i=1,\dots,N}$ be a sequence of functions $\mathcal{X} \to \mathbb{R}$, such that we interpolate

$$\widehat{P}(x) := \sum_{i=1}^{N} P(x_i)\phi_i(x) \tag{85}$$

We take

$$\phi_i(x) = \begin{cases} 1 & \text{if } \cos(x - x_i) > \cos\left(\frac{\Delta}{2}\right) \\ 0 & \text{else} \end{cases} \tag{86}$$

[11] This holds because we assume the prior to never be strictly zero, and the Fisher information to always be finite (Section S2.1.1).

We first set

$$\ell^1(x,y) := \min\{|x-y+k2\pi| : k \in \mathbb{Z}\} = \arccos(\cos(x-y))$$

where arccos is the usual principal value (mapping to $[0,\pi]$). This function describes the arc-length distance between two points on the unit circle. Then we formally define:

**Definition S29.** *For circular $X$, we define the circular $L_1$ estimator (median) as*

$$\arg\min_{\theta\in[0,2\pi)} \int_0^{2\pi} \ell^1(\theta,x)\widehat{P}(x)dx \tag{87}$$

We note that this, unlike interval stimulus spaces, the median may sometimes be multi-valued on circular spaces; we address this below. We connect this definition to the definition of the circular median from Otieno and Anderson-Cook [14] as follows:

**Lemma S30.** *Let $\widehat{\theta}$ be a solution to (87), and $\widehat{\theta}^\dagger$ its antipode (that is, $\widehat{\theta}^\dagger = (\widehat{\theta}+\pi)\%(2\pi)$). Then the diameter $\widehat{\theta} : \widehat{\theta}^\dagger$ divides the circle into two semicircles that carry equal mass of $\widehat{P}$.*

*Proof.* We first note that the derivative of the distance function is given by the following function, a circular analogue of the sign function, defined, for $x \in \mathbb{R}$, as:

$$circsign(x) := sign(\sin(x)) = \begin{cases} 1 & x \in \ldots,(-2\pi,-\pi),(0,\pi),(2\pi,3\pi),\ldots \\ 0 & x \equiv 0(\operatorname{mod}\pi) \\ -1 & x \in \ldots,(-\pi,0),(\pi,2\pi),(3\pi,4\pi),\ldots \end{cases}$$

Then:

$$\partial_x \ell^1(x,y) = circsign(x-y)$$

To show this, without loss of generality, it is sufficient to show the claim at $y=0$. It then follows from the fact

$$\frac{d}{dx} arccos(\cos(x)) = sign(\sin(x))$$

which can be shown by case distinction. Now, by differentiating Eq. 87, we obtain:

$$0 = \int circsign(\widehat{\theta}-x)\widehat{P}(x)dx$$

This is equivalent to stating that the mass of $\widehat{P}$ is equal on both semicircles. □

Let $x_a$ be a point among $x_1,\ldots,x_N$, and let $x_b$ be its antipode. Set

$$A = P(x_a) \qquad B = P(x_b) \qquad C = \sum_{j:\sin(x_j-x_a)>0} P(x_j) \qquad D = \sum_{j:\sin(x_j-x_a)<0} P(x_j)$$

We now want to choose $\alpha \in [-\frac{1}{2},\frac{1}{2}]$ such that $\widehat{\theta} = x_a + \alpha\Delta$. Then, among the patch $[x_a - \frac{\Delta}{2}, x_a + \frac{\Delta}{2}]$ with length $\Delta$, it is partitioned into sub-patches of length $(\frac{1}{2}+\alpha)\Delta$ (on the left) and $(\frac{1}{2}-\alpha)\Delta$ (on the right). The overall mass on the left of $\widehat{\theta}$ is

$$\int_{x:sin(x-\widehat{\theta})<0} \widehat{P}(x)dx = \Delta D + \left(\frac{1}{2}+\alpha\right)\Delta A + \left(\frac{1}{2}-\alpha\right)\Delta B \tag{88}$$

The overall mass on the right of $\widehat{\theta}$ is

$$\int_{x:sin(x-\widehat{\theta})>0} \widehat{P}(x)dx = \Delta C + \left(\frac{1}{2}-\alpha\right)\Delta A + \left(\frac{1}{2}+\alpha\right)\Delta B \tag{89}$$

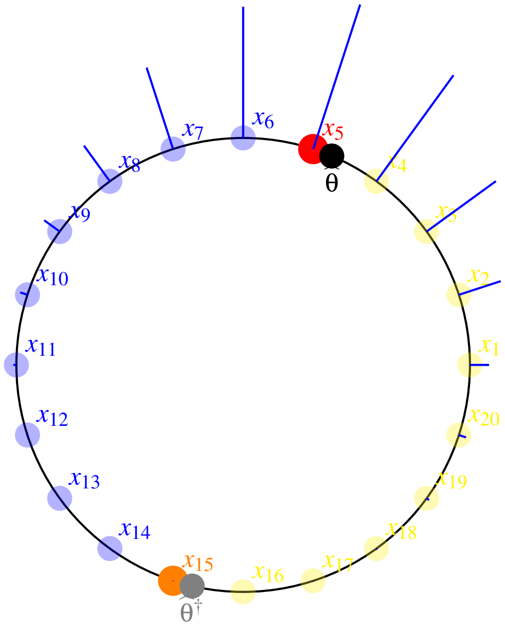

Figure S1: Illustration of the definition of the $L_1$ estimator on a discretization of the circle. We show a discretized distribution with a histogram with blue bars. The points $x_5$ and $x_{15}$ are highlighted in red and orange, respectively. The circular median $\widehat{\theta}$ is shown in black, and its antipode $\widehat{\theta}^\dagger$ in gray. The semicircles between $\widehat{\theta}$ and $\widehat{\theta}^\dagger$ carry equal weight.

Setting these two equal results in

$$\frac{D-C}{2B-2A} = \alpha \tag{90}$$

Among all $x_a$'s, we choose the one that ensures $\alpha \in [-\frac{1}{2}, \frac{1}{2}]$ while minimizing the $L_1$ loss. There are cases where the circular median is not unique; in particular, when $\widehat{P}$ is uniform or has very wide spread. [12] In general, however, the circular median is well-defined on any localized probability distribution, as is generally relevant to Bayesian modeling of perception, where (assuming the stimulus has been perceived and encoded at all) the posterior will be localized in a region of stimulus space. Importantly, the median on a nowhere-zero probability density $\widehat{P}$ is generally well-behaved and unambiguous, in contrast to the more challenging circular median of a finite number of datapoints (see Otieno and Anderson-Cook [14] for discussion of the latter).[13]

[12] For instance, when $\widehat{P}$ is uniform (not localized) or when it is proportional to $\sin^2(x)$ (which has two modes at antipodal points). For,

$$\int_{\widehat{\theta}}^{\widehat{\theta}+2\pi} \mathrm{sign}(\sin\left(x-\widehat{\theta}\right))\, \sin^2(x)\, dx$$

evaluates to 0, independent of $\widehat{\theta}$, because

$$\int_{\widehat{\theta}}^{\widehat{\theta}+\pi} \sin^2(x)\, dx = \frac{\pi}{2}$$

for any $\widehat{\theta}$.

[13] In the rare case that multiple candidate solutions nonetheless yield the same minimal $L_1$ loss, we employ a tie-breaking strategy that selects the solution closest to the overall probability mass. Specifically, for each candidate $\widehat{\theta}_i$ that minimizes the $L_1$ loss, we compute

$$\int (1-\cos\left(x-\widehat{\theta}_i\right))\widehat{P}(x)dx$$

Among all candidate solutions with optimal $L_1$ loss, the candidate minimizing this term is chosen as the final estimator.

As before, we obtain the gradient by differentiating Eq. 90.

$$\frac{\partial \alpha}{\partial P(x_i)} = \begin{cases} \frac{-1}{2B-2A} & \sin(x_i - x_a) > 0 \\ \frac{1}{2B-2A} & \sin(x_i - x_a) < 0 \\ \frac{(D-C)}{2(B-A)^2} & i = a \\ -\frac{(D-C)}{2(B-A)^2} & i = b \end{cases}$$

Then

$$\frac{\partial \widehat{\theta}}{\partial P(x_i)} = \frac{\partial \alpha}{\partial P(x_i)} \cdot \Delta \tag{91}$$

Again, implicit differentiation would lead to the same result.

#### S3.3 Computation of Encoding Resources

In general, with Gaussian encoding noise, the Fisher Information is given as

$$\mathcal{J}(\theta) = \frac{F'(\theta)^2}{\sigma^2} \tag{92}$$

We compute the plotted resource allocation $\sqrt{\mathcal{J}(\theta)}$ as in [5]. Here, we recapitulate the calculation for completeness. At $i = 1, \dots, N-1$, write:

$$V_i := F(\theta_{i+1}) - F(\theta_i) \tag{93}$$

so that

$$F'(\theta_i) \approx \frac{F(\theta_{i+1}) - F(\theta_i)}{\theta_{i+1} - \theta_i} = \frac{V_i N}{\mathrm{Vol}(\mathcal{X})} \tag{94}$$

For an interval stimulus space, we obtain

$$\sqrt{\mathcal{J}(\theta_i)} = \frac{F'(\theta_i)}{SD(m|\theta_i)} \approx \frac{V_i N}{\sigma \cdot \mathrm{Vol}(\mathcal{X})} \tag{95}$$

For a circular variable where sensory noise has von Mises parameter $\kappa$, we analogously take

$$\frac{V_i \sqrt{\kappa} N}{\mathrm{Vol}(\mathcal{X})} \tag{96}$$

We note that, at large noise, the standard definition of the Fisher information results in an additional term involving Bessel functions, which is close to 1 when noise is small. We follow [5] in disregarding it in visualization, in order to maintain the direct functional relationship between Fisher information and encoding slope (i.e., $\mathcal{J}(\theta) = \frac{F'(\theta)^2}{\sigma^2}$).

We note that in our simulations for circular stimulus spaces, we assume an orientation perception setup. While the stimulus space is represented as [0,360] in the implementation, we convert it to [0,180] for plotting purposes. In this transformation, bias and variability are divided by 2, whereas the resources $\sqrt{\mathcal{J}(\theta)}$ are multiplied by 2.

We compare three numerical methods of computing or approximating the Fisher information in Figure S2:

1. As described above: for a circular stimulus space

$$\sqrt{\mathcal{J}(\theta_i)} \approx \frac{V_i \sqrt{\kappa} N}{\mathrm{Vol}(\mathcal{X})} \tag{97}$$

and, for an interval stimulus space:

$$\sqrt{\mathcal{J}(\theta_i)} \approx \frac{V_i N}{\sigma \cdot \mathrm{Vol}(\mathcal{X})} \tag{98}$$

2. A finite differences approximation of the result of Theorem 1[14],

$$\sqrt{\mathcal{J}(\theta)} = \sqrt{4\pi}\frac{d}{d\theta}\mathbb{P}(m(\theta+h) \geq m(\theta)) \tag{99}$$

where $m(\theta)$ is an encoding sampled for stimulus $\theta$, operationalized on a circular stimulus space as

$$\sqrt{\mathcal{J}(\theta)} = \sqrt{4\pi}\frac{\mathbb{P}(\sin(m(\theta_{i+1}) - m(\theta_i) > 0) + \frac{1}{2}\mathbb{P}(m(\theta_{i+1}) = m(\theta_i)) - 0.5}{\theta_{i+1} - \theta_i} \tag{100}$$

and on an interval stimulus space as

$$\sqrt{\mathcal{J}(\theta)} = \sqrt{4\pi}\frac{\mathbb{P}(m(\theta_{i+1}) > m(\theta_i)) + \frac{1}{2}\mathbb{P}(m(\theta_{i+1}) = m(\theta_i)) - 0.5}{\theta_{i+1} - \theta_i} \tag{101}$$

where $m$ is discretely distributed over $F(\theta_1), \ldots, F(\theta_N)$ as specified in the implementation.

3. The general definition of the Fisher information

$$\mathcal{J}(\theta) = \int_{\mathcal{Y}} \left(\frac{\partial}{\partial\theta}\log p(m|\theta)\right)^2 p(m|\theta)dm \tag{102}$$

where the derivative is evaluated by a finite-difference approximation, and the likelihood is evaluated over the discrete grid.

4. The inverse variance of the encoding transformed back into stimulus space:

$$\frac{1}{\mathcal{J}(\theta)} \approx \mathbb{E}_{m|\theta}\left\{\left(F^{-1}(m) - \mathbb{E}_{m|\theta}[F^{-1}(m)]\right)^2\right\} \tag{103}$$

which is valid when noise is small.

Results in Figure S2 show that the methods result in numerically similar results. The third operationalization results in slightly smoothed estimates due to the finite-difference and low-noise approximations, which break exact analytic equivalence to the other expressions. Similarly, the fourth operationalization results in more smoothed estimates; it is a valid approximation only when noise is small.

[14]Theorem 1 differs by applying the decoding function to obtain $\widehat{\theta}$. These two variants are equivalent, since the decoding function is monotonic.

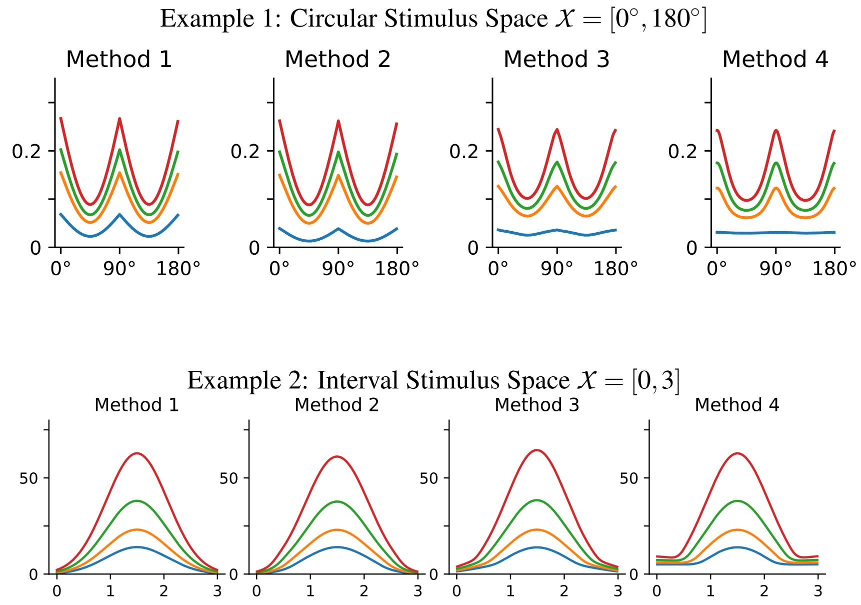

Figure S2: Comparing methods of computing or approximating the resource allocation in the implementation, as described in Section S3.3 for a circular (top) or interval (bottom) stimulus space. Method 1 is used throughout plots in this paper, as in [5], as it directly reflects the general correspondence between the Fisher information and the slope $F'$ of the encoding function. The other methods are equivalent when noise is small, though the correspondence becomes approximative when noise is larger.

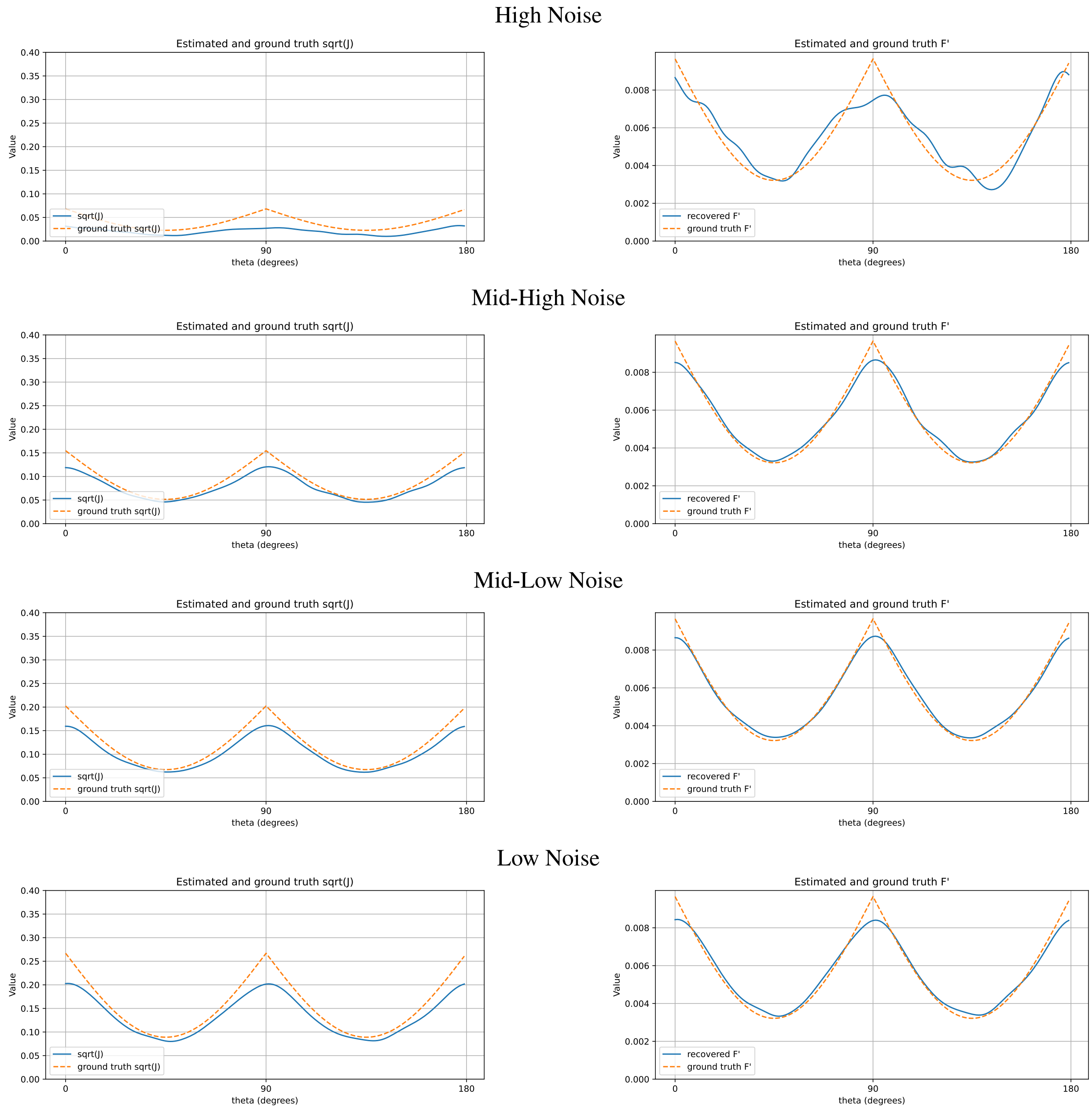

Figure S3: Here, we provide an example of how to recover the encoding $\sqrt{J}(\theta)$ from simulated data, using Theorem 1. Recovering the encoding using Theorem 1, at four different levels of sensory noise. 40K trials were simulated, separately at each noise level, with $p = 2$ and a periodic prior. We compute the empirical probability $P = P\left(\hat{\theta}(\theta + \frac{h}{2}) > \hat{\theta}(\theta - \frac{h}{2})\right)$ and then put $\sqrt{J}(\theta) = \frac{\left(P-0.5\right)\sqrt{4\pi}}{h\Delta\theta}$ with $h = 2^\circ$ (of the orientation space $[0^\circ, 180^\circ]$) as an estimate of the derivative. The result is convolved with a Gaussian kernel with $\sigma = 6^\circ$.

### S4 Further Simulations Results

We provide extensive simulations for a variety of models on circular and interval spaces, expanding on Figures 4 and 5 in the main text.

#### S4.1 Operationalization of Attraction and Repulsion

Throughout the SI Appendix, we visualize not only the overall fitted bias, but also its decomposition into Attraction and Repulsion components. As shown in [5], the bias for the $L_p$ estimator has the form

$$\mathrm{Bias}(\theta) = \underbrace{\frac{1}{\mathcal{J}(\theta)}(\log p_{prior}(\theta))'}_{\text{Prior Attraction}} + \underbrace{\frac{p+2}{4}\left(\frac{1}{\mathcal{J}(\theta)}\right)'}_{\text{Likelihood Repulsion}} + O(\sigma^4) \tag{104}$$

when $p > 0$, and

$$\mathrm{Bias}(\theta) = \underbrace{\frac{1}{\mathcal{J}(\theta)}(\log p_{prior}(\theta))'}_{\text{Prior Attraction}} + \underbrace{\frac{1}{4}\left(\frac{1}{\mathcal{J}(\theta)}\right)'}_{\text{Likelihood Repulsion}} + O(\sigma^4) \tag{105}$$

in the case $p = 0$.

The first component expresses classical attraction to the prior, traditionally considered a hallmark of Bayesian models. The second component expresses repulsion away from regions allocated high encoding resources, towards stimuli that are less well encoded; it appears because such inhomogeneities in encoding precision make the posterior asymmetric [28, 26, 27, 18, 5].

This decomposition of the bias clarifies how prior, encoding, and loss function jointly determine the bias. Our theoretical and simulation results indicate that the decomposition can be reliably recovered from behavioral data. To illustrate this, thoughout the SI Appendix, we visualize not only the fitted overall bias, but also the attraction and repulsion components. As the decomposition is exact only in the low-noise limit (indicated by the $O(\sigma^4)$ residual term), we require an operationalization of the decomposition that is adapted to larger noise levels. To achieve this, we follow [5] in operationalizing the attraction component as the decoding bias ($\mathbb{E}[\widehat{\theta}|m] - F^{-1}(m)$) for the MAP estimator ($p = 0$), which has the analytical form

$$\frac{1}{\mathcal{J}(\theta)}(\log p_{prior}(\theta))' + O(\sigma^4) \tag{106}$$

matching the Prior Attraction up to a higher-order residual. Also following [5], we operationalize the repulsion component as the difference between the overall prior and this component. By definition, and in accordance with the analytical decomposition (104–105), the attractive component is independent of the loss function, whereas the repulsive component depends on it. In the special case where encoding is exactly uniform, we instead plot the full bias as attraction, with a zero repulsive component (following [5]; Figure S7). As in [5], separate visualization of these components via such an operationalization serves an illustrative purpose, but we caution that there may be other possible operationalizations of the components in the finite-noise regime.

**Difference between Analytical and Plotted Components** We note that, while this operationalization provides a general idea of the decomposition, it reflects higher-order information of the residual $O(\sigma^4)$ and thus sometimes shows behavior not expected of the two components in the analytical decomposition (104–105). Specifically, the plotted repulsive component may somewhat depend on the prior because, as noise becomes larger, the prior can impact the higher-order residual $O(\sigma^4)$. As a consequence, even if the encoding is near-uniform and the repulsion is zero in the small-noise limit, the plotted repulsive component at large noise need not be zero.

**Behavior in Interval Stimulus Spaces** In interval stimulus spaces, close to the boundary, the bias receives an additional term, indicating a regression effect into the interior of the interval (Theorem 3 in [5]). The operationalization of attraction and repulsion described above naturally partitions this regression effect into components grouped with attraction and repulsion, respectively. Thus, rather than separately plotting the regression effect as a third component, we plot attraction and repulsion components that both include portions of the boundary effect, as determined by the operationalization.

#### S4.2 Models on Circular Spaces (Supplement to Figure 5)

For circular spaces, we investigate the following combinations:

1. Periodic encoding and uniform prior (Figures S4 and S8)
2. Periodic encoding and periodic (shifted) prior (Figures S5 and S9)
3. Periodic encoding and periodic prior (Figures S6 and S10)
4. Uniform encoding and periodic prior (Figures S7 and S11)

Across settings, we find that the encoding and prior are closely fitted when given the correct loss function, and that the loss function is closely identified when the number of noise levels and trials is sufficiently large. Further, in Figure S12, we provide an explicit example of models with different loss functions and priors that have the same response distribution in the limit of zero noise.

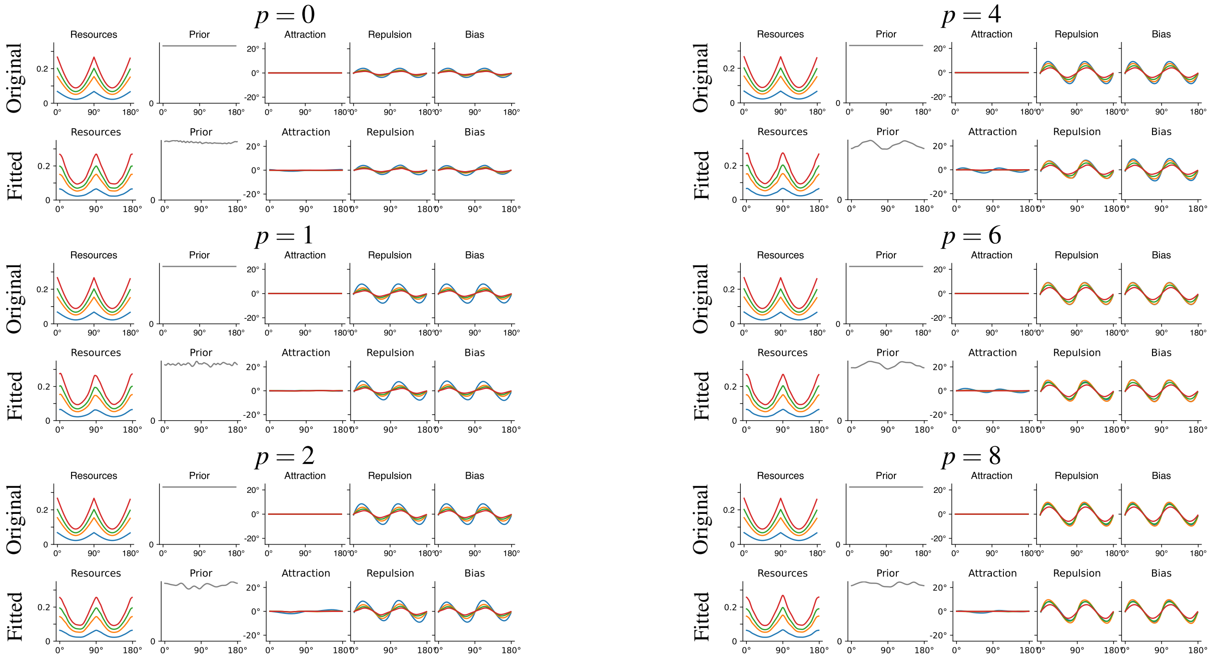

Figure S4: Circular Stimulus Space: Identifiability of encoding and prior (40K trials, 4 levels of sensory noise). For each loss function exponent, we show the ground-truth model at that exponent, and a fit when assuming this exponent. See Section S4.1 for discussion of Attraction and Repulsion components. Importantly, across the different ground truth exponents, encoding resources and prior are well recovered when the loss function is known.

Further, see Figure S13 for recoverability of these four models at $p = 2$, specifically at one or four levels of sensory noise. Further, see Figure S14 for results at one or two noise levels, broken down by pairs of noise levels.

#### S4.3 Models on Interval Spaces

For interval spaces, we investigate the following combinations:

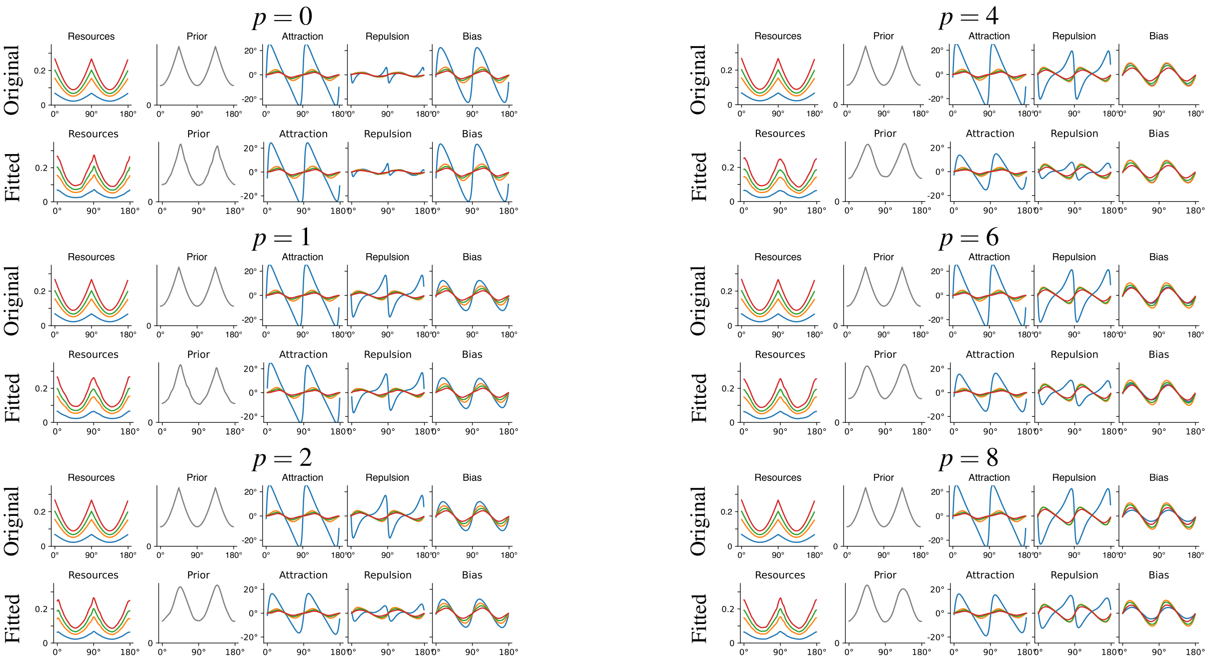

Figure S5: Circular Stimulus Space: Identifiability of encoding and prior (40K trials, 4 levels of sensory noise). For each loss function exponent, we show the ground-truth model at that exponent, and a fit when assuming this exponent. See Section S4.1 for discussion of Attraction and Repulsion components. Importantly, across the different ground truth exponents, encoding resources and prior are well recovered when the loss function is known.

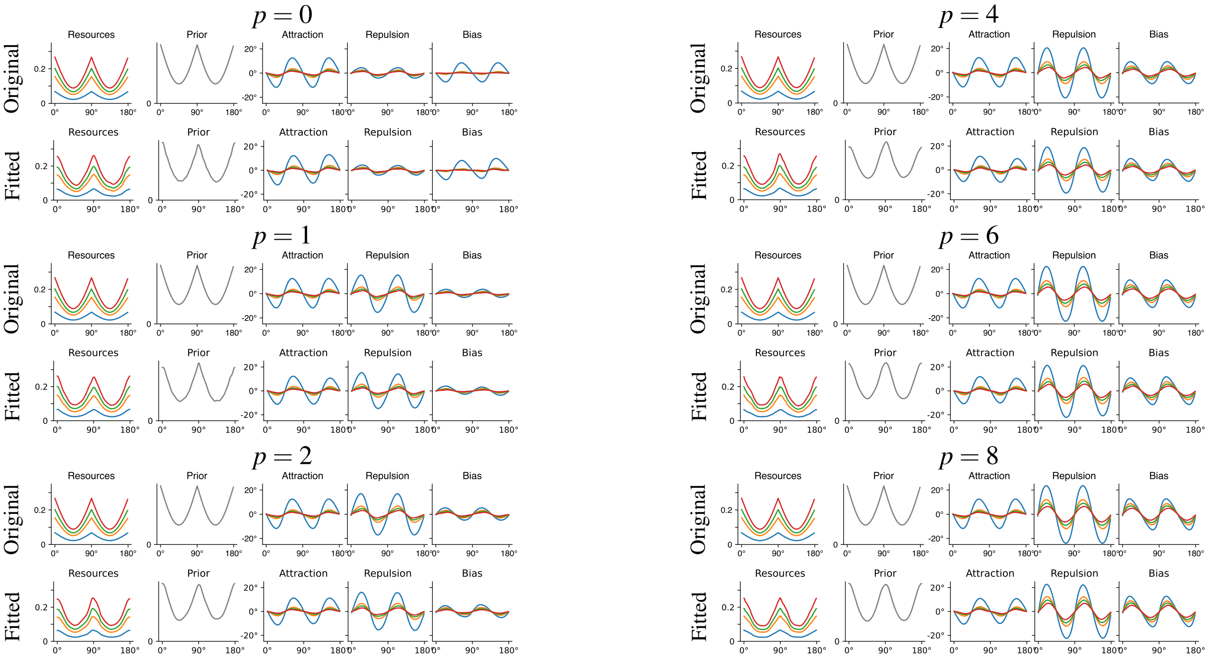

Figure S6: Circular Stimulus Space: Identifiability of encoding and prior (40K trials, 4 levels of sensory noise). For each loss function exponent, we show the ground-truth model at that exponent, and a fit when assuming this exponent. See Section S4.1 for discussion of Attraction and Repulsion components. Importantly, across the different ground truth exponents, encoding resources and prior are well recovered when the loss function is known.

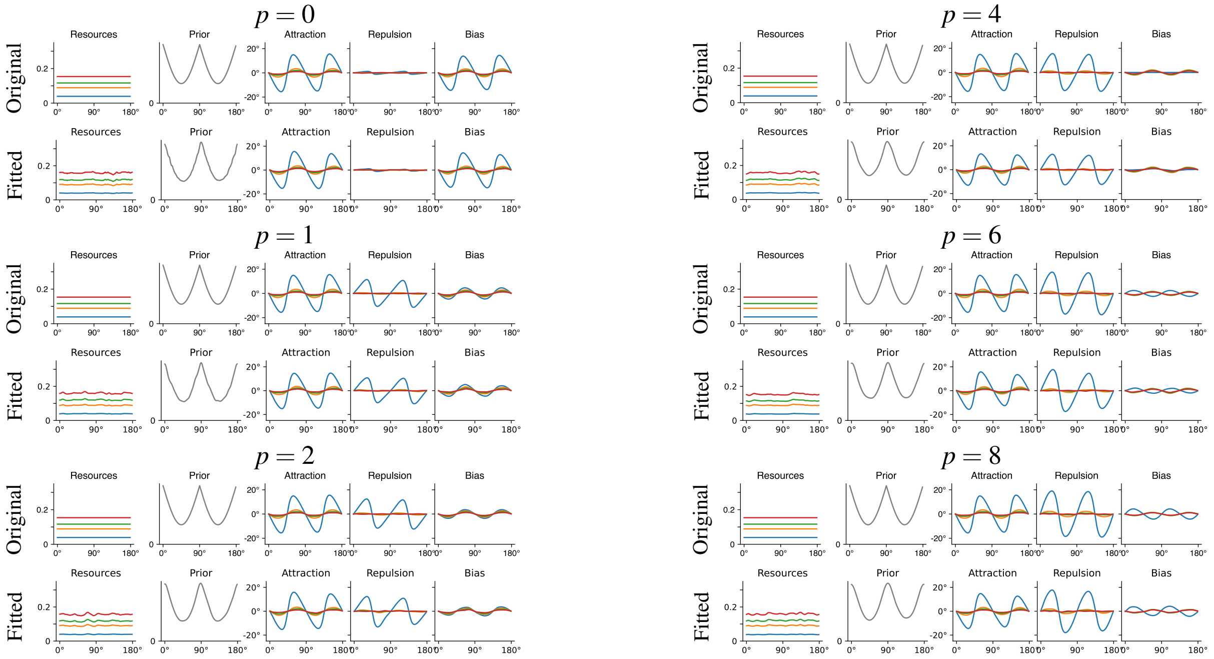

Figure S7: Circular Stimulus Space: Identifiability of encoding and prior (40K trials, 4 levels of sensory noise). For each loss function exponent, we show the ground-truth model at that exponent, and a fit when assuming this exponent. See Section S4.1 for discussion of Attraction and Repulsion components. Importantly, across the different ground truth exponents, encoding resources and prior are well recovered when the loss function is known.

1. Unimodal encoding and uniform prior (Figures S15 and S19)
2. Unimodal encoding and unimodal prior (Figures S16 and S20)
3. Bimodal encoding and uniform prior (Figures S17 and S21)
4. Uniform encoding and unimodal prior (Figures S18 and S22)

Across settings, we find that the encoding and prior are closely fitted when given the correct loss function, and that the loss function is closely identified when the number of noise levels and trials is sufficiently large.

#### S4.4 Randomly Generated Models (Supplement to Figure 4)

We provide additional extensive simulations complementing Main Text Figure 4 in Figures S23–S28, with three randomly generated model at each loss function exponent. Across the board, encodings, priors, and loss functions are recovered.

**Random Model Generation** For each of the random models, we initialized Python's `random` generator with a three-digit seed determined as follows: The first digit is 1 for the prior and 2 for the encoding. The second digit is the loss function. The third digit is 1, 2, 3, giving us a total of three random models per loss function exponent.

We then sampled the encoding and prior each by setting

$$\sum_{i=0}^{4} \left(X_i \sin(i\theta_i) + Y_i \cos(i\theta_i)\right) \tag{107}$$

where $X_i, Y_i \sim Uniform([-0.5, 0.5])$ drawn using the `random()` function, and applying the softmax function to the resulting vector to obtain an element of $\mathcal{F}(\mathcal{X})$.

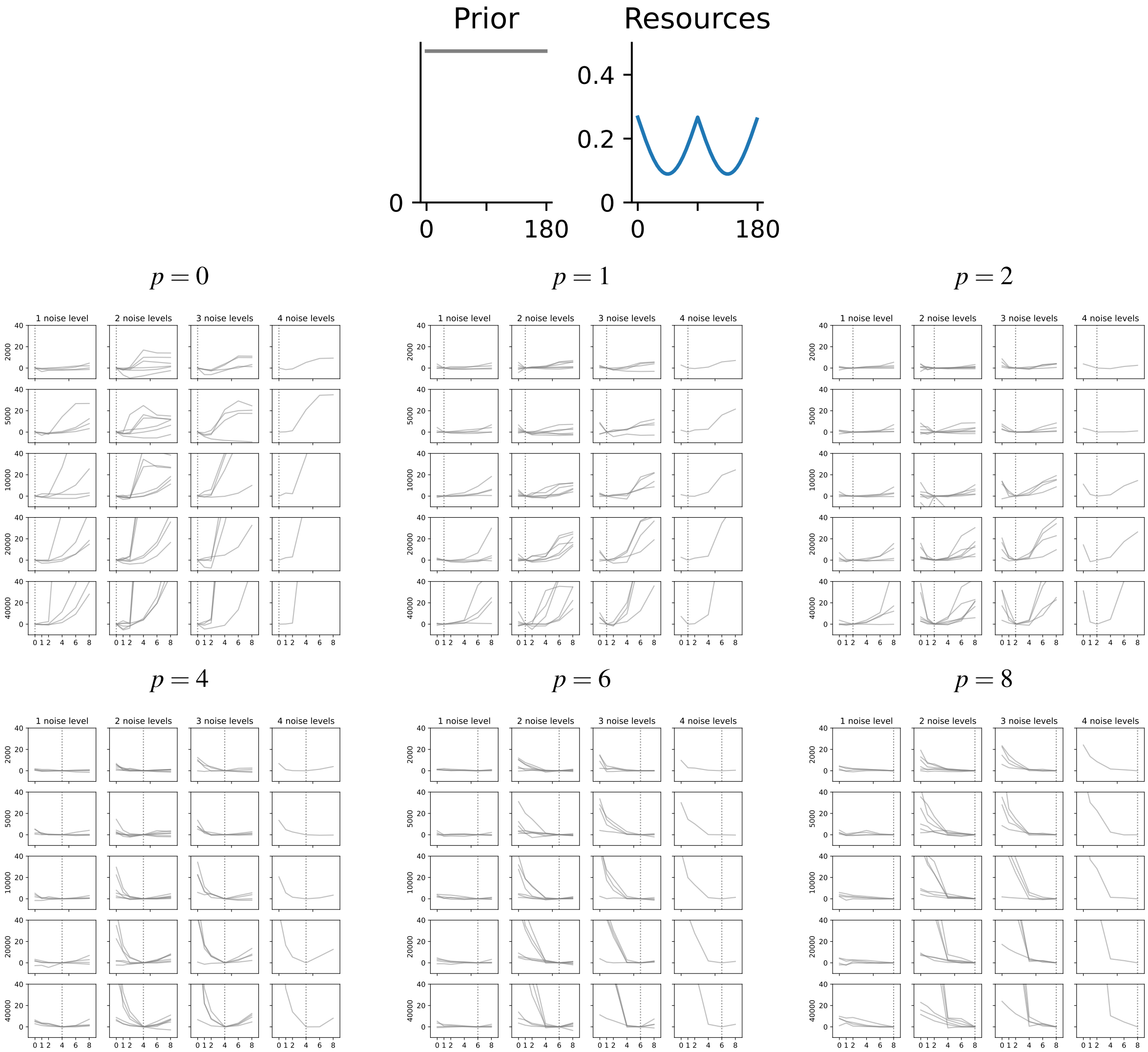

Figure S8: Circular Stimulus Space: Identifiability of the loss function depending on the number of trials (rows, from 2K to 40K), the number of of sensory noise levels (columns, from 1 to 4), and the ground truth loss function exponent (facets, $p$ from 0 to 8). Each curve represents one synthetic dataset with the given number of trials and noise levels, with the prior and encoding resources shown at the top. In each case, we include all possible combinations of the four overall noise levels, i.e., there are four cases at 1 noise level, $\binom{4}{2} = 6$ combinations at 2 noise levels, $\binom{4}{3} = 4$ combinations at 3 noise levels, and a single combination at 4 noise levels. We plot the model fit at each exponent as quantified by the difference to model fit at the ground-truth exponent, whose position is indicated with a dotted vertical line. Note that, in order to keep the y-axis range consistent across panels for easy comparison, values greater than 40 are not shown.

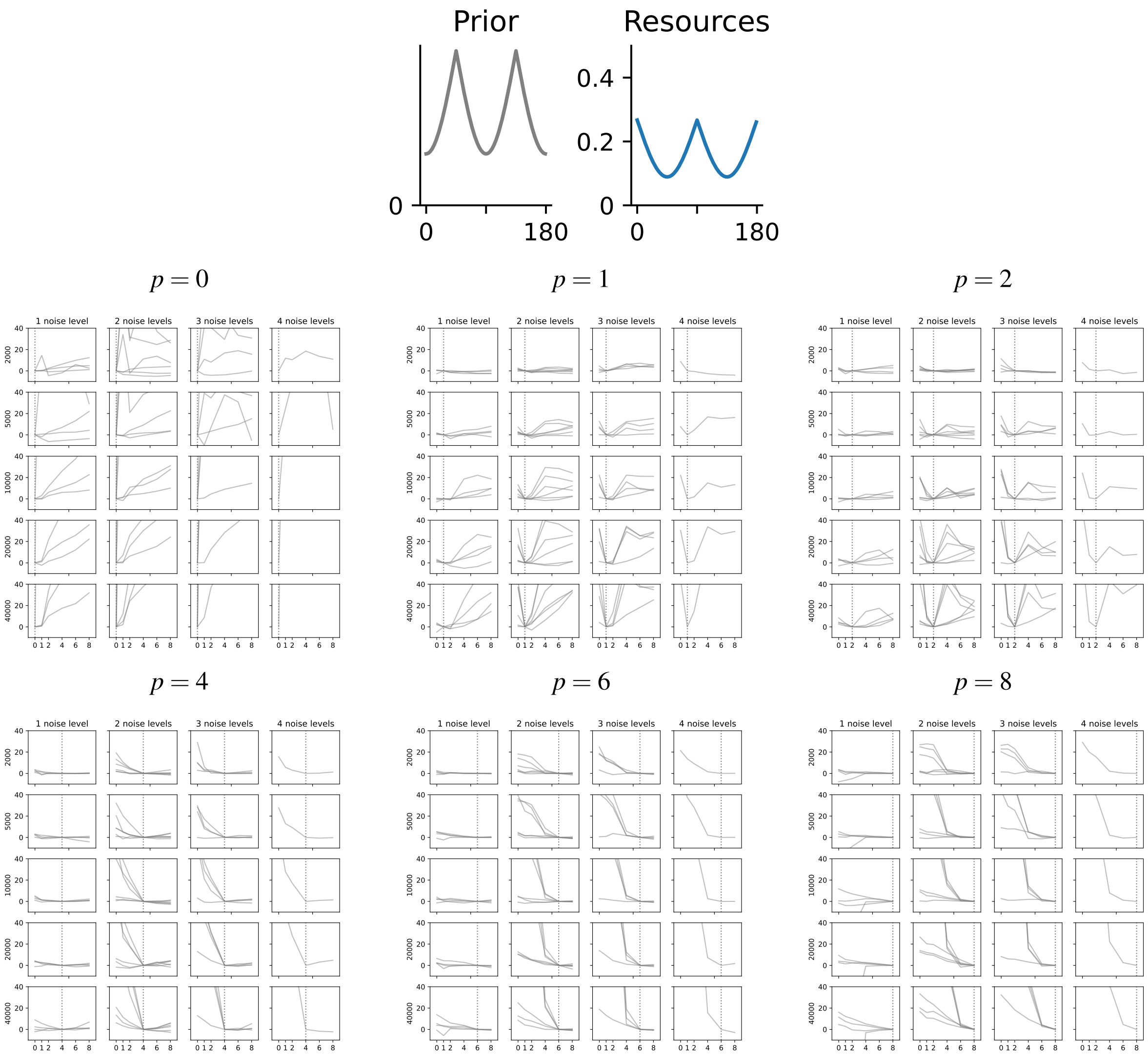

Figure S9: Circular Stimulus Space: Identifiability of the loss function depending on the number of trials (rows, from 2K to 40K), the number of of sensory noise levels (columns, from 1 to 4), and the ground truth loss function exponent (facets, $p$ from 0 to 8). Each curve represents one synthetic dataset with the given number of trials and noise levels, with the prior and encoding resources shown at the top. In each case, we include all possible combinations of the four overall noise levels, i.e., there are four cases at 1 noise level, $\binom{4}{2} = 6$ combinations at 2 noise levels, $\binom{4}{3} = 4$ combinations at 3 noise levels, and a single combination at 4 noise levels. We plot the model fit at each exponent as quantified by the difference to model fit at the ground-truth exponent, whose position is indicated with a dotted vertical line. Note that, in order to keep the y-axis range consistent across panels for easy comparison, values greater than 40 are not shown.

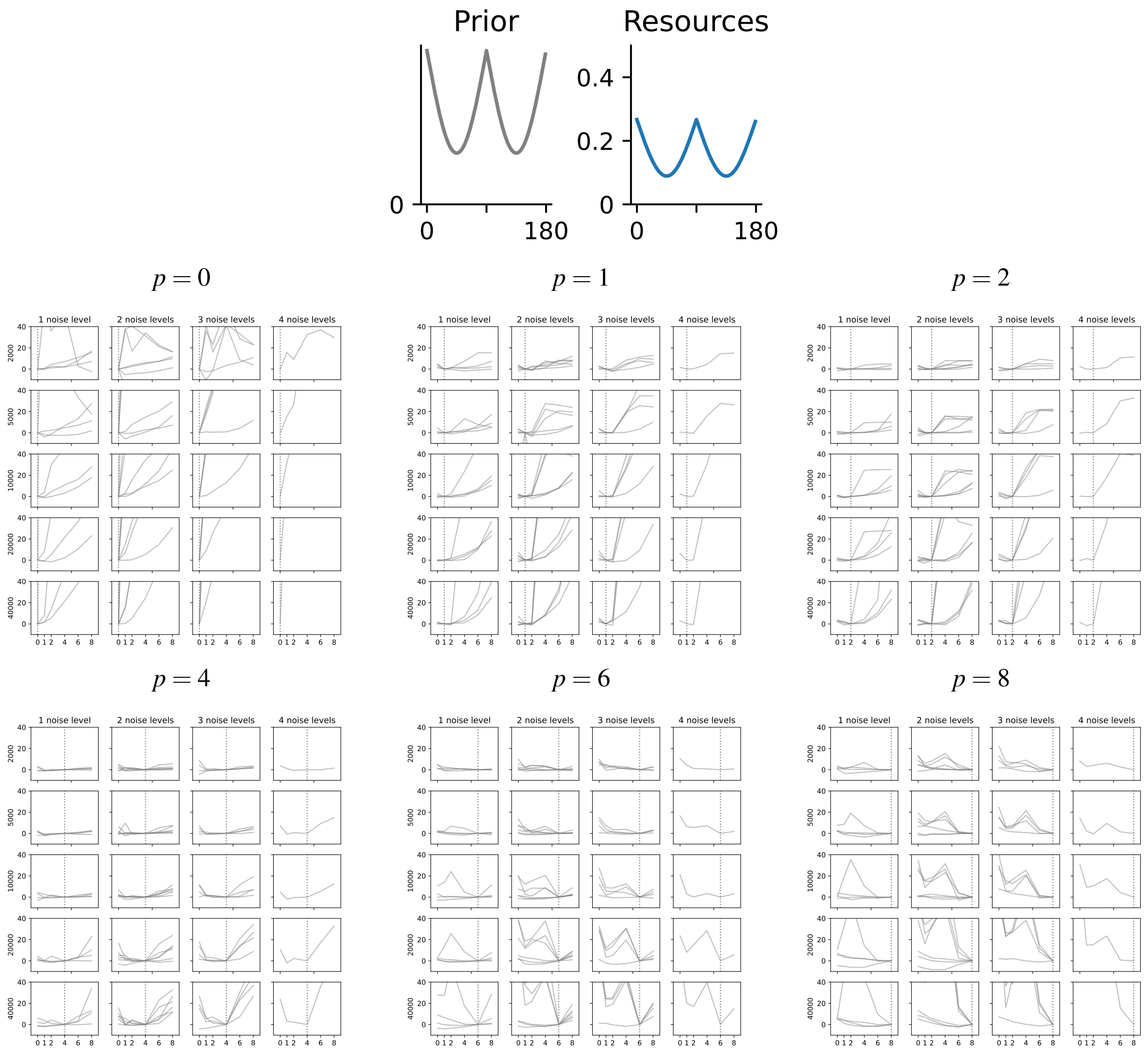

Figure S10: Circular Stimulus Space: Identifiability of the loss function depending on the number of trials (rows, from 2K to 40K), the number of of sensory noise levels (columns, from 1 to 4), and the ground truth loss function exponent (facets, $p$ from 0 to 8). Each curve represents one synthetic dataset with the given number of trials and noise levels, with the prior and encoding resources shown at the top. In each case, we include all possible combinations of the four overall noise levels, i.e., there are four cases at 1 noise level, $\binom{4}{2} = 6$ combinations at 2 noise levels, $\binom{4}{3} = 4$ combinations at 3 noise levels, and a single combination at 4 noise levels. We plot the model fit at each exponent as quantified by the difference to model fit at the ground-truth exponent, whose position is indicated with a dotted vertical line. Note that, in order to keep the y-axis range consistent across panels for easy comparison, values greater than 40 are not shown.

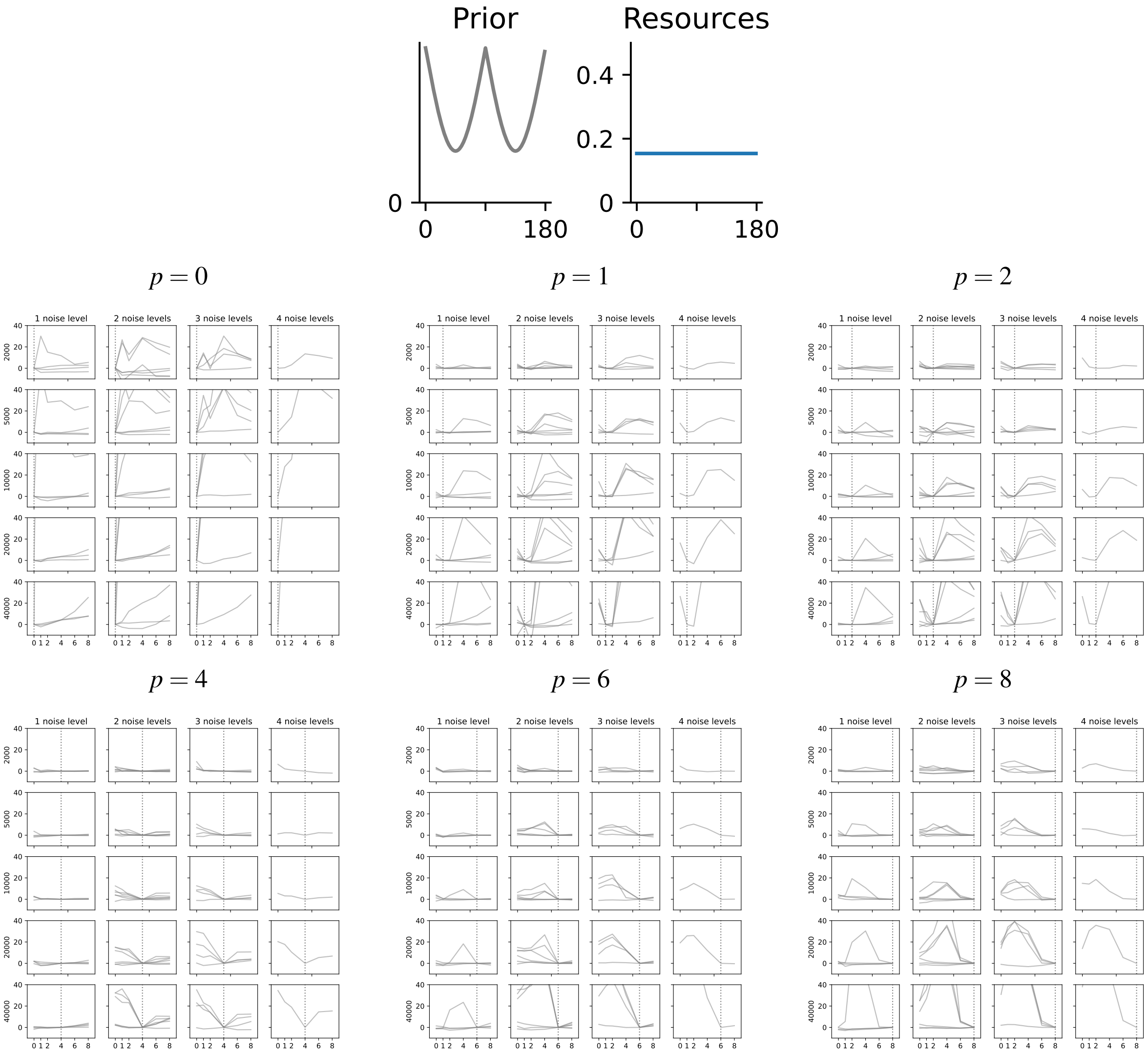

Figure S11: Circular Stimulus Space: Identifiability of the loss function depending on the number of trials (rows, from 2K to 40K), the number of of sensory noise levels (columns, from 1 to 4), and the ground truth loss function exponent (facets, $p$ from 0 to 8). Each curve represents one synthetic dataset with the given number of trials and noise levels, with the prior and encoding resources shown at the top. In each case, we include all possible combinations of the four overall noise levels, i.e., there are four cases at 1 noise level, $\binom{4}{2} = 6$ combinations at 2 noise levels, $\binom{4}{3} = 4$ combinations at 3 noise levels, and a single combination at 4 noise levels. We plot the model fit at each exponent as quantified by the difference to model fit at the ground-truth exponent, whose position is indicated with a dotted vertical line. Note that, in order to keep the y-axis range consistent across panels for easy comparison, values greater than 40 are not shown.

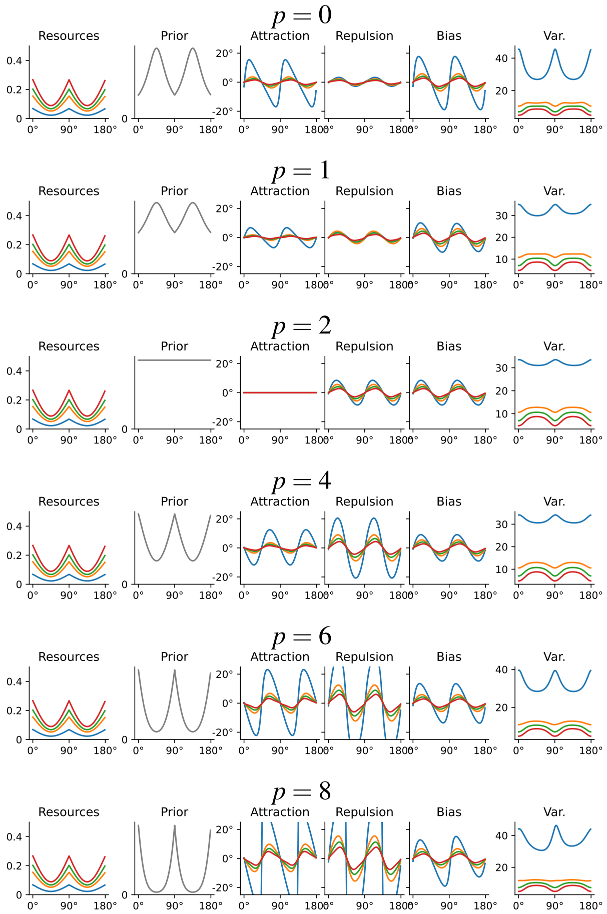

Figure S12: Models that have the same response distribution in the limit of zero noise (same encoding; prior transformed by Eq. 3), but different behavior for nonzero noise. For such models, the loss function may be poorly identified at one level of sensory noise, but well identified when more levels of sensory noise are available (see Figure S8). Theorems 3–4 show that this situation is typical for most combinations of prior and encoding.

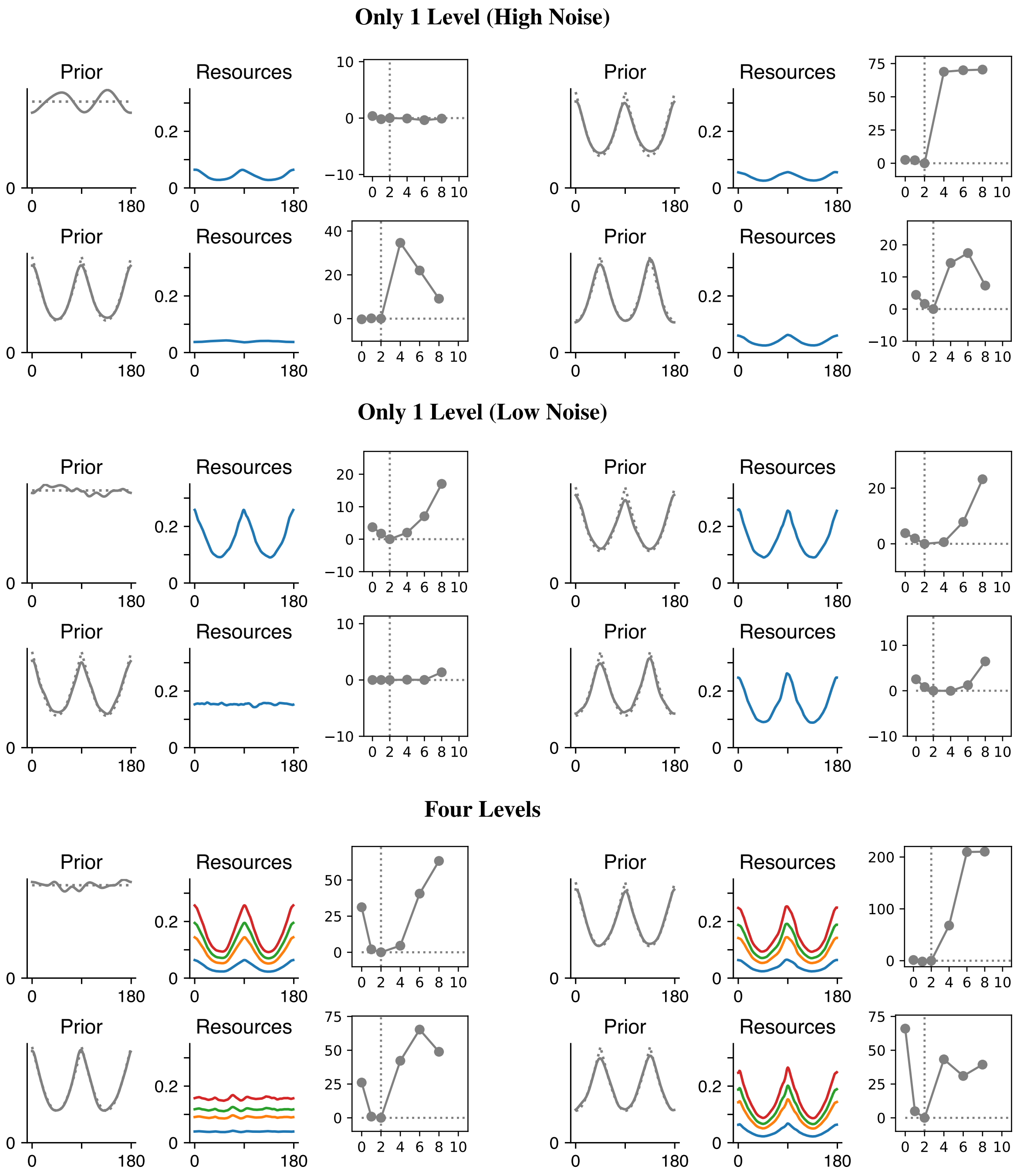

Figure S13: Fits at one or four levels of sensory noise, for four combinations of prior (left) and encoding (middle), with ground truth exponent $p = 2$. We show both the prior and encoding recovered when fitting at the ground truth exponent ($p = 2$), and NLL over exponents (right). Results at 40K trials. At 1 level of sensory noise, NLL does not reliably single out the correct loss function. High noise can lead to a large $\Delta NLL$, but does not reliably distinguish high from low exponents. The shape of the prior is also recovered incorrectly in some cases even when presupposing the correct $p$. Low noise leads to more reliable recovery, but the statistical signal in $\Delta NLL$ can be weak. At four levels of sensory noise, both the prior and the loss function are recovered reliably.

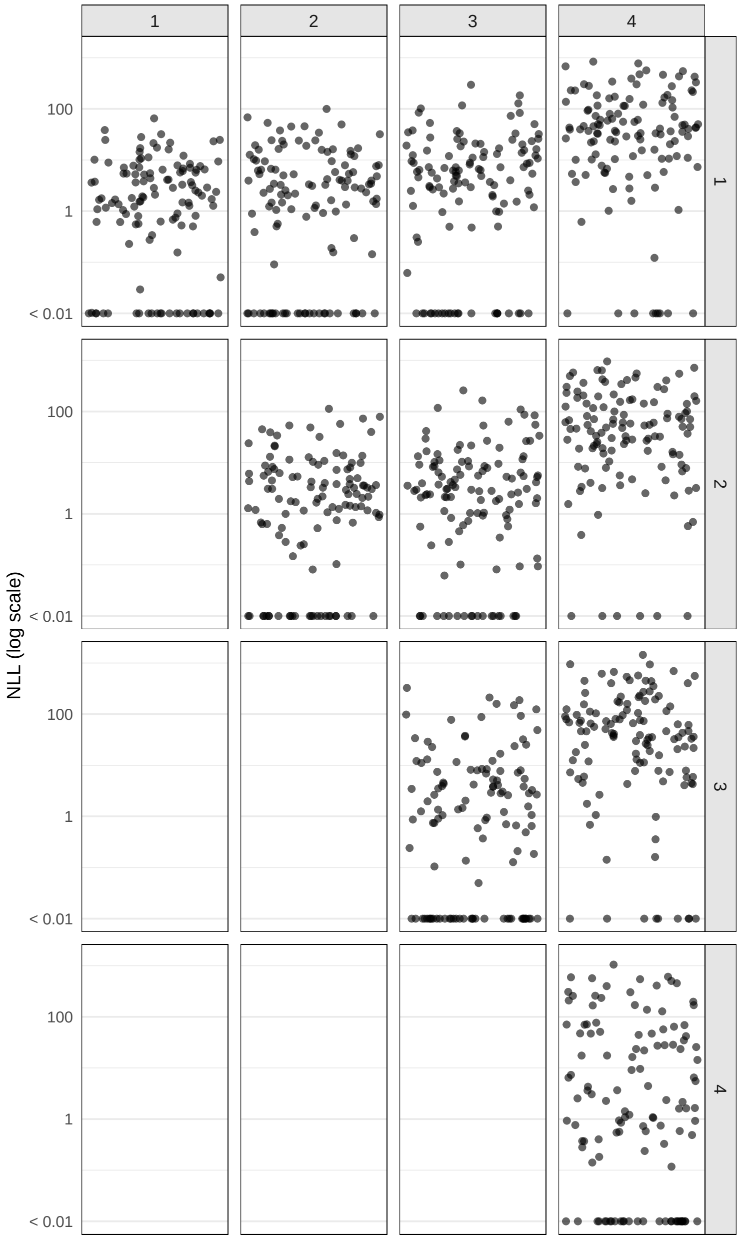

Figure S14: Subset of the datapoints of Main Paper Figure 5c at N=40,000 trials and one or two noise levels, broken down by the two noise levels (1=smallest noise, 4=largest noise). Panels on the diagonal correspond to settings with only one noise level. As we also found in Figure S13, higher noise levels tend to lead to higher $\Delta$ NLL, though, on their own, they often are not sufficient for singling out the ground-truth loss (Figure S13). Here, we find that across noise magnitudes, combining with a different noise level boosts $\Delta$ NLL (and thus identifiability), and mainly so when combining with a substantially different level (e.g., combining level 1 with level 4). Intuitively, more distinct noise levels may be more likely to provide complementary evidence about the model; Theorem 3 formalizes this by requiring that the ratio of noise magnitudes be bounded away from 1.

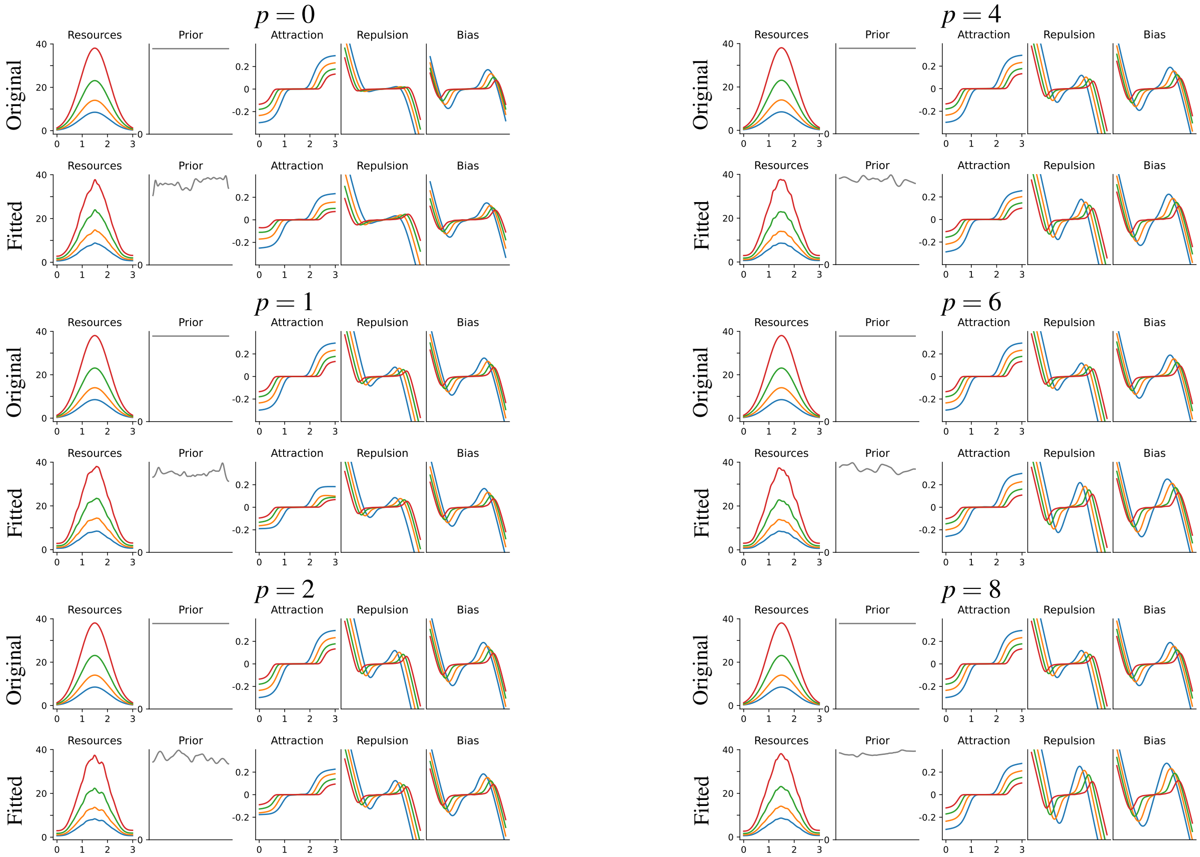

Figure S15: Interval Stimulus Space: Identifiability of encoding and prior (40K trials, 4 levels of sensory noise). We note that, in the vicinity of the boundary, the biases have an additional component, added to the attraction and repulsion terms in Equation 2 of the main paper, described formally in Theorem 3 of Hahn and Wei [5]. See Section S4.1 for discussion of Attraction and Repulsion components.

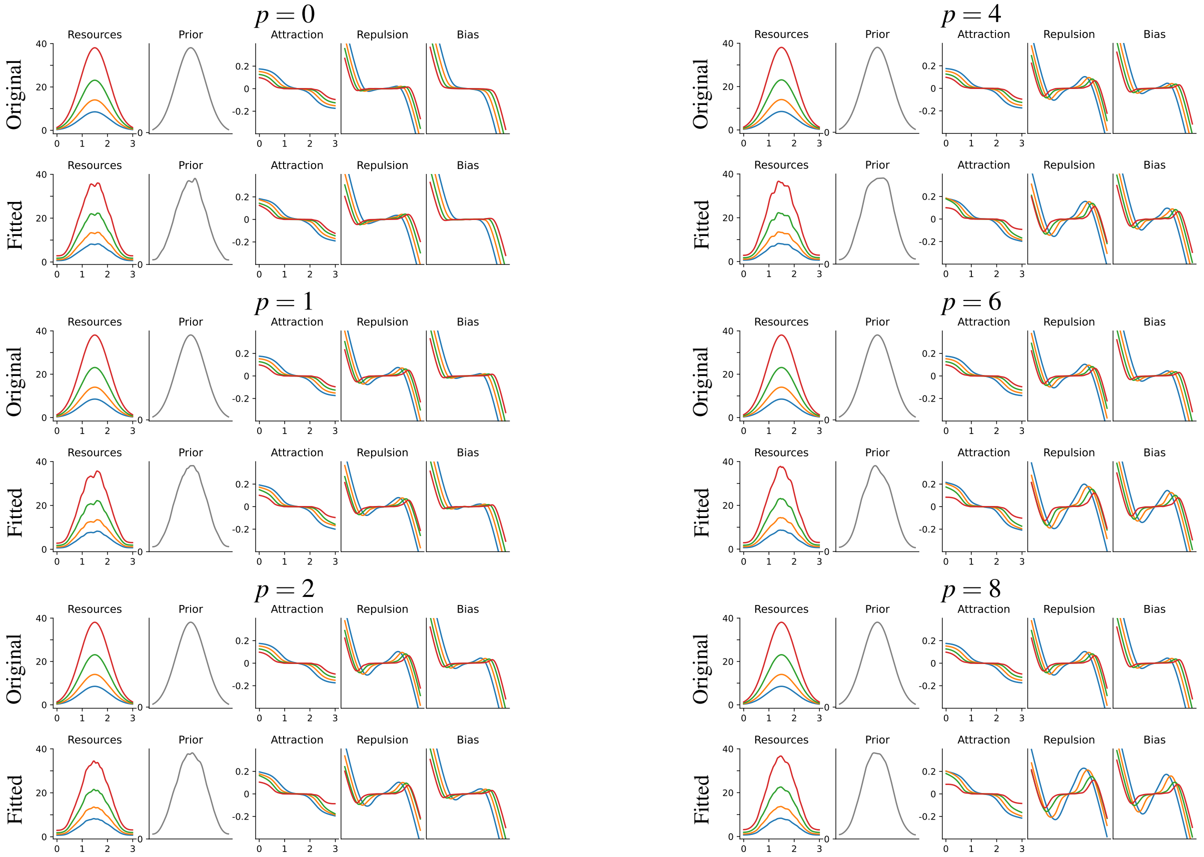

Figure S16: Interval Stimulus Space: Identifiability of encoding and prior (40K trials, 4 levels of sensory noise). While we here assume in fitting that the loss function is already known, results in Figure S8 show that the loss function is also largely identifiable in this situation. We note that, in the vicinity of the boundary, the biases have an additional component, added to the attraction and repulsion terms in Equation 2 of the main paper, described formally in Theorem 3 of Hahn and Wei [5]. We note that the model of Polanía et al. [17] is conceptually similar to this situation; a repulsive bias in the center and an attractive bias closer to the boundaries is observed there. See Section S4.1 for discussion of Attraction and Repulsion components.

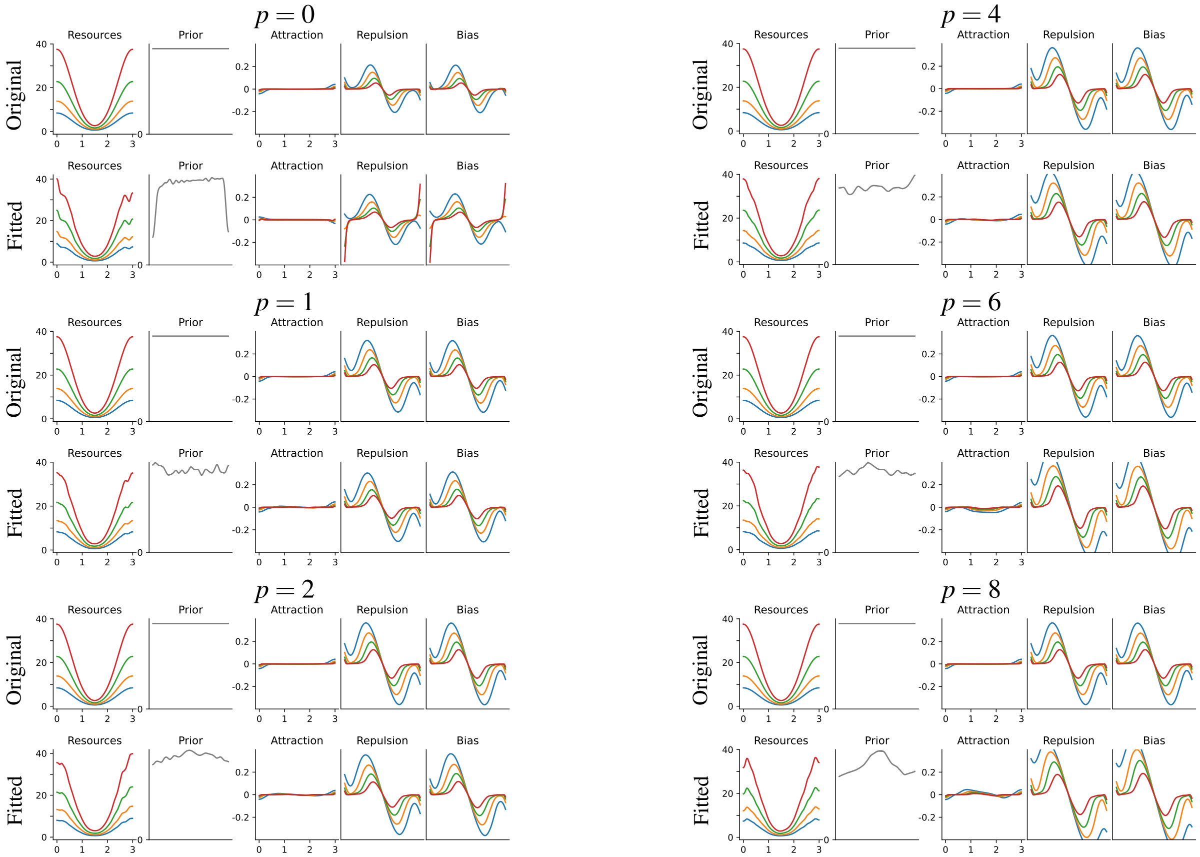

Figure S17: Interval Stimulus Space: Identifiability of encoding and prior (40K trials, 4 levels of sensory noise). We note that, in the vicinity of the boundary, the biases have an additional component, added to the attraction and repulsion terms in Equation 2 of the main paper, described formally in Theorem 3 of Hahn and Wei [5]. See Section S4.1 for discussion of Attraction and Repulsion components.

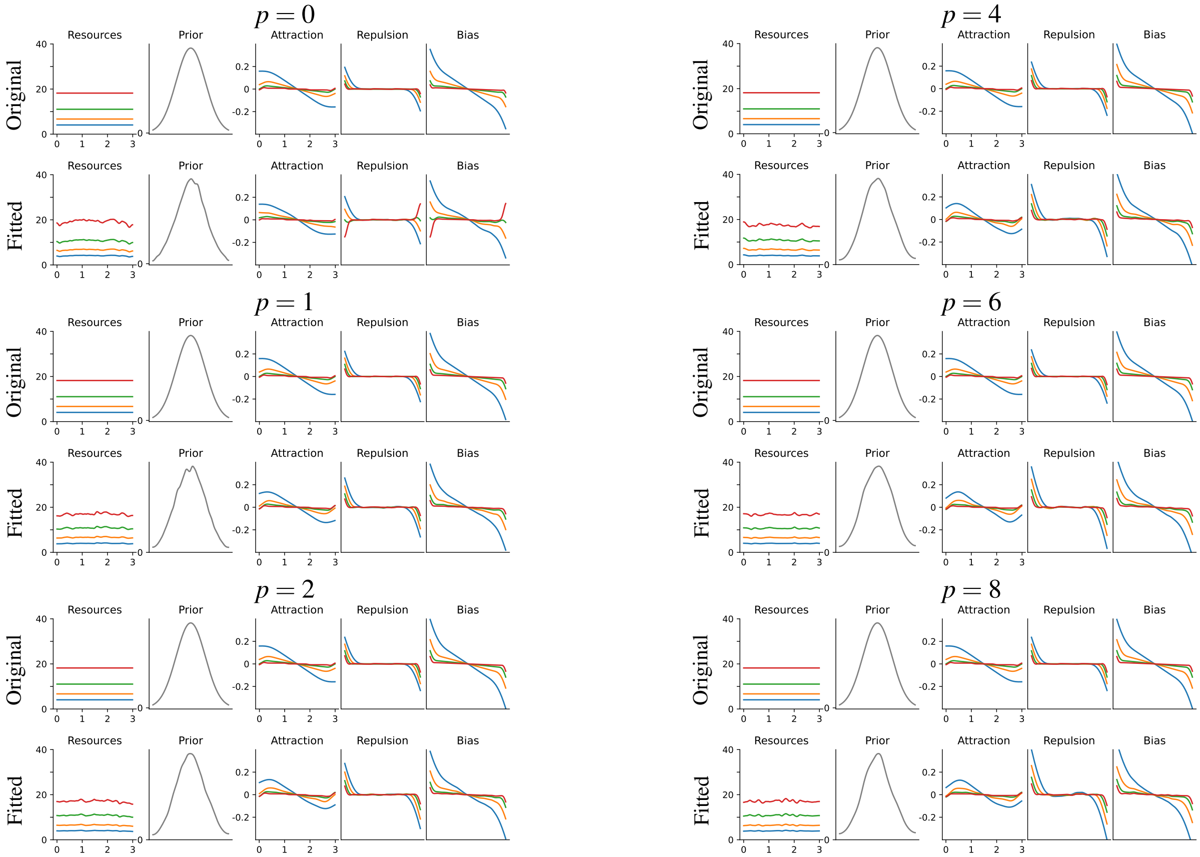

Figure S18: Interval Stimulus Space: Identifiability of encoding and prior (40K trials, 4 levels of sensory noise). We note that, in the vicinity of the boundary, the biases have an additional component, added to the attraction and repulsion terms in Equation 2 of the main paper, described formally in Theorem 3 of Hahn and Wei [5]. See Section S4.1 for discussion of Attraction and Repulsion components.

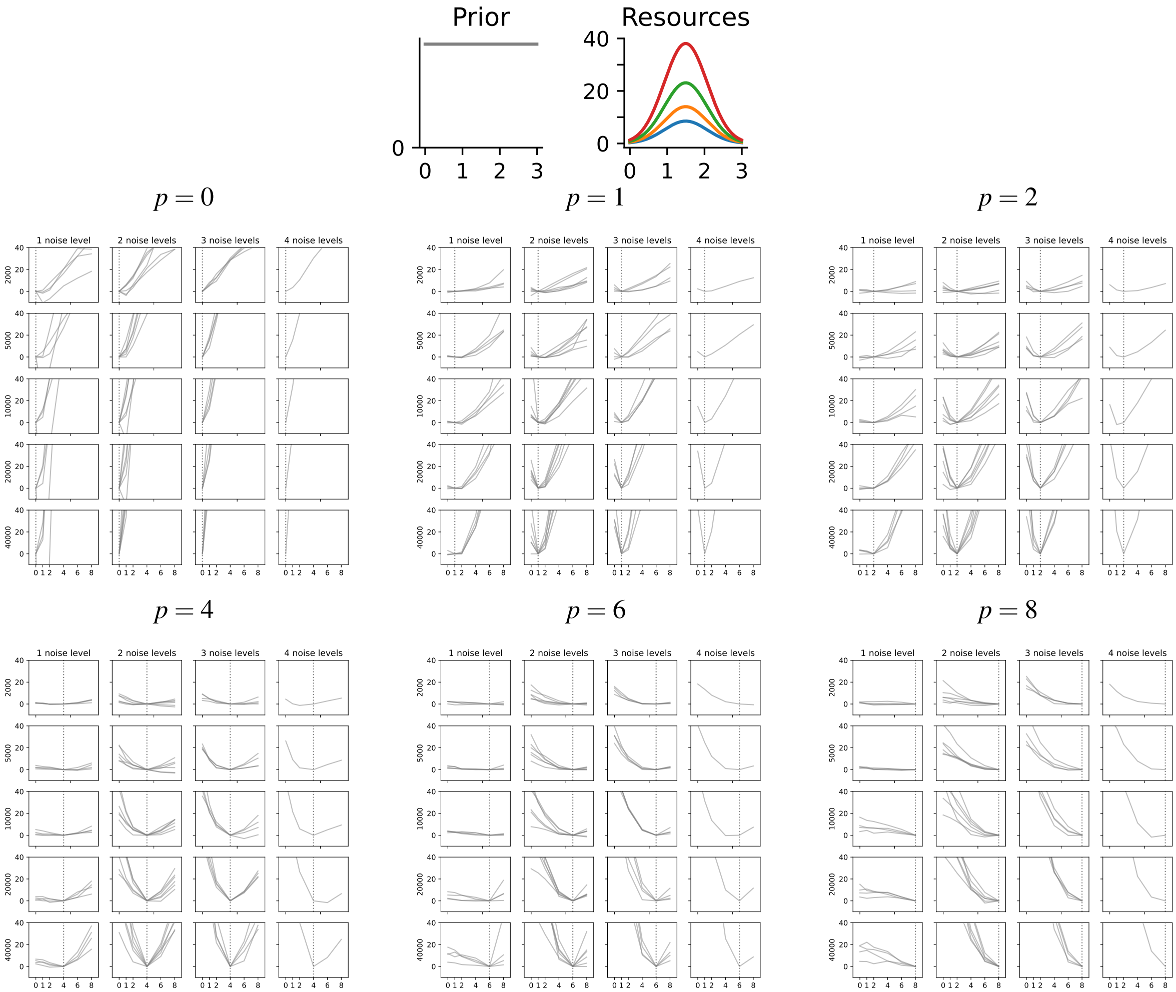

Figure S19: Interval Stimulus Space: Identifiability of the loss function depending on the number of trials and of sensory noise levels.

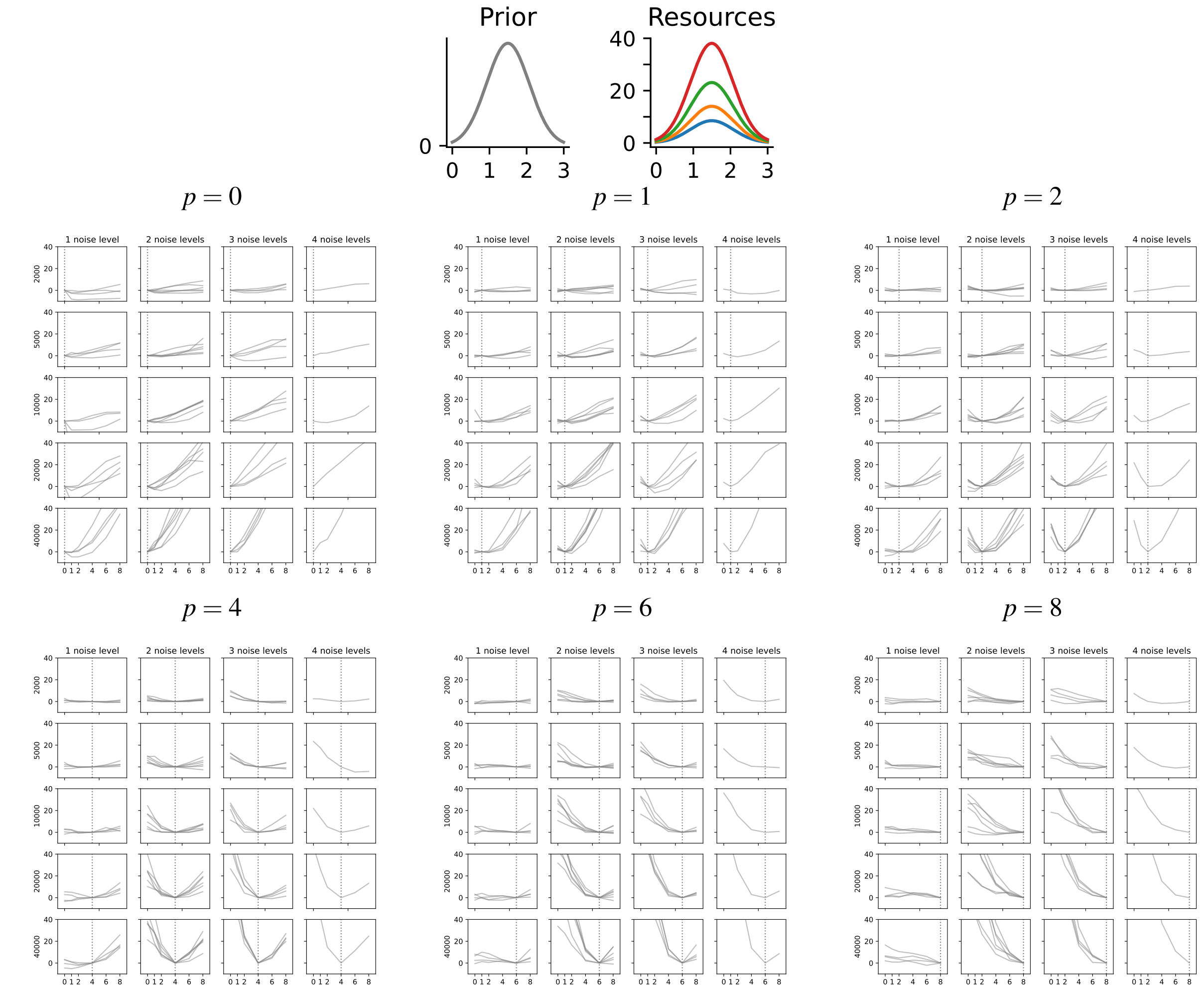

Figure S20: Interval Stimulus Space: Identifiability of the loss function depending on the number of trials and of sensory noise levels.

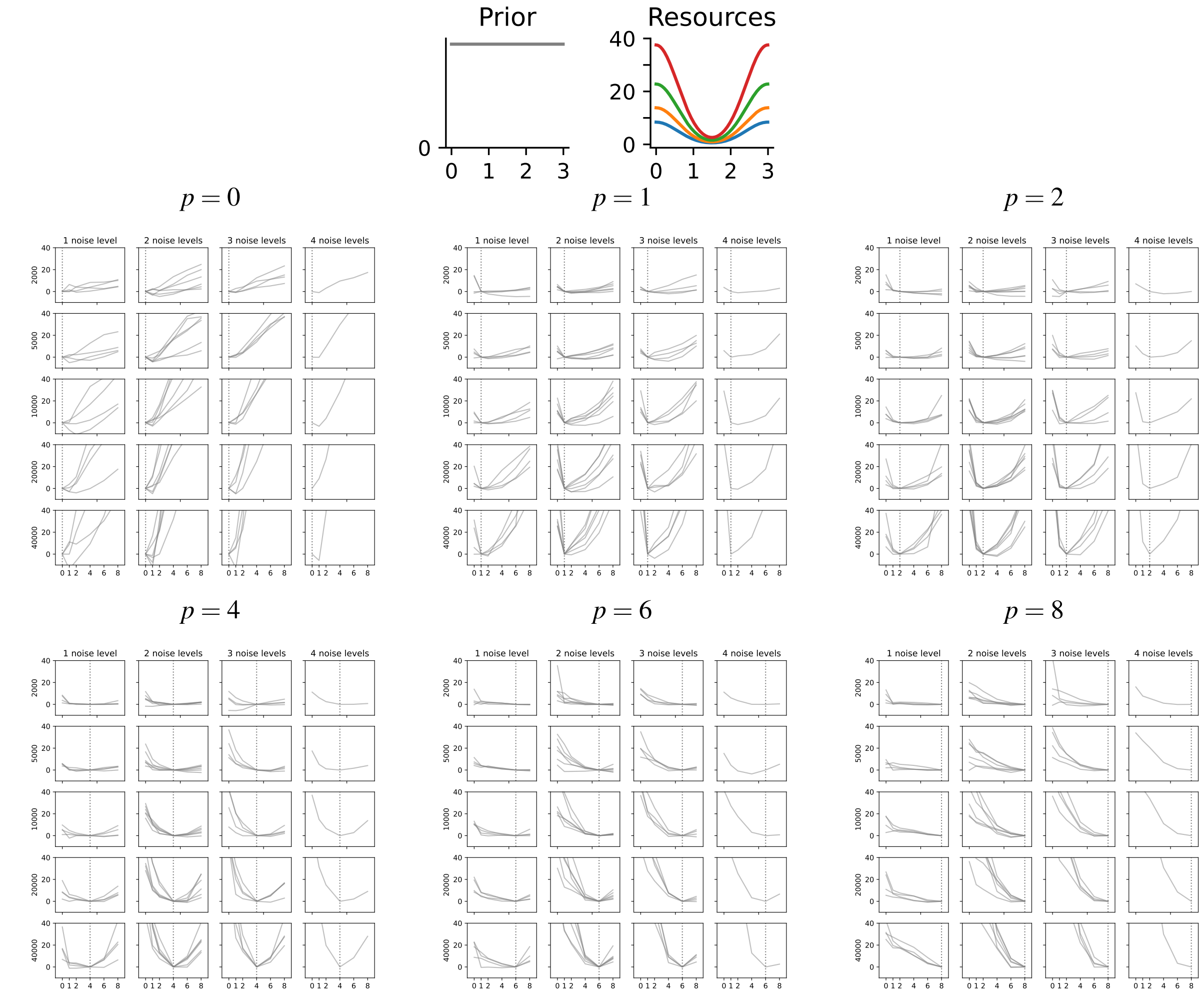

Figure S21: Interval Stimulus Space: Identifiability of the loss function depending on the number of trials and of sensory noise levels.

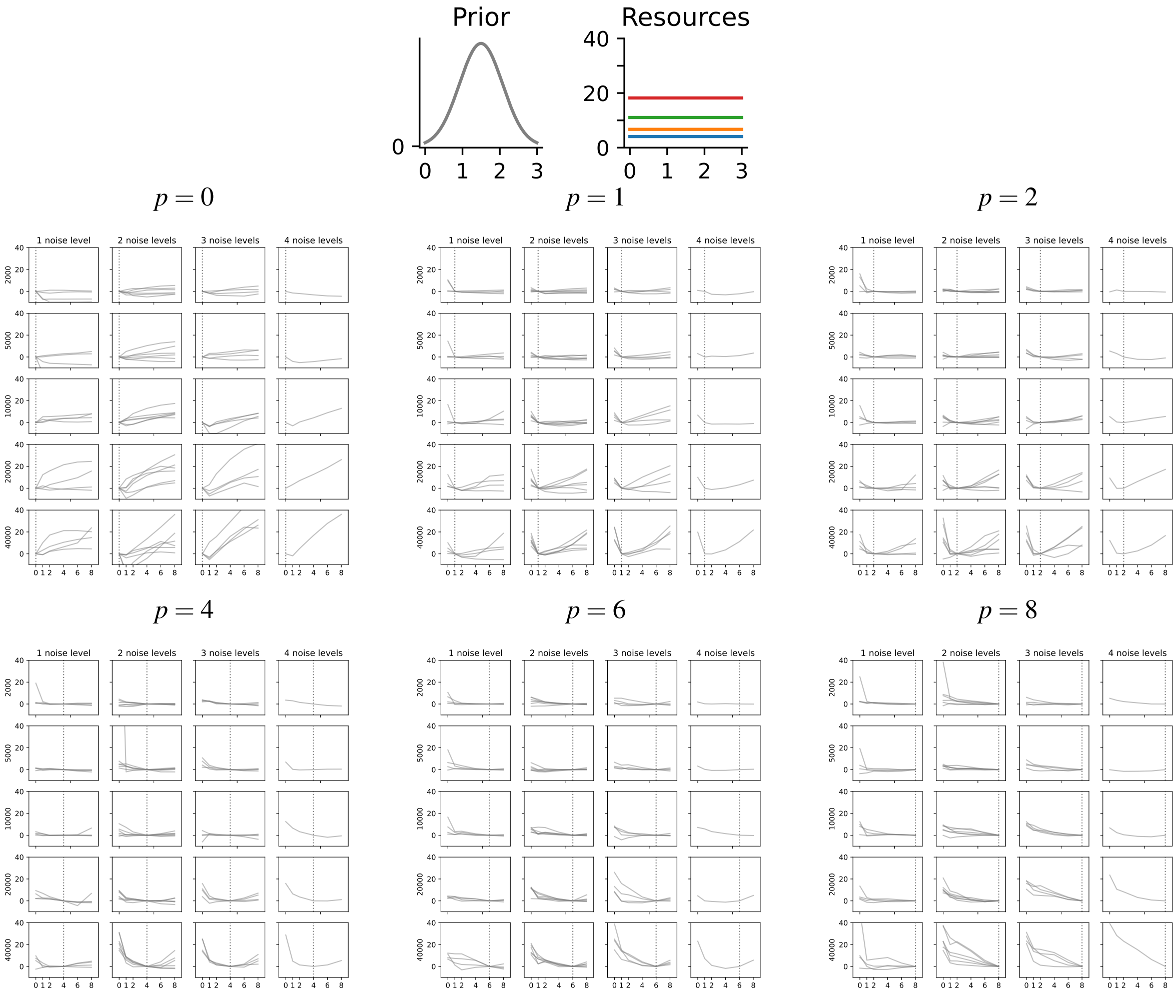

Figure S22: Interval Stimulus Space: Identifiability of the loss function depending on the number of trials and of sensory noise levels.

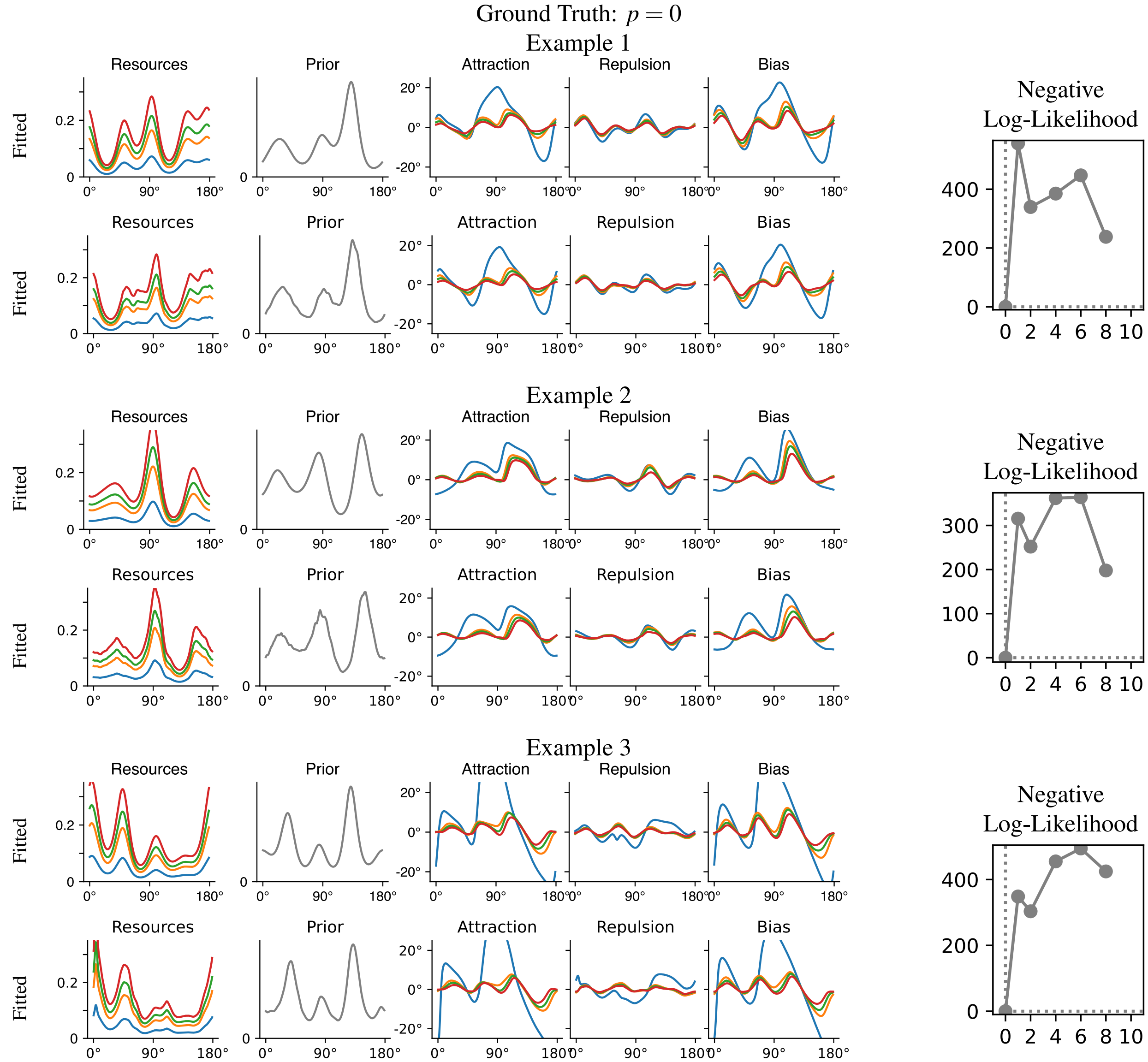

Figure S23: Supplement to Figure 4: Randomly constructed models. We simulate three models at the ground truth loss function exponent indicated. The loss function is clearly identified by model fit, and, when fitting at this loss, prior and encoding are recovered. See Section S4.1 for discussion of Attraction and Repulsion components.

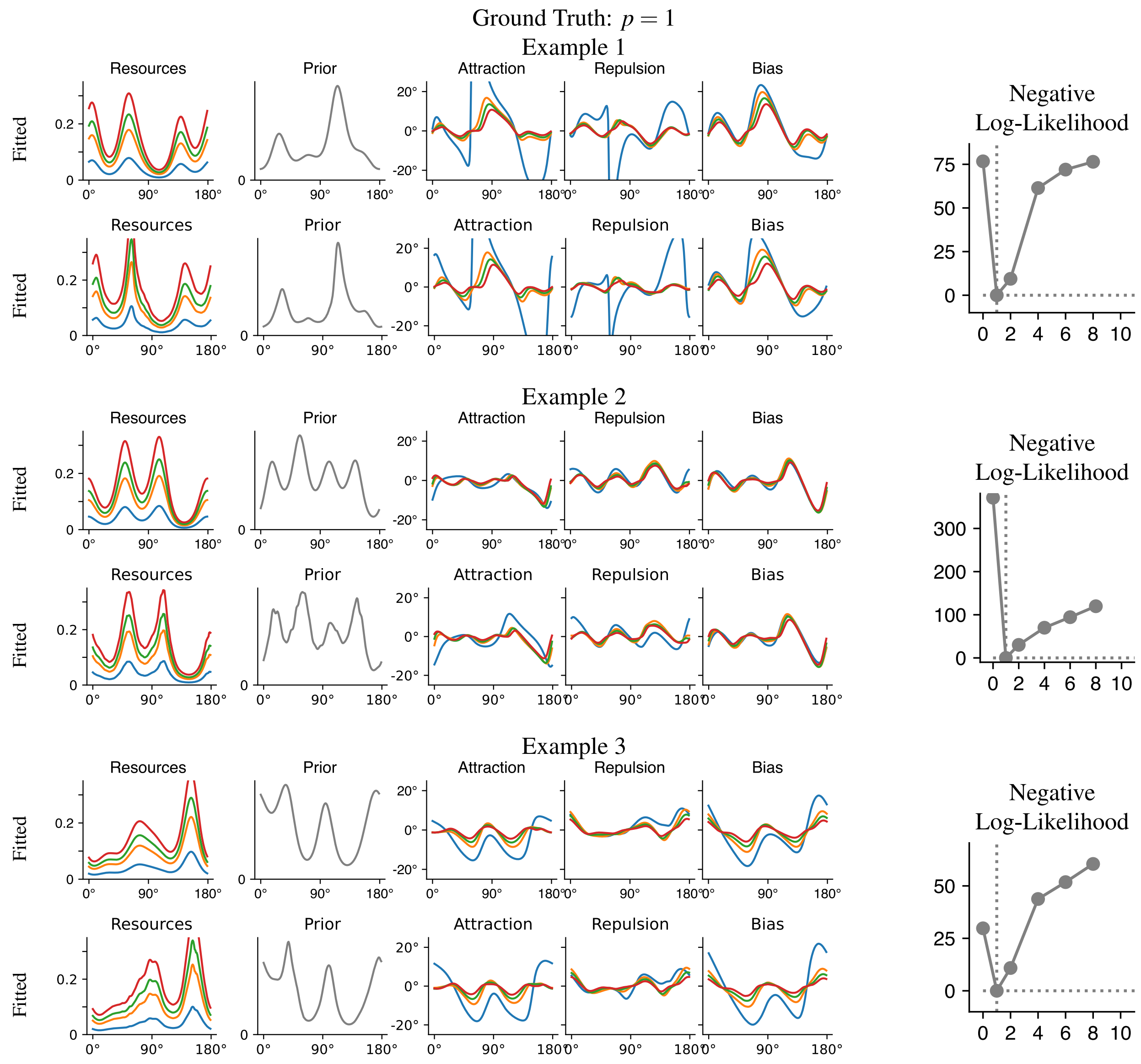

Figure S24: Supplement to Figure 4: Randomly constructed models. We simulate three models at the ground truth loss function exponent indicated. The loss function is clearly identified by model fit, and, when fitting at this loss, prior and encoding are recovered. See Section S4.1 for discussion of Attraction and Repulsion components.

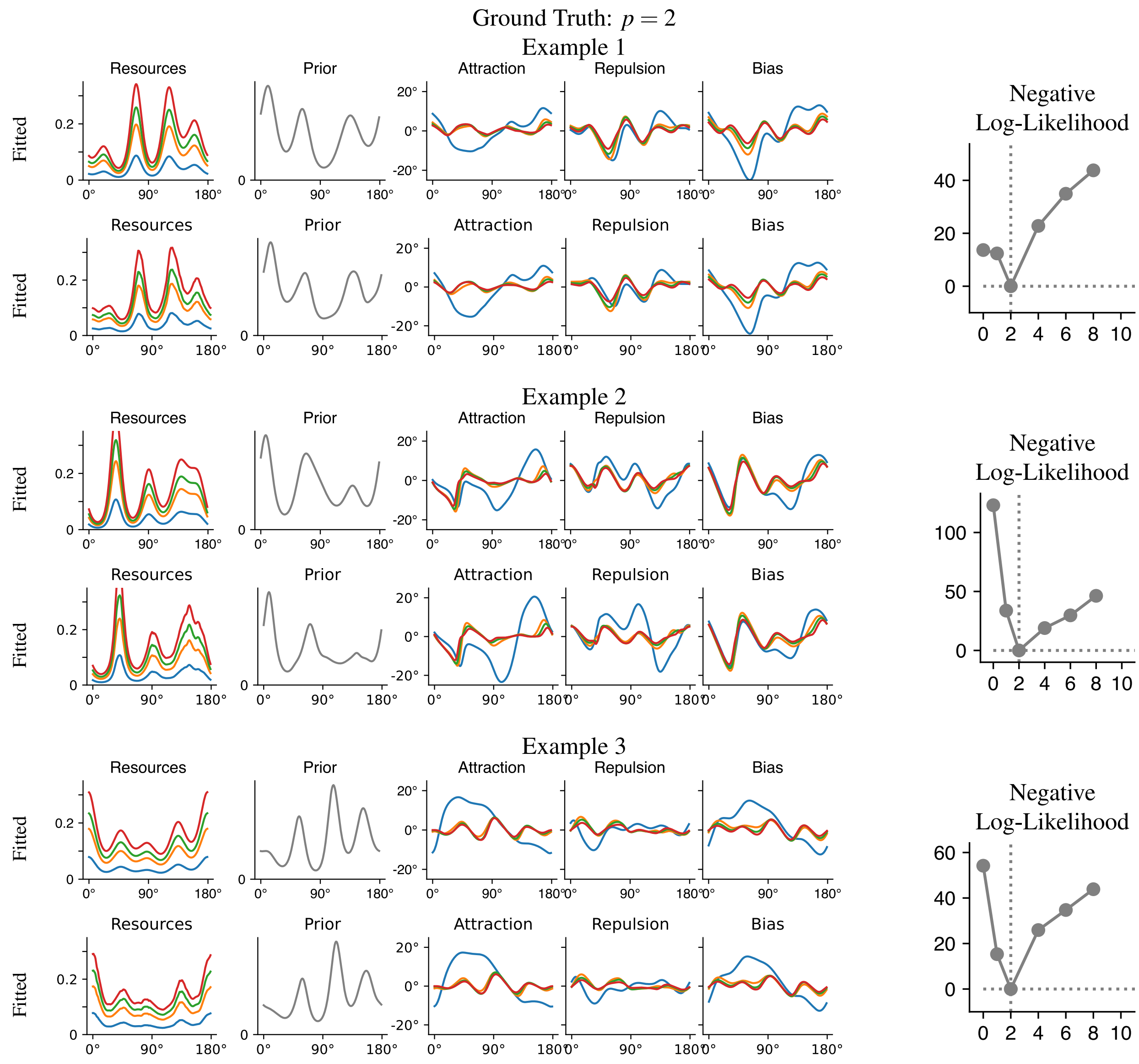

Figure S25: Supplement to Figure 4: Randomly constructed models. We simulate three models at the ground truth loss function exponent indicated. The loss function is clearly identified by model fit, and, when fitting at this loss, prior and encoding are recovered. See Section S4.1 for discussion of Attraction and Repulsion components.

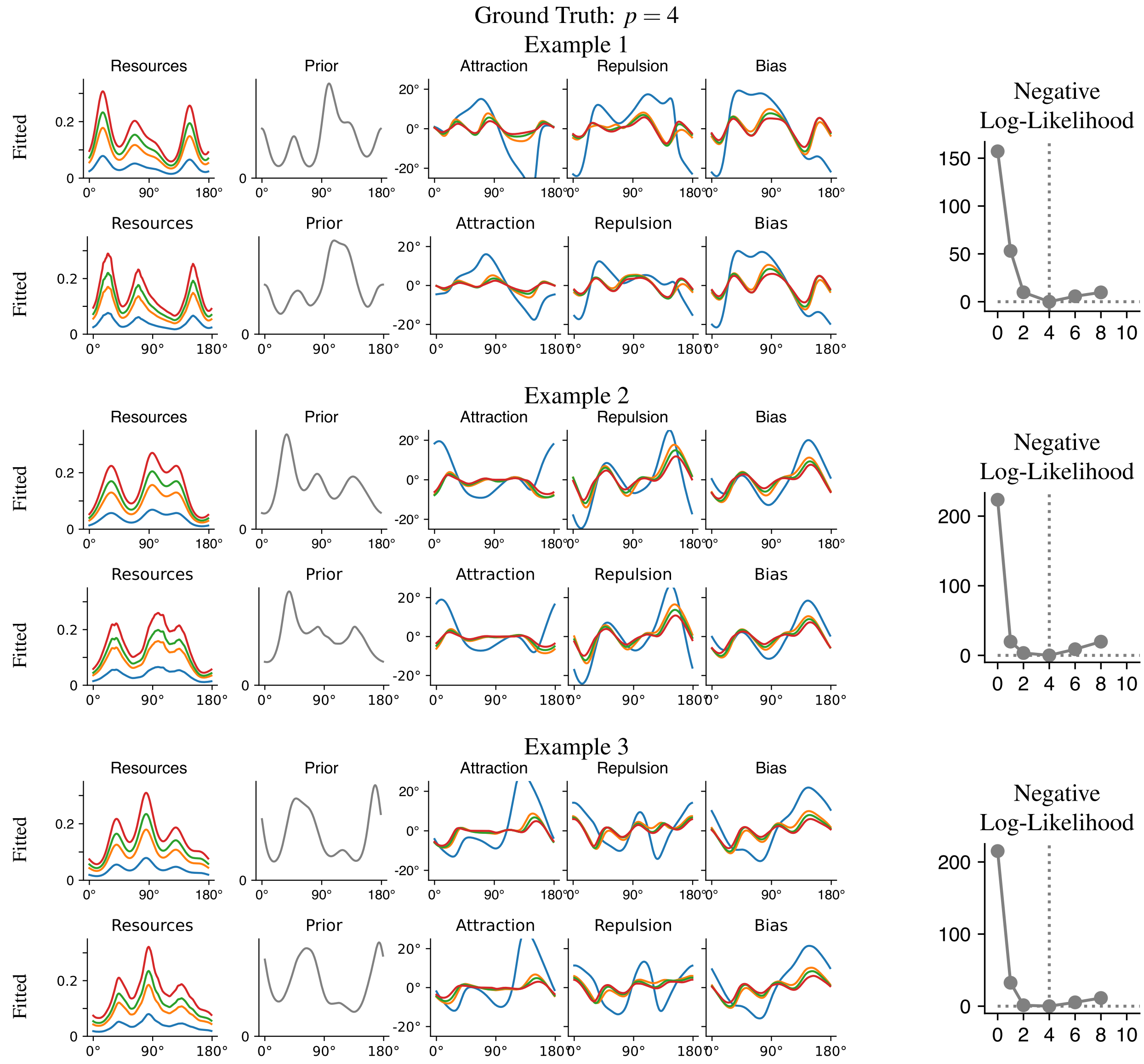

Figure S26: Supplement to Figure 4: Randomly constructed models. We simulate three models at the ground truth loss function exponent indicated. The loss function is clearly identified by model fit, and, when fitting at this loss, prior and encoding are recovered. See Section S4.1 for discussion of Attraction and Repulsion components.

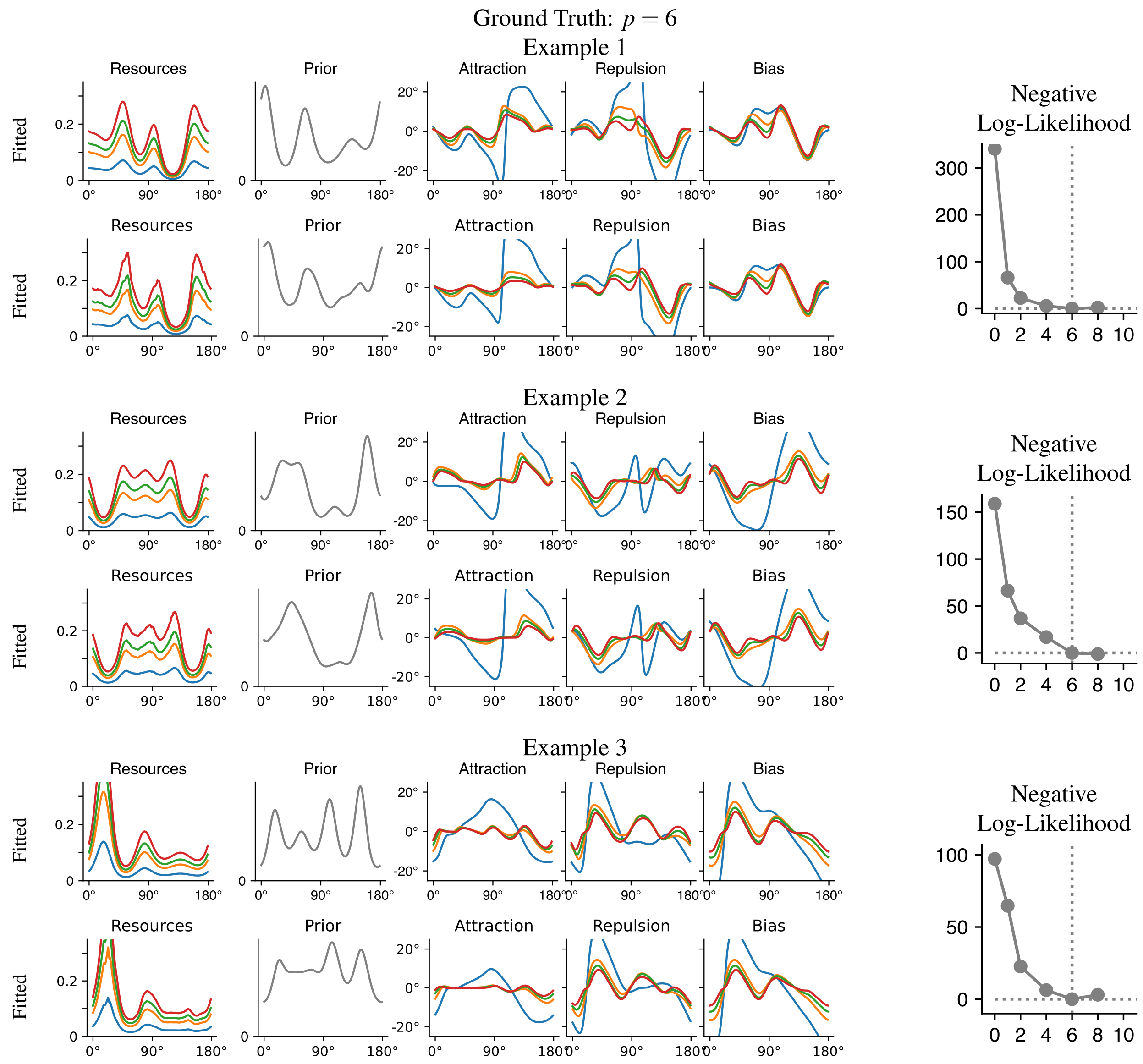

Figure S27: Supplement to Figure 4: Randomly constructed models. We simulate three models at the ground truth loss function exponent indicated. The loss function is clearly identified by model fit, and, when fitting at this loss, prior and encoding are recovered. See Section S4.1 for discussion of Attraction and Repulsion components.

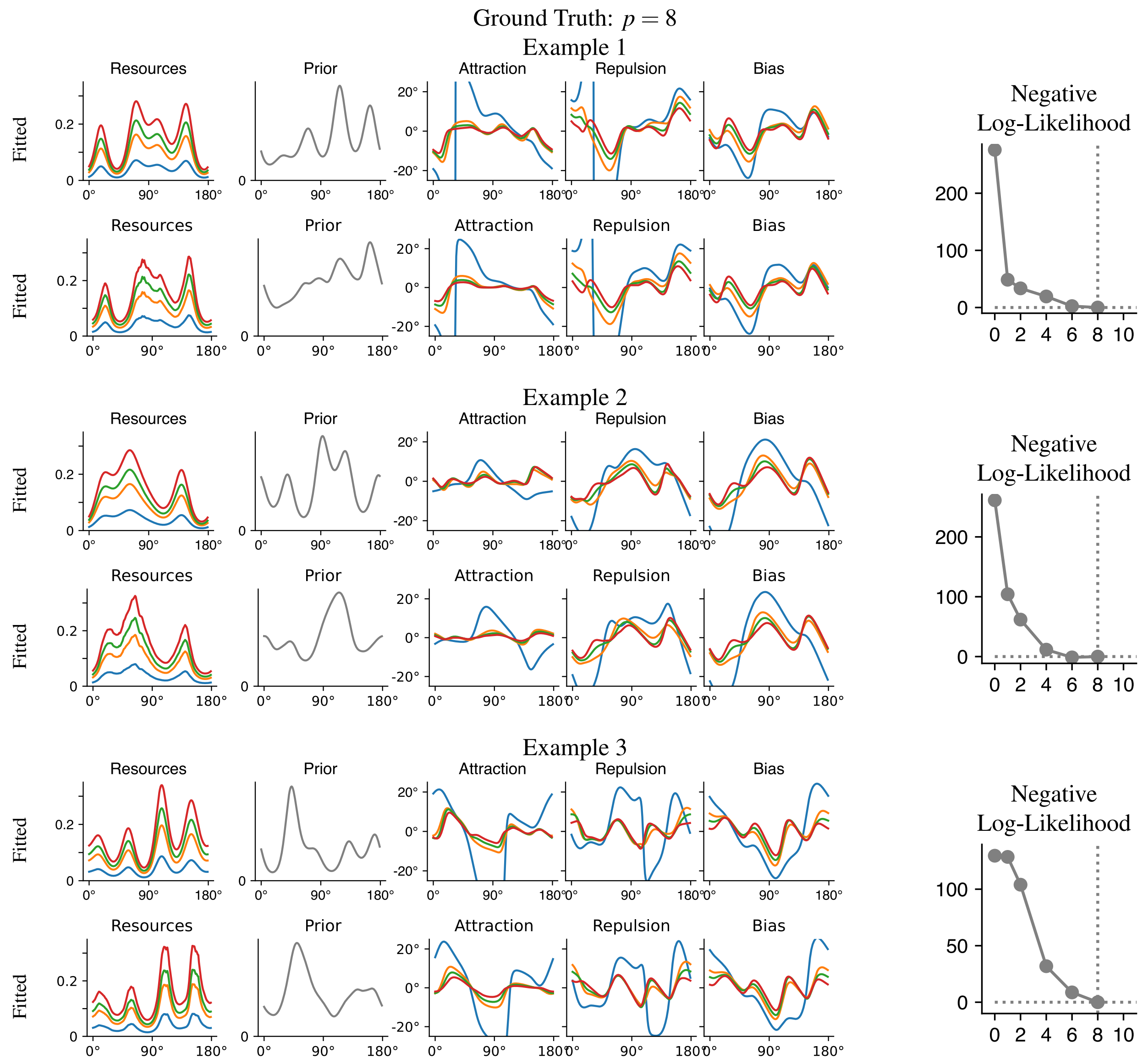

Figure S28: Supplement to Figure 4: Randomly constructed models. We simulate three models at the ground truth loss function exponent indicated. The loss function is clearly identified by model fit, and, when fitting at this loss, prior and encoding are recovered. See Section S4.1 for discussion of Attraction and Repulsion components.

### S5 Identifiability in Other Situations

#### S5.1 Identifiability under Two-Alternative Forced Choice Task (2AFC)

##### S5.1.1 Theoretical Guarantee

Here, we show how our identifiability guarantees apply for two-alternative forced-choice (2AFC) tasks. In such experiments, subjects are typically presented with two stimuli in the same trial, and asked to judge which one is larger.

We assume the following generative model of a single trial in the 2AFC task:

1. Encode $\theta_1, \theta_2$ independently, according to the sensory noise levels $\sigma_1, \sigma_2$ at which each of them is presented: $m_1 = F(\theta_1) + \sigma_1 \varepsilon_1$, $m_2 = F(\theta_2) + \sigma_2 \varepsilon_2$ where $\varepsilon_1, \varepsilon_2 \sim N(0,1)$ independent.
2. Decode $\widehat{\theta_1}$ from $m_1$, $\widehat{\theta_2}$ from $m_2$.
3. Respond based on which of these is larger. There may be a nonzero lapse rate, determining

This is essentially the model assumed by Girshick et al. [4], Stocker and Simoncelli [21].

We first point out that, except for the role of motor noise, forced choice provides no information beyond that contained in response distributions. This because one can simulate a 2AFC task by encoding and decoding the two stimuli separately and compare the response distributions. We now investigate to what extent 2AFC response distributions might provide *the same* information as response distributions, and allow recoverability. 2AFC response distributions where both stimuli are subjected to the same sensory noise recover exactly the encoding, but provide no information about prior and loss function, due to (25). We now investigate the more general setting of 2AFC Response distributions when the two stimuli are subjected to different amounts of noise. These can be expected to provide information about the *relative bias* between two conditions. Intuitively, comparing this across multiple conditions should enable recovering the bias. Formally, we obtain a guarantee analogous to Theorem 2, though three levels of sensory noise are required:

**Theorem S31.** *There is a function $\Phi$ mapping 2AFC response distributions to $\tilde{M} \in \mathfrak{M}$ such that the following holds. Assume $0 < \sigma_1^2 < \sigma_2^2 < \sigma_3^2$ are three sensory noise variances. Assume the response distribution is derived at noise levels $\sigma_1^2, \sigma_2^2, \sigma_3^2$ and lapse rate $\lambda > 0$ from the ground-truth model $M = \langle F', p_{prior}, p \rangle \in \mathfrak{M}$. Then for the model*

$$\tilde{M} = \langle \tilde{F}', p_{\tilde{prior}}, \tilde{q} \rangle \in \mathfrak{M} \tag{108}$$

*output by $\Phi$, we have – provided $M \notin \Omega$ – the following for all $\theta$*

$$\widehat{F}(\theta) = F(\theta) + O(\sigma_1^2) \qquad p_{\tilde{prior}}(\theta) = p_{prior}(\theta) + O(\sigma_1^2) \qquad \tilde{q} = p \text{ for } \sigma_1^2 \text{ small}$$

*in the limit where $\sigma_1, \sigma_2, \sigma_3 \to 0$, provided:*

$$0 < C_1 < \frac{\sigma_1}{\sigma_2}, \frac{\sigma_2}{\sigma_3} < C_2 < 1 < \infty$$

*for some $C_1, C_2, C_3, C_4$, and $\sigma_2 - \sigma_1 \neq \sigma_3 - \sigma_2$. Constants in the $O(\cdot)$ expressions depend on $C_1, C_2, C_3, C_4$, and on the regularity of $F$ and $p_{prior}$ and their derivatives, but not on $\theta$.*

*Proof.* As in the proof of Theorem 2, we introduce a parameter $t$ such that $\sigma_1, \sigma_2, \sigma_3$ are constant multiples of $t$ as $t \to 0$. First, we note that the lapse rate can be identified, when $t$ is small, from the response distribution when $\theta_1$ and $\theta_2$ are very far away. Second, $F$ and $\sigma_1, \sigma_2, \sigma_3$ can already be inferred from response distributions under equal sensory noise by Theorem S10. We now specifically consider a pair of sensory noise magnitudes, say, $\sigma_1, \sigma_2$. Assume that $\theta_1$ and $\theta_2$ are perceived at sensory noise magnitudes $\sigma_1, \sigma_2$, respectively. Then, for the respective decoded estimates $\widehat{\theta_1}$, $\widehat{\theta_2}$:

$$\begin{aligned}\mathbb{P}(\widehat{\theta_1} < \widehat{\theta_2}) =& \mathbb{P}\left(f_1(F(\theta_1) + \sigma_1\varepsilon_1) < f_2(F(\theta_2) + \sigma_2\varepsilon_2)\right) \\ =& \mathbb{P}\left(F(\theta_1) + \sigma_1\varepsilon_1 < f_1^{-1}(f_2(F(\theta_2) + \sigma_2\varepsilon_2))\right)\end{aligned}$$

where $f_1, f_2$ are the two decoding functions (Definition S11) mapping $F^{-1}(m)$ to the estimate $\widehat{\theta}$, under noise magnitudes $\sigma_1, \sigma_2$, respectively. This permits recovering the function

$$f_1^{-1} \circ f_2 : \theta \mapsto f_1^{-1}(f_2(\theta))$$

In the low-noise regime, writing $\sigma_1 = at$ and $\sigma_2 = bt$, we can write

$$f_1(\theta) = \theta + at\,\mathcal{C}_{dec}(\theta) + a^2t^2\mathcal{D}_{dec}(\theta) + O(t^3)$$
$$f_2(\theta) = \theta + bt\,\mathcal{C}_{dec}(\theta) + b^2t^2\mathcal{D}_{dec}(\theta) + O(t^3)$$

where $\mathcal{C}_{dec}$ and $\mathcal{D}_{dec}$ as per Eq. 55. We now aim to show the following:

$$f_1^{-1}(f_2(\theta)) = \theta + (b-a)t\,\mathcal{C}_{dec}(\theta) + t^2\Big[(b^2-a^2)\mathcal{D}_{dec}(\theta) - a(b-a)\mathcal{C}_{dec}(\theta)\mathcal{C}'_{dec}(\theta)\Big] + O(t^3). \tag{109}$$

Now, comparing the expression for $f_1^{-1}(f_2(\theta))$ between two different noise level pairs (e.g. $\sigma_1$ and $\sigma_2$ vs $\sigma_2$ and $\sigma_3$) permits identifying $\mathcal{C}_{dec}$ and $\mathcal{D}_{dec}$ up to error $O(t)$ each. This, in turn, permits identifying the full model as long as it is not in $\Omega$. This is because two models have identical $F, \mathcal{C}_{dec}, \mathcal{D}_{dec}$ if and only they have identical $F, \mathcal{C}, \mathcal{D}$ (see Eq. 61 for $\mathcal{D}$).

It now remains to show (109). We are given the functions

$$f_1(x) = x + at\,\mathcal{C}_{dec}(x) + a^2t^2\mathcal{D}_{dec}(x) + O(t^3), \tag{110}$$

$$f_2(x) = x + bt\,\mathcal{C}_{dec}(x) + b^2t^2\mathcal{D}_{dec}(x) + O(t^3), \tag{111}$$

where $t > 0$ is small. Our goal is to express

$$f_1^{-1}(f_2(\theta))$$

in terms of $\mathcal{C}_{dec}(\theta)$, $\mathcal{D}_{dec}(\theta)$, and derivatives of $\mathcal{C}_{dec}(\theta)$. Assume that the inverse has the expansion

$$x = f_1^{-1}(f_2(\theta)) = \theta + \varepsilon_1 t + \varepsilon_2 t^2 + O(t^3),$$

with unknown coefficients $\varepsilon_1$ and $\varepsilon_2$ to be determined. Since

$$f_1(x) = x + at\,\mathcal{C}_{dec}(x) + a^2t^2\mathcal{D}_{dec}(x) + O(t^3),$$

substitute $x = \theta + \varepsilon_1 t + \varepsilon_2 t^2$:

$$f_1(\theta + \varepsilon_1 t + \varepsilon_2 t^2) = \theta + \varepsilon_1 t + \varepsilon_2 t^2 + at\,\mathcal{C}_{dec}(\theta + \varepsilon_1 t + \varepsilon_2 t^2) + a^2t^2\mathcal{D}_{dec}(\theta + \varepsilon_1 t + \varepsilon_2 t^2) + O(t^3).$$

We now expand $\mathcal{C}_{dec}(\theta + \varepsilon_1 t + \varepsilon_2 t^2)$ using Taylor's theorem:

$$\mathcal{C}_{dec}(\theta + \varepsilon_1 t + \varepsilon_2 t^2) = \mathcal{C}_{dec}(\theta) + \mathcal{C}'_{dec}(\theta)(\varepsilon_1 t + \varepsilon_2 t^2) + \frac{1}{2}\mathcal{C}''_{dec}(\theta)(\varepsilon_1 t)^2 + O(t^3).$$

Retaining terms up to $t^2$, we have

$$\mathcal{C}_{dec}(\theta + \varepsilon_1 t + \varepsilon_2 t^2) = \mathcal{C}_{dec}(\theta) + \varepsilon_1 t\,\mathcal{C}'_{dec}(\theta) + O(t^2).$$

Similarly, for $\mathcal{D}_{dec}(\theta + \varepsilon_1 t + \varepsilon_2 t^2)$ we need only

$$\mathcal{D}_{dec}(\theta + \varepsilon_1 t + \varepsilon_2 t^2) = \mathcal{D}_{dec}(\theta) + O(t).$$

Substituting these into the expansion for $f_1(x)$ yields:

$$\begin{aligned} f_1(\theta + \varepsilon_1 t + \varepsilon_2 t^2) &= \theta + \varepsilon_1 t + \varepsilon_2 t^2 + at\Big(\mathcal{C}_{dec}(\theta) + \varepsilon_1 t\,\mathcal{C}'_{dec}(\theta)\Big) + a^2t^2\mathcal{D}_{dec}(\theta) + O(t^3) \\ &= \theta + \varepsilon_1 t + \varepsilon_2 t^2 + at\,\mathcal{C}_{dec}(\theta) + a\varepsilon_1 t^2\,\mathcal{C}'_{dec}(\theta) + a^2t^2\mathcal{D}_{dec}(\theta) + O(t^3). \end{aligned}$$

Simultaneously, from (111), we have

$$f_2(\theta) = \theta + bt\,\mathcal{C}_{dec}(\theta) + b^2t^2\mathcal{D}_{dec}(\theta) + O(t^3).$$

Since $f_1(x) = f_2(\theta)$, we equate coefficients of corresponding powers of $t$. First, at order $t^0$, we have $\theta = \theta$. Second, at order $t^1$:

$$\varepsilon_1 + a\,\mathcal{C}_{dec}(\theta) = b\,\mathcal{C}_{dec}(\theta).$$

Thus, we obtain:

$$\varepsilon_1 = (b-a)\mathcal{C}_{dec}(\theta).$$

Third, at order $t^2$:

$$\varepsilon_2 + a\varepsilon_1\,\mathcal{C}'_{dec}(\theta) + a^2\mathcal{D}_{dec}(\theta) = b^2\mathcal{D}_{dec}(\theta).$$

Substitute $\varepsilon_1 = (b-a)\mathcal{C}_{dec}(\theta)$ to get:

$$\varepsilon_2 + a(b-a)\mathcal{C}_{dec}(\theta)\mathcal{C}'_{dec}(\theta) + a^2\mathcal{D}_{dec}(\theta) = b^2\mathcal{D}_{dec}(\theta).$$

Thus,

$$\varepsilon_2 = b^2\mathcal{D}_{dec}(\theta) - a^2\mathcal{D}_{dec}(\theta) - a(b-a)\mathcal{C}_{dec}(\theta)\mathcal{C}'_{dec}(\theta),$$

or equivalently,

$$\varepsilon_2 = (b^2-a^2)\mathcal{D}_{dec}(\theta) - a(b-a)\mathcal{C}_{dec}(\theta)\mathcal{C}'_{dec}(\theta).$$

Substituting the values of $\varepsilon_1$ and $\varepsilon_2$ into the expansion for $x$ gives:

$$f_1^{-1}(f_2(\theta)) = \theta + (b-a)t\,\mathcal{C}_{dec}(\theta) + t^2\Big[(b^2-a^2)\mathcal{D}_{dec}(\theta) - a(b-a)\mathcal{C}_{dec}(\theta)\mathcal{C}'_{dec}(\theta)\Big] + O(t^3).$$

□

##### S5.1.2 Simulation

**Implementation** We focus on circular stimulus spaces. We start from the likelihoods $P(m_1|\theta_1)$, $P(m_2|\theta_2)$, and the derived estimates $\widehat{\theta}_1$, $\widehat{\theta}_2$. The generative model for a response stating that "$\theta_1 > \theta_2$" has the likelihood:

$$P(response = 1|\theta_1, \theta_2) = \int_{X\times X} \widehat{P}(\widehat{\theta}_1 = s|\theta_1)\widehat{P}(\widehat{\theta}_2 = t|\theta_2)1_{\sin(s-t)>0}d(s,t)$$

Recall that the implementation of Hahn and Wei [5] discretizes the stimulus space via a regularly-spaced grid $x_1, \dots, x_N$; it is thus necessary to define $\widehat{P}(\widehat{\theta}_1 = x_s|\theta_1)$ for points $x_s$ on the grid. We define von Mises kernels ($k = 1,2; i = 1,\dots,N$) and normalize these:

$$\phi_{i,k}(j) = \frac{\exp\Big(\gamma\cdot\cos\Big(\widehat{\theta}_{k,i} - x_j\Big)\Big)}{\sum_s \exp\Big(\gamma\cdot\cos\Big(\widehat{\theta}_{k,i} - x_s\Big)\Big)} \tag{112}$$

where we choose $\gamma = 500$ to make the kernels very narrow, and use these to project the distributions $P(\widehat{\theta}_k|\theta_k)$ onto the grid:

$$\widehat{P}(\widehat{\theta}_k = x_j|\theta_k) \propto \sum_{s=1}^{N} \phi_{s,k}(j)P(m_k = x_s|\theta_k) \tag{113}$$

We then define

$$P(response = 1|\theta_1, \theta_2) = \sum_{s,t:\sin(x_s-x_t)>0} \widehat{P}(\widehat{\theta}_1 = x_s|\theta_1)\widehat{P}(\widehat{\theta}_2 = x_t|\theta_2) + \frac{1}{2}\sum_s \widehat{P}(\widehat{\theta}_1 = x_s|\theta_1)\widehat{P}(\widehat{\theta}_2 = x_s|\theta_2)$$

For simplicity, our simulations assume a zero lapse rate.

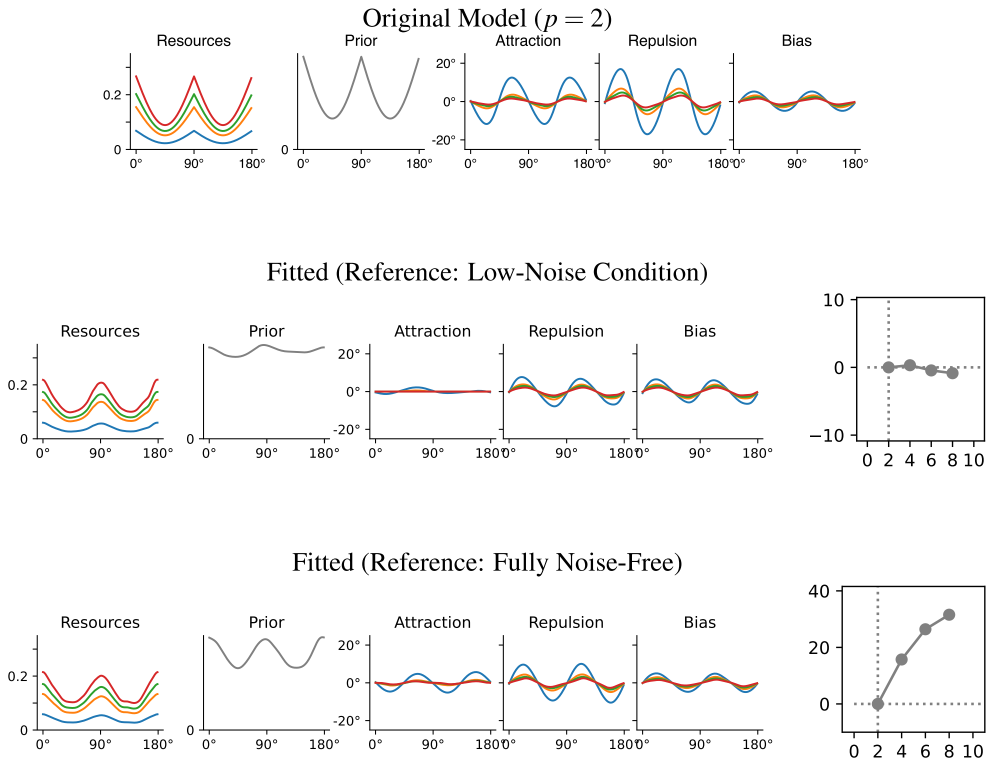

Figure S29: Results from simulated 2AFC paradigm, at N=40K trials (Section S5.1.2). We considered the cases where the reference either has the same noise level as the least-noisy test condition or is noise-free. Fits were only obtained at $p \geq 2$ due to numerical challenges at lower $p$. The encoding is generally recovered well; the prior and the loss function are recovered more accurately when the reference is noise-free.

**Experiment** We generate uniformly distributed $\theta_1$, and then choose $\theta_2$ uniformly at a distance of $\leq 40^\circ$ when viewing the full sensory space as a full $360^\circ$ circle ($\cos(\theta_1 - \theta_2) \gtrapprox 0.76$). We note that, in principle, staircase procedures could be used to optimize the selection of stimuli in an experiment. However, such an adaptive procedure is substantially more complex to implement than the simple estimation paradigm. We compared two setups: one where the reference is provided at the same noise as the lowest-noise test condition, and one where the the reference is encoded without any noise. Results show that, while the encoding is fitted in both cases, only the latter setup permits identifying prior and loss function (Figure S29). Overall, this result suggests 2AFC can be used in principle to identify models, as it theoretically provides the same information as the estimation paradigm. However, at a finite number of trials, at least without adaptive choice of stimuli, the simple estimation procedure is likely to be more efficient.

#### S5.2 Effect of Varying Stimulus Noise

Here, we discuss the extent to which models are identifiable when one varies not sensory noise, but rather stimulus (external) noise. The model with both sensory and stimulus noise takes the form

$$m = F(\theta + \varepsilon) + \delta \tag{114}$$

where $\varepsilon \sim \mathcal{N}(0, \tau^2)$, where $\tau^2$ here denotes the magnitude of stimulus noise. Whereas sensory noise reflects unavoidable noise of the imperfect neural encoding, stimulus noise is applied to the stimulus. For example, multiple Gabor patches or color patches ma be presented in an array, each with some noise applied to the stimulus value (e.g.,

[23, 4, 13]). Hahn and Wei [5] (Theorem 2) proved that the bias (for positive even $p$) is given as

$$\underbrace{\left(\frac{1}{\mathcal{J}(\theta)}+\tau^2\right)(\log p_{prior}(\theta))'}_{Prior\ Attraction}+\underbrace{\left[1+\frac{p-2}{4}\frac{1}{1+\tau^2\mathcal{J}(\theta)}\right]\left(\frac{1}{\mathcal{J}(\theta)}\right)'}_{Likelihood\ Repulsion}+O\left((\sigma^4+\tau^4+\sigma^2\tau^2)\right) \tag{115}$$

where $\mathcal{J}$ is the Fisher Information of the sensory encoding, extending the decomposition reviewed in Section S2.2. That is, stimulus noise increases attraction, and induces (for $p > 2$) a loss-function-dependent reduction of the repulsion. While this result is proven for positive even exponents ($p = 2, 4, 6, \dots$), simulations Hahn and Wei [5, SI Appendix, Figure S5] show that similar behavior holds at other exponents. Furthermore, the variability has the form

$$\tau^2+\frac{1}{\mathcal{J}(\theta)}+O\left((\sigma^4+\tau^4+\sigma^2\tau^2)\right) \tag{116}$$

Based on this, in principle, one can identify models by:

1. Comparing the variability at two levels of stimulus noise to recover $\frac{1}{\mathcal{J}(\theta)}$ up to higher-order error.
2. Then, in the expression for the bias, all except for the green terms are known up to higher-order error:

$$\underbrace{\left(\frac{1}{\mathcal{J}(\theta)}+\tau^2\right)(\log p_{prior}(\theta))'}_{Prior\ Attraction}+\underbrace{\left[1+\frac{p-2}{4}\frac{1}{1+\tau^2\mathcal{J}(\theta)}\right]\left(\frac{1}{\mathcal{J}(\theta)}\right)'}_{Likelihood\ Repulsion}+O\left((\sigma^4+\tau^4+\sigma^2\tau^2)\right) \tag{117}$$

   Comparing the bias at two levels of stimulus noise permits identifying these unknowns.

This applies to *all* models, even those in the exceptional set $\Omega$ of our Theorem 2. Thus, with unbounded amounts of behavioral data, all models are identifiable in principle.

An interesting question is whether varying stimulus noise or sensory noise might be more effective in ensuring identifiability. Stimulus noise increases the variance:

$$\tau^2+\frac{1}{\mathcal{J}(\theta)}+O(\sigma^4) \tag{118}$$

and also increases the prior attraction term. In contrast, importantly, when $p > 2$, stimulus noise *reduces* the loss-function-dependent repulsion, as shown by Equation 115: the higher $\tau^2$ is, the more the likelihood repulsion approaches a loss-function-independent limit

$$\left(\frac{1}{\mathcal{J}(\theta)}\right)' \tag{119}$$

Overall, thus, stimulus noise broadens the response distribution while simultaneously *reducing* the loss-function-dependent repulsion. This is in contrast to sensory noise, where increasing it leads to an *increase* in the loss-function-dependent repulsion. We thus expect that adding stimulus noise substantially reduces the signal-to-noise ratio compared to the setting where stimulus noise is zero but sensory noise is varied.

In practice, we indeed find that, at the same number of trials, varying stimulus noise might be less effective in achieving identifiability than sensory noise (Figure S30).

#### S5.3 Identifiability when Encoding Varies with Noise Level

**Theoretical Discussion** While our Theorems 3 and 4 apply in the setting where the encoding is shared between noise levels, here we discuss the case where the encoding varies between noise levels. For each noise level $\sigma$ and associated encoding $F'_\sigma$, we have the model $M_\sigma := \langle F'_\sigma, p_{prior}, p\rangle$, where $p$ is the loss function exponent. We first discuss the case without motor noise. Here, Theorem 1 can be used to recover the encoding in each noise level to high accuracy. Theorem 2 can then be used to obtain the prior, conditional on the loss function exponent. By comparing the bias at each noise level $\sigma$ to the expansion from Lemma S23, we can expect to recover the loss function exponent

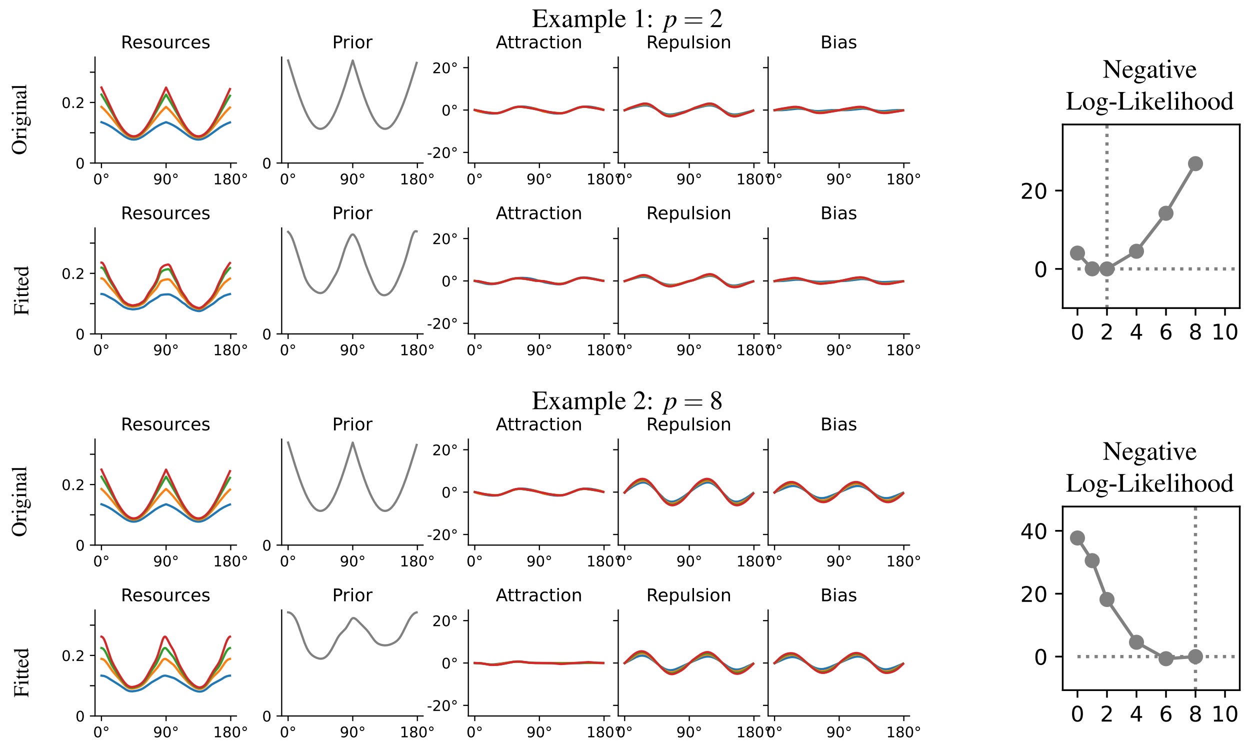

Figure S30: Varying stimulus noise, while keeping sensory noise fixed (40K trials). This can also lead to recoverability, as theoretically demonstrated in Section S5.2. Comparison to results varying sensory noise (Figures S6 and S10) suggests that varying stimulus noise results in a weaker signal then when varying sensory noise; also, the prior at $p = 8$ is recovered less successfully then when sensory noise is varied. Overall, varying stimulus noise can support identifiability, but varying sensory noise is more advisable when feasible.

$p$ for most models outside of a small exceptional set, similar to the set $\Omega$ of Theorem 3. The situation is more complex in the presence of motor noise. A general approach is to deconvolve the response distribution with a motor variance $\tau^2$, and then apply the same reasoning as above. While $\tau^2$ is unknown, one may expect that only one possible $\tau^2$ may lead to a consistent result, at least on models outside of a small exceptional set. Thus, we expect that the theoretical guarantees of Theorems 3 and 4 extend to many models where the encoding varies between noise levels.

**Simulations** Here, we evaluate the identifiability of models on the situation where the shape of the encoding may itself depend on the noise level. We considered a situation where the encodings are similar across noise levels, but vary in their steepness (Figure S31), and where they are independent (Figure S32).

### S6 Applications to Experimental Data

For the orientation perception data, we show fits at one level or at five levels of noise at 1K trials in Figure S33.

For the time interval perception data, we show model fit statistics when either presupposing a Weber's law-based encoding (as in the main paper) or a freely fitted encoding (alternative) in Figure S34; even in the latter case, data generated from a bimodal prior enables identifiability.

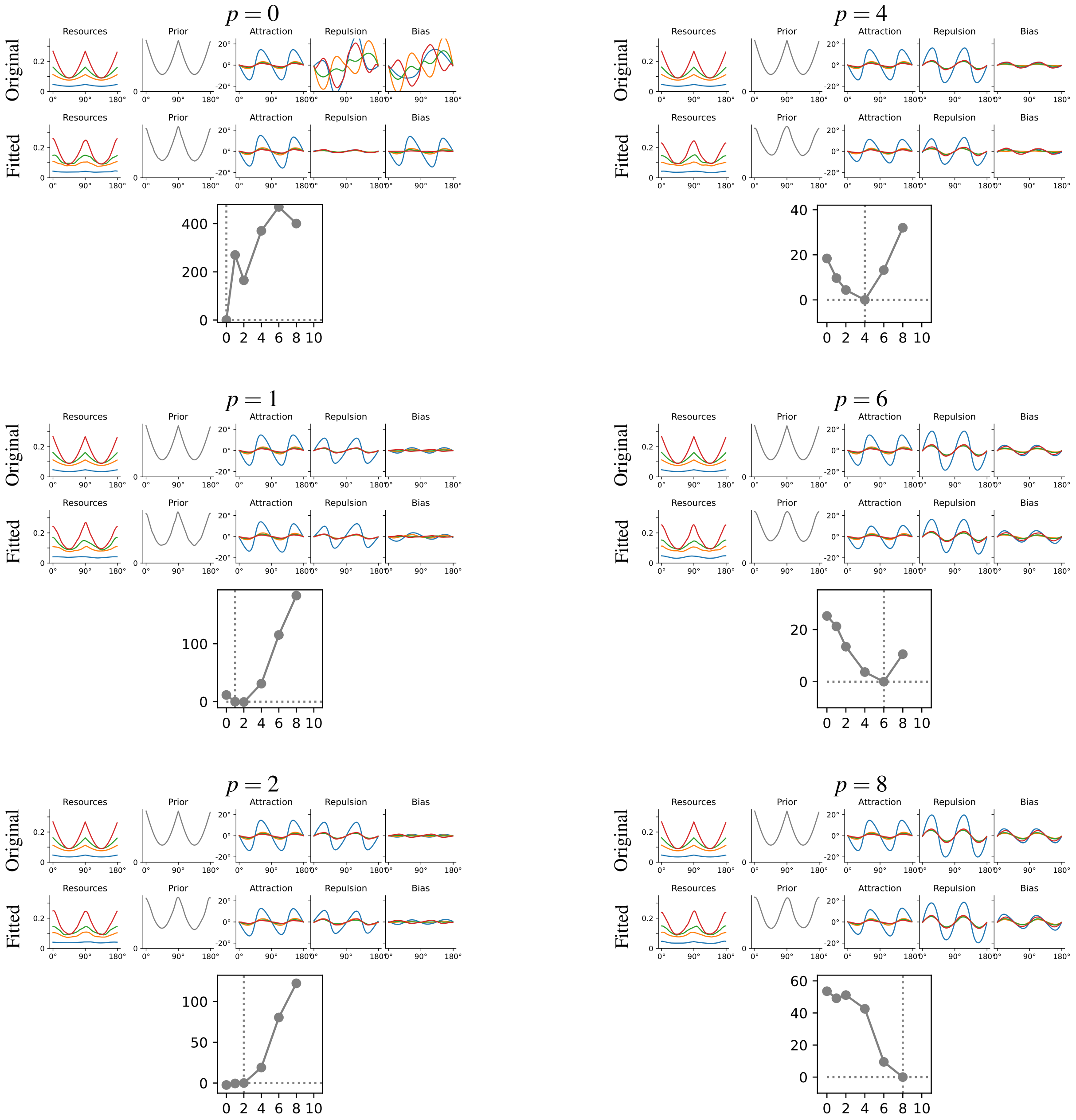

Figure S31: Models may be identifiable even when the encoding varies between noise conditions. Here, we consider a model where the encoding resources, at sensory noise $\sigma_i$ ($i = 1,2,3,4$), are $F'(\theta) \propto 1 - \alpha_i \sin(\theta)$. In particular, the encoding has a similar overall shape across noise conditions but is flatter when noise is higher; this might be a reasonable assumption, for instance, when the noise is due to decreased contrast. Nonetheless, the ground-truth loss function remains well-identified.

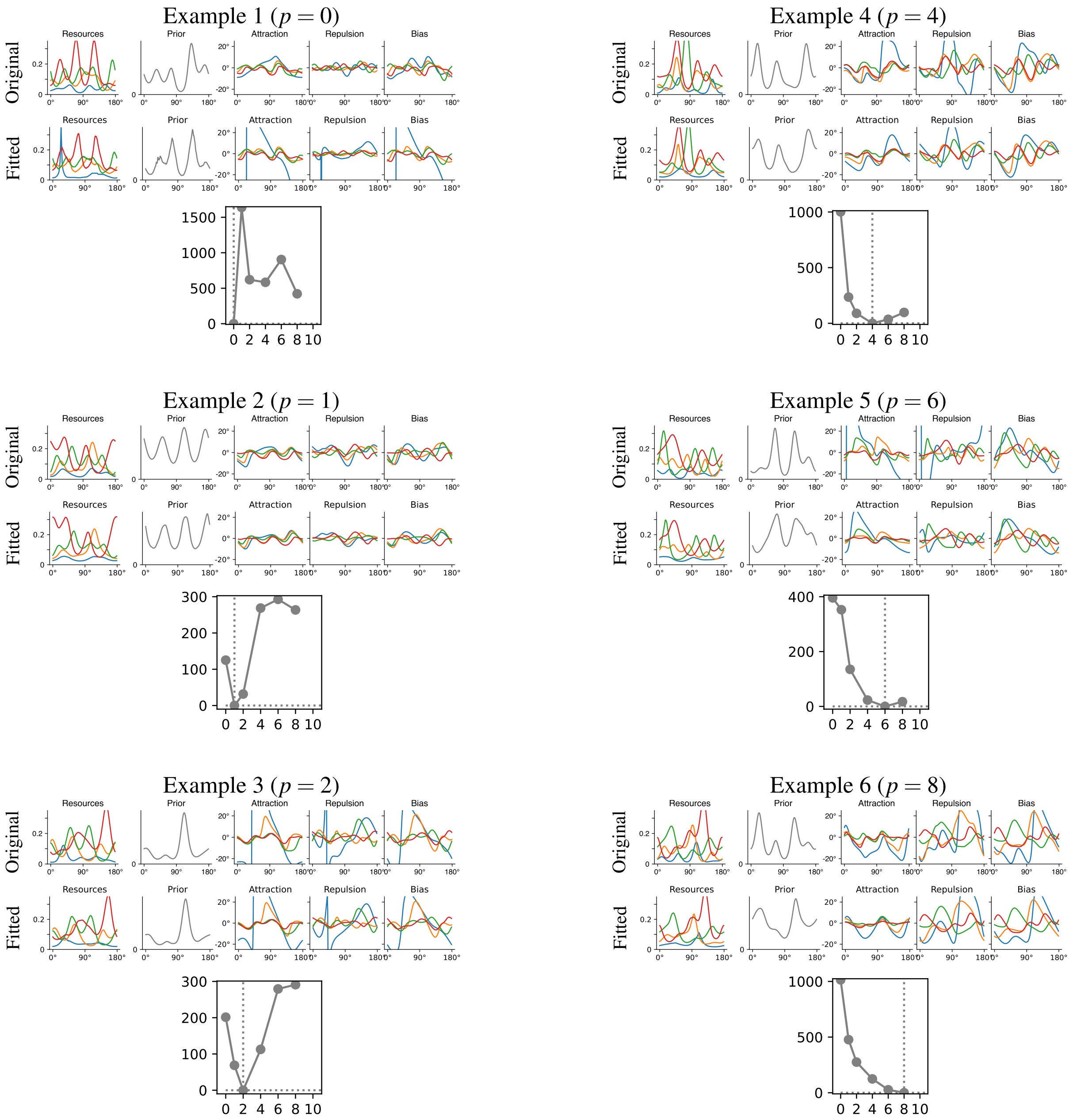

Figure S32: Models may be identifiable even when the encoding varies between noise conditions. Here, the encoding resources are independently randomly generated in each noise condition, while the prior is shared across conditions. Even in this situation, the ground-truth loss function can be identified.

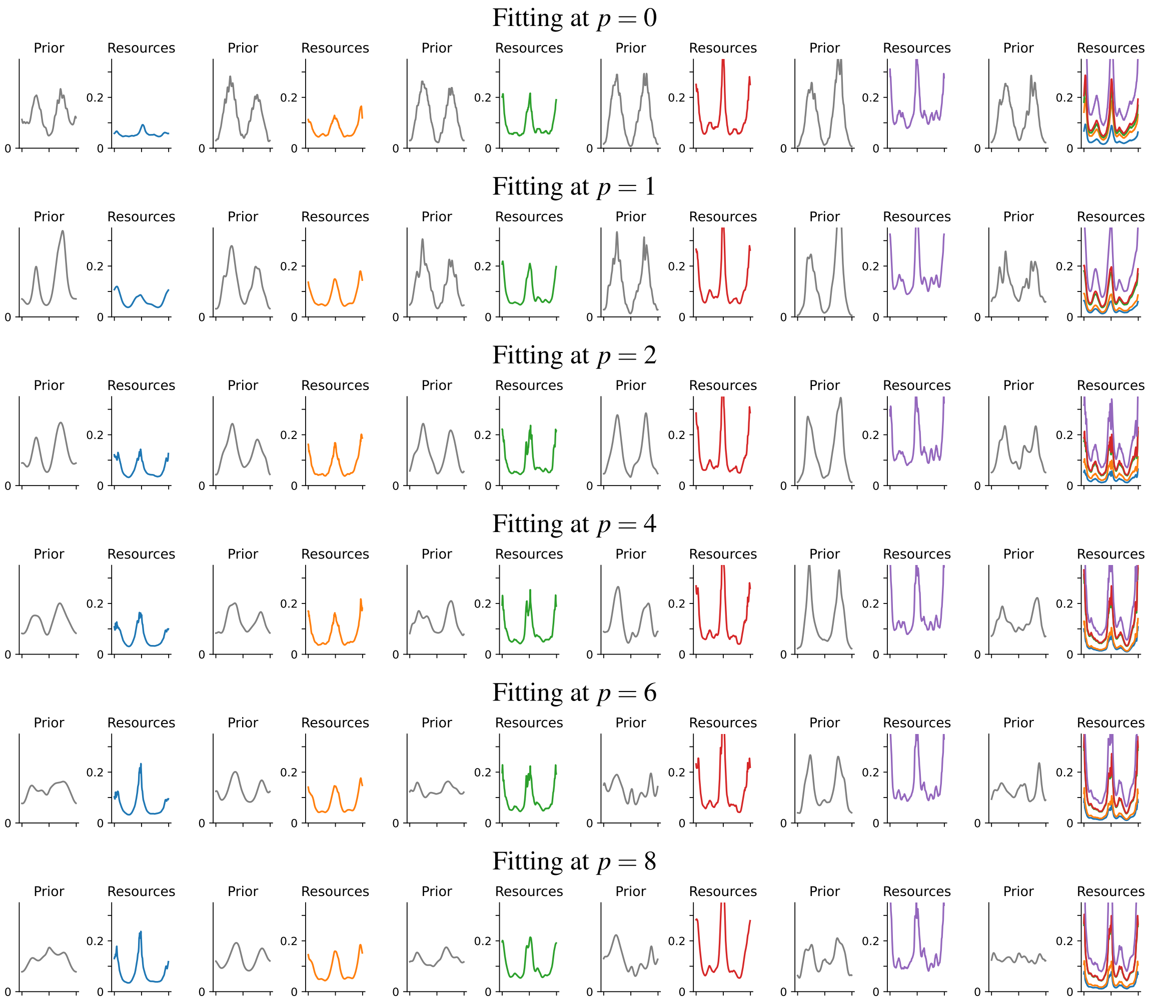

Figure S33: Orientation perception (data collected by [1]). Fitting to 1K trials from the dataset of de Gardelle et al at one level of noise (left=large noise, right=small noise; first five column) or at all five levels (rightmost column). In each case, we show the prior (left) and the encoding (right). Coloring of noise levels is as in Main Paper, Figures 6–7. When fitting at a single level of noise, the encoding is fitted consistently even across loss functions, in accordance with the theory. However, the prior is fitted inconsistently: fitting with low exponents consistently leads to a prior peaking at oblique directions; fitting with higher exponents leads to a variety of results. As model fit is very similar across loss function exponents at one level of noise, for each level of sensory noise (Main Paper, Figure 7), loss function and prior are not identifiable. In contrast, when combining noise levels, model fit vary between loss functions (Main Paper, Figure 7). At high exponents, an approximately uniform prior is fitted. Compare Main Paper, Figure 7 for the fit at $p = 8$ on the full dataset (9,936 trials), which is highly consistent with the fit obtained at 1K trials but five noise levels.

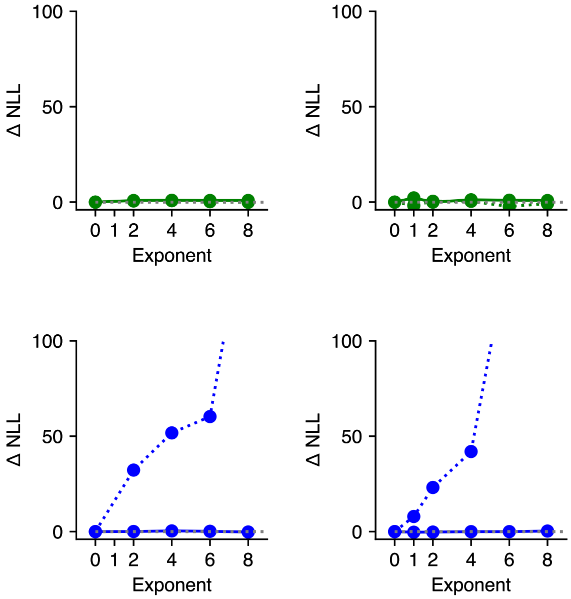

Figure S34: Model fit statistics for simulated datasets based on the data collected by Remington et al. [19], when presuming an encoding consistent with Weber's law (left, same as Main Paper, Figure 8i) or with a freely fitted encoding (right). Coding of color and line type is as in Main Paper Figure 8i. Even when the encoding is freely fitted, the bimodal prior (dotted, blue) allows identification of the loss function when data at multiple levels of noise is available.

### References

[1] Vincent de Gardelle, Sid Kouider, and Jérôme Sackur. An oblique illusion modulated by visibility: non-monotonic sensory integration in orientation processing. *Journal of vision*, 10 10:6, 2010.

[2] Matthias Fritsche, Eelke Spaak, and Floris P. de Lange. A bayesian and efficient observer model explains concurrent attractive and repulsive history biases in visual perception. *eLife*, 9:e55389, 2020.

[3] Nikos Gekas, Matthew Chalk, Aaron R. Seitz, and Peggy Seriès. Complexity and specificity of experimentally induced expectations in motion perception. *Journal of Vision*, 14:P355 – P355, 2013.

[4] Ahna Reza Girshick, Michael S. Landy, and Eero P. Simoncelli. Cardinal rules: Visual orientation perception reflects knowledge of environmental statistics. *Nature neuroscience*, 14:926 – 932, 2011.

[5] Michael Hahn and Xue-Xin Wei. A unifying theory explains seemingly contradictory biases in perceptual estimation. *Nature Neuroscience*, 27(4):793–804, 2024.

[6] Mehrdad Jazayeri and Michael N. Shadlen. Temporal context calibrates interval timing. *Nature neuroscience*, 13:1020–1026, 2010.

[7] Kari Karhunen. Über lineare methoden in der wahrscheinlichkeitsrechnung. *Ann Acad Sci Fennicae*, 37:1, 1947.

[8] Samuel Karlin and Herman Rubin. The theory of decision procedures for distributions with monotone likelihood ratio. *The Annals of Mathematical Statistics*, pages 272–299, 1956.

[9] EL Lehmann. Ordered families of distributions. *The Annals of Mathematical Statistics*, pages 399–419, 1955.

[10] Michel Loeve. Functions aleatoires du second ordre. *Processus stochastique et mouvement Brownien*, pages 366–420, 1948.

[11] Paul Milgrom and Chris Shannon. Monotone comparative statics. *Econometrica: Journal of the Econometric Society*, pages 157–180, 1994.

[12] Michael J. Morais and Jonathan W. Pillow. Power-law efficient neural codes provide general link between perceptual bias and discriminability. In *Advances in Neural Information Processing Systems*, 2018.

[13] Maria Olkkonen, Patrice McCarthy, and Sarah R. Allred. The central tendency bias in color perception: effects of internal and external noise. *Journal of vision*, 14 11, 2014.

[14] B Sango Otieno and Christine M Anderson-Cook. A more efficient way of obtaining a unique median estimate for circular data. *Journal of Modern Applied Statistical Methods*, 2:168–176, 2003.

[15] Athanasios Papoulis. *Random variables and stochastic processes*. McGraw Hill, 1965.

[16] Frederike H. Petzschner and Stefan Glasauer. Iterative bayesian estimation as an explanation for range and regression effects: A study on human path integration. *The Journal of Neuroscience*, 31:17220 – 17229, 2011.

[17] Rafael Polanía, Michael Woodford, and Christian C Ruff. Efficient coding of subjective value. *Nature neuroscience*, 22(1):134–142, 2019.

[18] Arthur Prat-Carrabin and M. Woodford. Bias and variance of the bayesian-mean decoder. In *Advances in Neural Information Processing Systems*, 2021.

[19] Evan D. Remington, Tiffany V Parks, and Mehrdad Jazayeri. Late bayesian inference in mental transformations. *Nature Communications*, 9, 2018.

[20] Moshe Shaked and J George Shanthikumar. *Stochastic orders*. Springer, 2007.

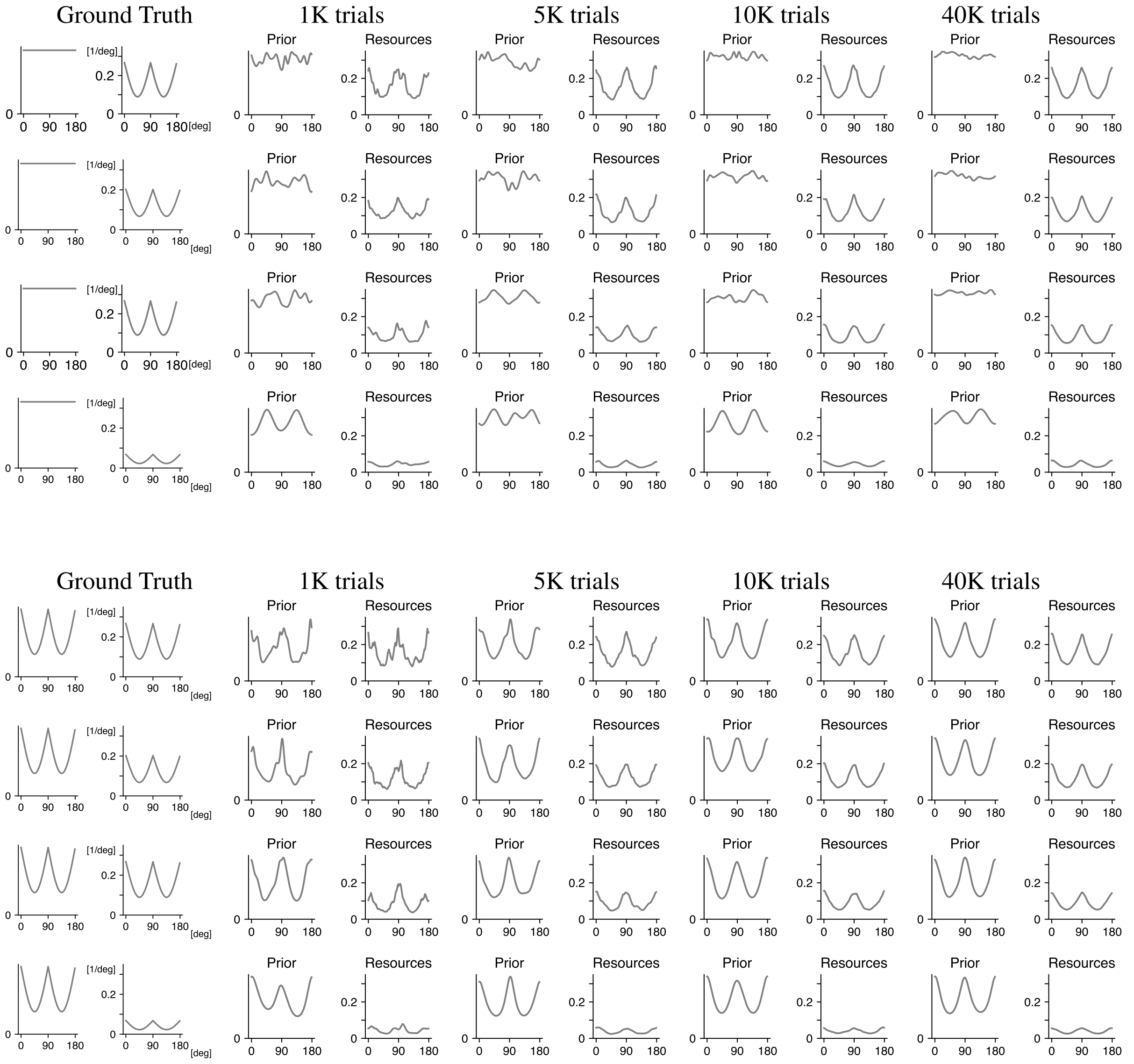

Figure S35: Supplement to Figure 2: We show two combinations of prior (left) and encoding (right), at four noise levels (rows, from low to high noise), by number of trials (columns). See Figure S36 for continuation with two more models.

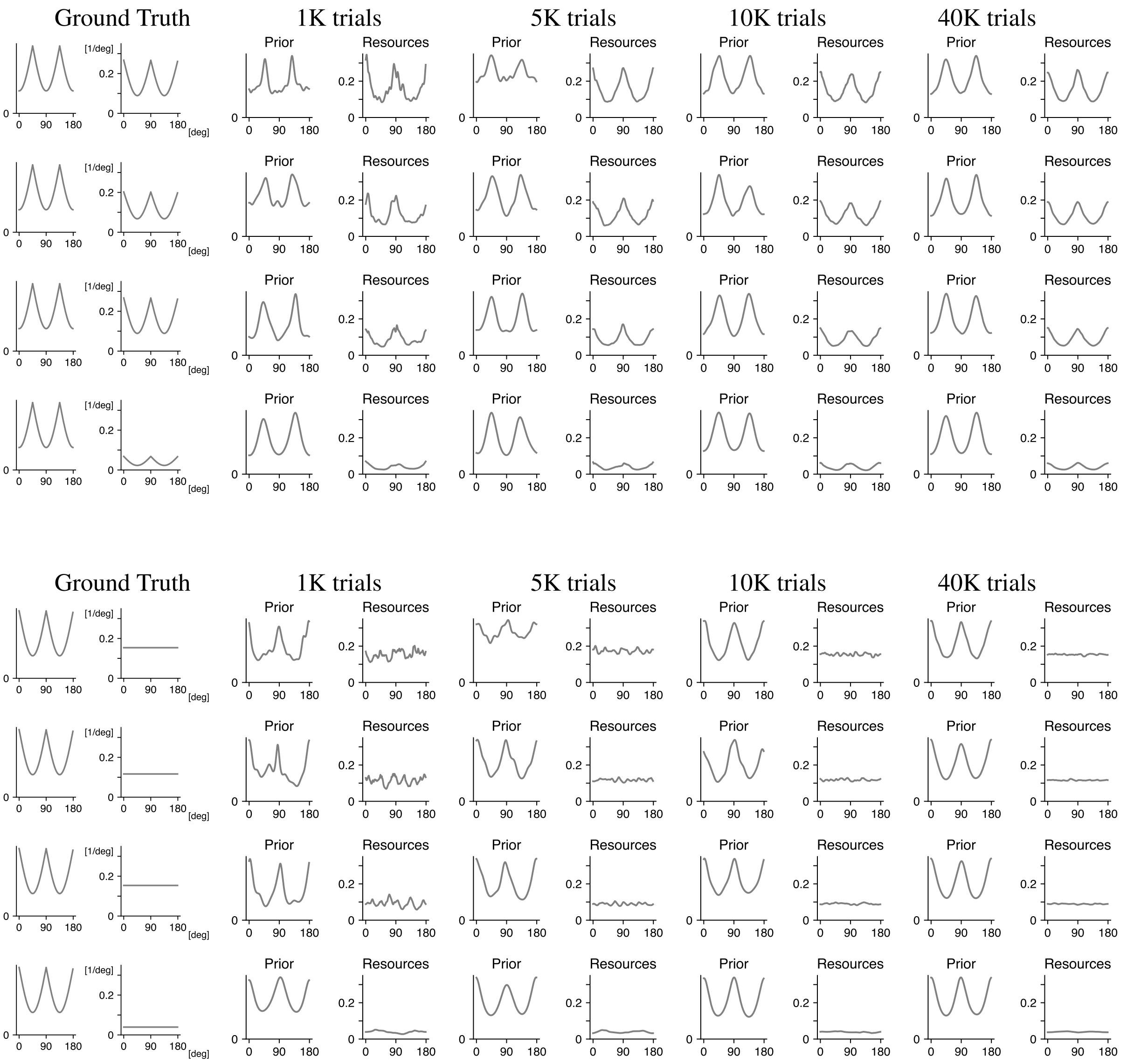

Figure S36: Supplement to Figure 2. We show two combinations of prior (left) and encoding (right), at four noise levels (rows, from low to high noise), by number of trials (columns).

[21] Alan A Stocker and Eero Simoncelli. Sensory adaptation within a bayesian framework for perception. *Advances in neural information processing systems*, 18, 2005.

[22] Alan A. Stocker and Eero P. Simoncelli. Noise characteristics and prior expectations in human visual speed perception. *Nature Neuroscience*, 9:578–585, 2006.

[23] Alessandro Tomassini, Michael J. Morgan, and Joshua A. Solomon. Orientation uncertainty reduces perceived obliquity. *Vision Research*, 50:541–547, 2010.

[24] Donald M Topkis. Minimizing a submodular function on a lattice. *Operations research*, 26(2):305–321, 1978.

[25] Donald M Topkis. *Supermodularity and complementarity*. Princeton university press, 1998.

[26] Xue-Xin Wei and A. Stocker. A bayesian observer model constrained by efficient coding can explain 'anti-bayesian' percepts. *Nature Neuroscience*, 18:1509–1517, 2015.

[27] Xue-Xin Wei and A. Stocker. Lawful relation between perceptual bias and discriminability. *Proceedings of the National Academy of Sciences*, 114:10244–10249, 2017.

[28] Xue-Xin Wei and Alan A Stocker. Efficient coding provides a direct link between prior and likelihood in perceptual bayesian inference. *Advances in neural information processing systems*, 25, 2012.

[29] Hang Zhang, Xiangjuan Ren, and Laurence T. Maloney. The bounded rationality of probability distortion. *Proceedings of the National Academy of Sciences of the United States of America*, 117:22024 – 22034, 2020.
